# Molecular machines measure the nanomechanics of biological membranes

**DOI:** 10.64898/2026.11.02.763668

**Authors:** Alexia Rottensteiner, Miguel Paez Perez, Sioned F. Jones, Mohamad Zandieh, Juliette L. Ciccone, Yongzheng Xing, Ulrike Eggert, Stefan Howorka

**Author notes:** Equally contributing authors (Alexia Rottensteiner, Miguel Paez Perez, Sioned F. Jones).

## Abstract

Mechanical properties such as elastic resistance to distortion define many materials and can routinely be measured with an indenting tip as simple as one’s finger. We have miniaturized the indentation principle by 8 orders of magnitude to analyze lipid bilayer nanomechanics, which are of relevance in biology and disease progression. Our nanoscale indenting tip is designed with DNA nanotechnology to be an integral part of a molecular machine which autonomously binds to membranes, actively indents localized membrane patches, measures the membranes’ elastic response to indentation, and converts the indentation degree into an optical readout signal. Compared to serially operated macro-to-microscale indentation probes, our miniaturization strategy enables deploying millions of the indenting machines for parallel analysis of membrane elasticity. The nanoscale molecular machines are also taken up by biological cells to reveal softer inner membrane compartments compared to stiffer cytoplasmic membranes, using indentation depths of solely up to 1 nm. We envision the DNA-based nanotools to contribute to the analysis of membrane mechanics of cells as well as soft polymeric interphases.

---

The mechanical properties of soft and deformable interphases are of crucial importance for polymeric materials and for biological membranes, where they influence cell function^1–4^ as well as the development of cancer^5–9^, malaria^10,11^, and sickle cell anaemia^11–13^. A key mechanical property is the elastic resistance to distortion, which is quantified by Young’s modulus^14–17^ to reflect lipid packing and membrane organization^18–22^. Membrane elasticity can be measured by indenting the bilayer with cantilever tips of an atomic force microscope^23–25^, membrane-linked beads controlled by optical tweezers^26–28^, and related micropipette aspiration^29,30^, among others^31,32^. As these classical serial techniques use macro-to-microscale scale actuators and external controllers, they are of lower throughput and cannot directly reach cell-internal membranes^33^. To overcome these limitations, we wondered whether indentation could be miniaturized with nanosized cantilever-type tips within functionally self-contained molecular devices. Small size would allow millions of devices to be deployed for parallel analysis of membrane elasticity as well as for cellular uptake to probe internal organelle membranes. The molecular devices could aid understanding biological membrane mechanics, and constitute a nanoengineering feat as four functions would have to be integrated. (i) To specifically locate the nanodevices at bilayer membranes, (ii) to actively indent a nanoscale membrane region, (iii) to measure the membrane’s elastic response to indentation, and (iv) to report on the indentation degree by conversion to a simple signal for readout of elasticity.

Here we describe a multifunctional Molecular Machine (MM) that indents bilayer membranes to measure their local elasticity in a parallel fashion and inside cells (Fig. 1). To design the device, we chose bottom-up DNA nanotechnology, which yields nanometre dimensions, precise nanomechanical movement and controllable forces^34–42^, with defined interactions with lipid bilayer membranes^43–46^, as successfully demonstrated with other nanodevices that puncture^47–52^, shape^53–59^ and fuse bilayers^60–63^, or serve as artificial cytoskeletons^53,54,64^. Our DNA nanodevice features, as a first component, a box-like base (Fig. 1a, Membrane binding of Molecular Machine) which carries cholesterol lipid anchors for membrane localization of MM, to fulfil function (i). The base measures 20 x 12 x 15 nm and is composed of 30 interconnected duplexes. The design of MM relies on the DNA origami approach (Supplementary Figs. 1 and 2), where a scaffold strand (Supplementary Table 1) and complementary, shorter staple strands (Supplementary Table 2) anneal into the DNA nanostructure^34,35^. The sequences of the staple strands were computed with DNA origami design software caDNAno^65^.

**Fig. 1.**
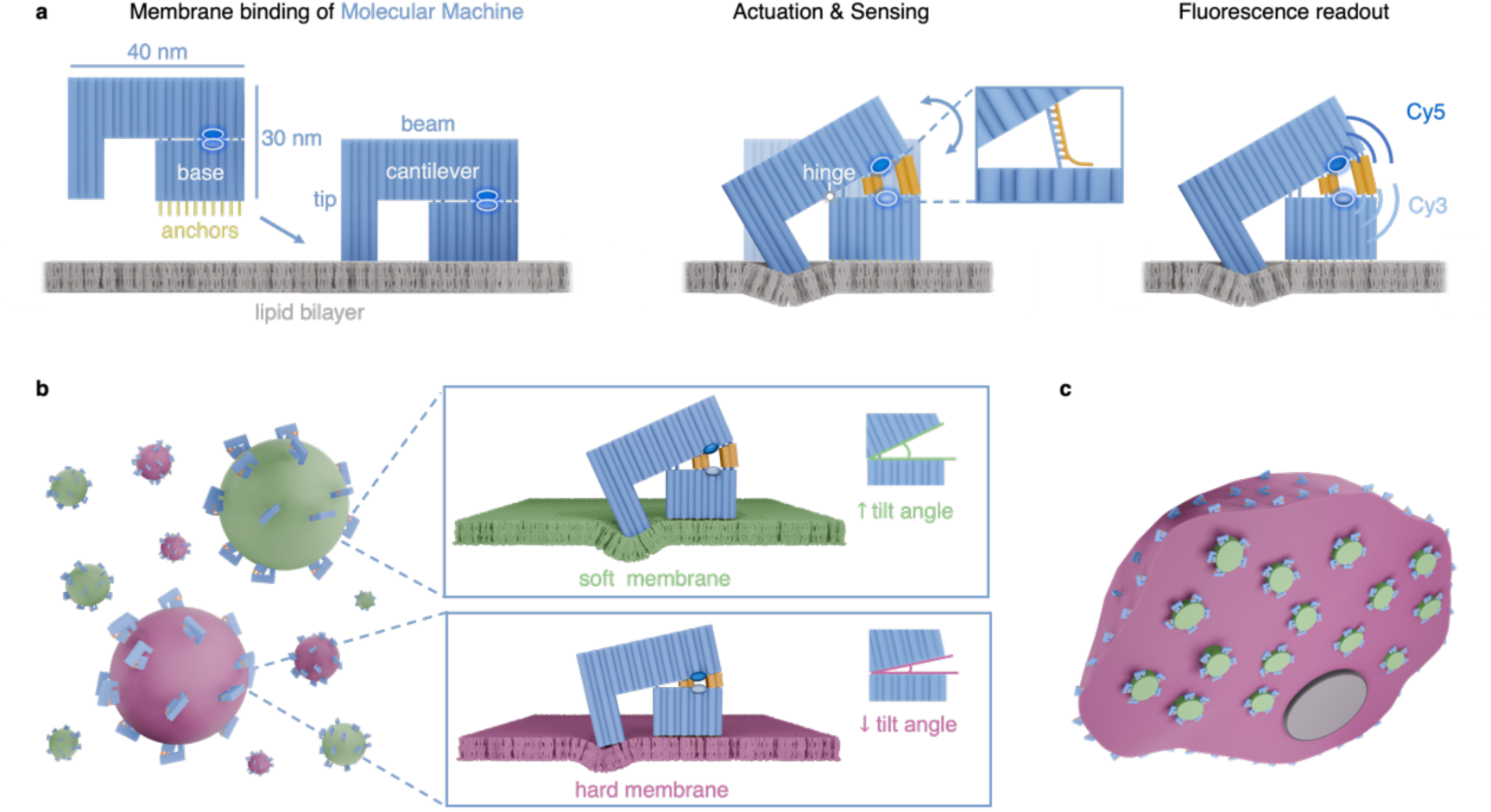
**Molecular Machines (MMs) measure lipid bilayer nanomechanics in a parallel manner and inside cells**. **a**, MM is composed of bundled DNA duplexes represented as blue cylinders. MM’s base binds via cholesterol anchors (yellow) to bilayer membranes. Hybridization of opening oligonucleotides (orange, one shown in inset) to single-stranded MM segments (blue, inset) generates duplexes (orange cylinders) which actuate the MM cantilever. Actuation pushes the MM tip downward and locally indents the membrane. The elastic membrane deformation is measured via a fluorescent FRET readout of incorporated Cy3 (light blue) and Cy5 (dark blue). **b**, MM indentation allows parallel quantification of local membrane elasticities, whereas the opening tilt angle is larger for softer (green) membranes than for stiffer membranes (magenta). Orange cylinders of different lengths indicate duplexes where corresponding opening strands of corresponding length are hybridized (hybridization step not shown). **c**, In cells, MM measures the elasticity of the plasma membrane (magenta) as well as internal membranes (green). The cell nucleus is shown in dark grey.

**Fig. 2.**
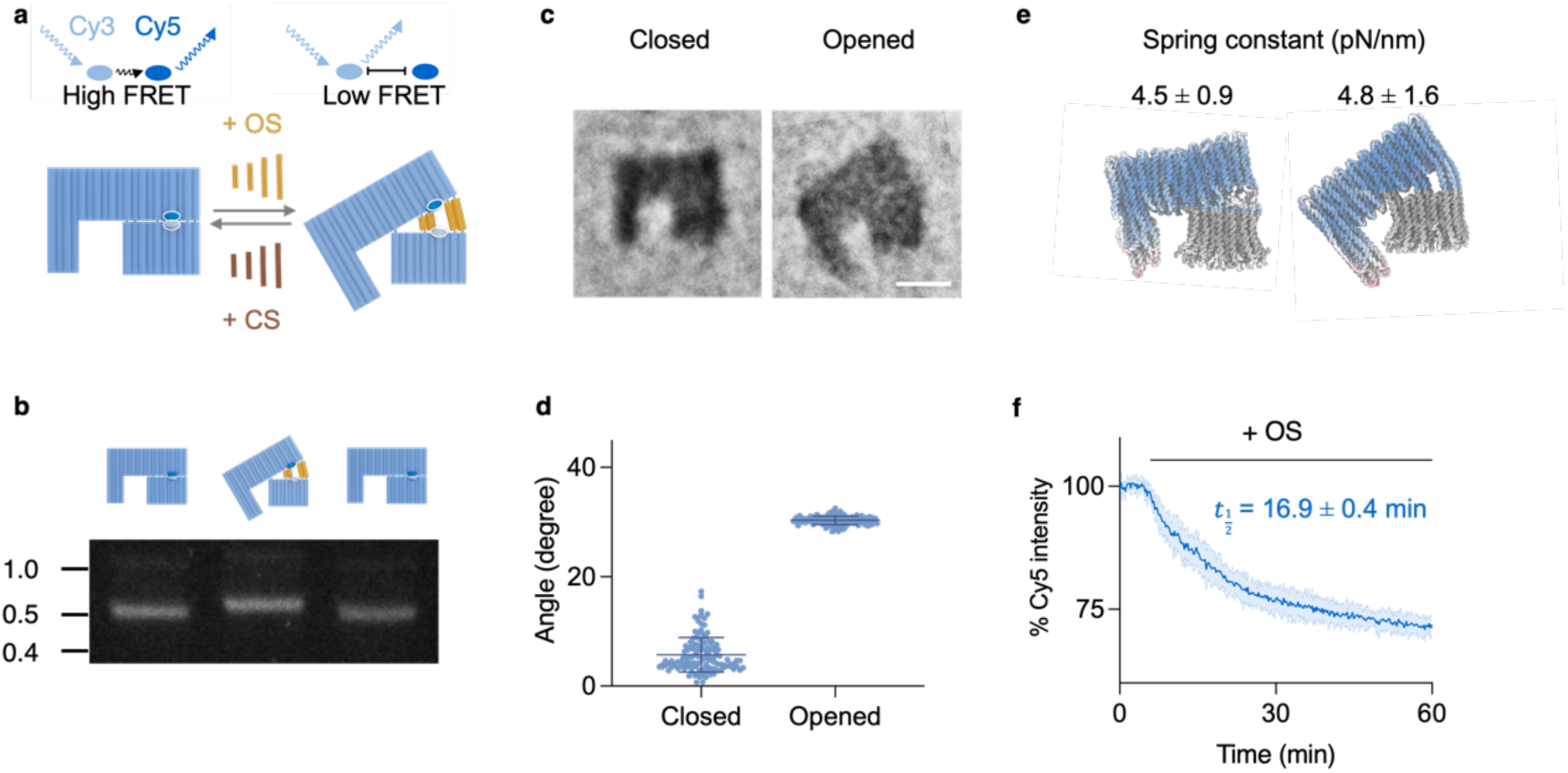
**Actuated opening and closing of MM is tracked with fluorescence readout**. **a**, Reversible actuation of MM by opening and closing strands changes MM’s tilt angle and the associated fluorescence signals of Cy3 and Cy5 distance markers. **b**, MM opening and closing changes MM’s gel electrophoretic migration. **c**, Overlay of stained TEM images of MM in closed and opened state; scale bar 20 nm. **d**, Analysis of MM tilt angles from TEM images (n = 128 and n = 131) for closed and opened state, respectively). **e**, oxDNA simulated structure of MM in closed and opened conformation with the cantilever’s spring constant obtained from the average RMSD of the cantilever (Supplementary Fig. 14). **f**, Cy5 FRET-dependent kinetic emission traces tracking MM opening upon addition of OS at 5 min. Traces were processed via 2^nd^ order smoothing (4 neighbours) integrated within GraphPad Prism. Mean trace ± SD from n = 11.

For active membrane indentation (ii), MM features as a second component, an L-shaped cantilever with a vertical tip (Fig. 1a, Membrane binding of MM, Actuation & Sensing) made of interconnected duplexes (Supplementary Figs. 1 and 2). The cantilever tip is 15 nm high and the horizontal beam is 40 nm wide (Fig. 1a; Supplementary Fig. 2). The beam is connected to the MM base via hinges (Fig. 1a, Actuation & Sensing) made of single-stranded DNA (Supplementary Figs. 2 and 3, Supplementary Table 3) to allow for controlled tilting cantilever movement. MM is kept in a closed, non-indenting state (Fig. 1a, Actuation & Sensing, semitransparent MM) by 12 single-stranded and structurally condensed actuator sequences positioned between the beam and base (Supplementary Figs. 3 and 4, Supplementary Table 4). To actuate MM into the open indenting state, opening strands are added to hybridize to the condensed actuator strands and form extended DNA duplexes (Fig. 1a, Actuation & Sensing; Supplementary Fig. 4, Supplementary Table 5). This transition exerts a force and pushes the right side of the beam up and simultaneously moves the cantilever tip down with a designed maximum tilt angle of 30°. The opening is engineered to be reversible by adding complementary closing strands (Supplementary Table 6), which remove the opening strands by toehold-mediated competitive hybridization.

**Fig. 3.**
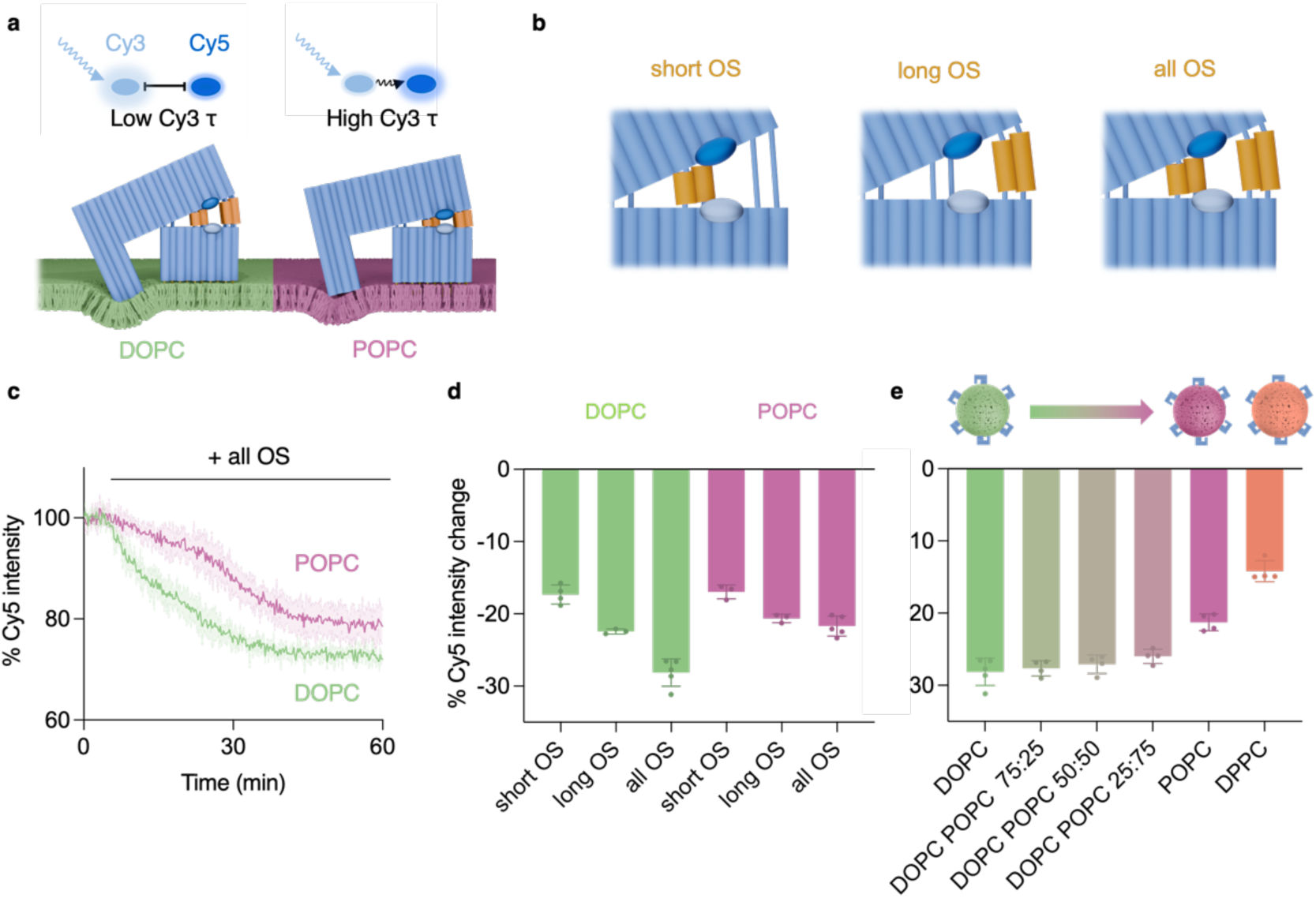
MM acuation indents lipid bilayers and reports on their characteristic elastic deformation using fluorescence readout. **a**, MM indentation of soft DOPC and stiff POPC membranes results in corresponding different tilt angles and Cy5 emission levels caused by FRET. **b,** Side view of MM opened with either six short, six long, or all OS. The resulting different tilt angles are not shown for clarity. **c**, Kinetic Cy5 emission traces for MM bound to DOPC and POPC vesicles upon addition of OS, in 50 mM HEPES 500 mM NaCl. All traced were processed via 2^nd^ order smoothing (4 neighbours) integrated within GraphPad Prism. Mean trace ± SD from n = 5. **d,** Percentage fluorescent Cy5 change upon indentation of DOPC and POPC vesicles by opening MM with short, long, or all OS, after 60 min incubation; mean ± SD from n = 3-5. **e,** Percentage Cy5 emission change for MM-mediated indentation of membrane vesicles composed of DOPC, DOPC:POPC of indicated ratios, POPC and DPPC upon addition of all OS, after 60 min incubation; mean ± SD from n = 5.

**Fig. 4.**
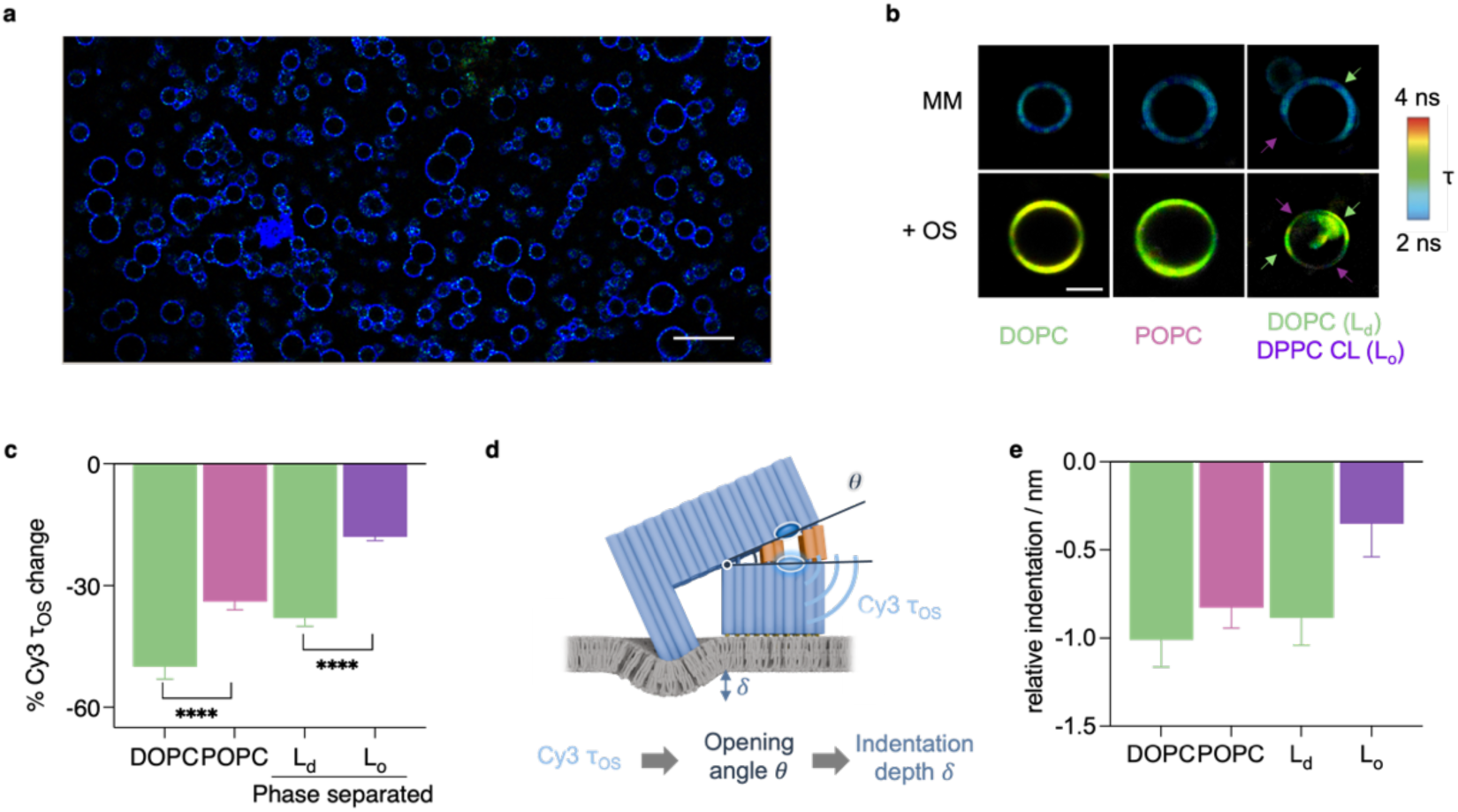
**MM reports on localized elastic indentation of membranes using readout with fluorescence lifetime imaging microscopy (FLIM)**. **a,** Confocal fluorescence microscopic image of DOPC GUVs decorated with closed MM. Scale bar, 50 µm **b**, FastFLIM images of MM-decorated GUVs composed of DOPC, POPC, and phase-separated 40/40/20 DOPC/DPPC/Cholesterol. Green and magenta arrows indicate L_d_ and Lo phases, respectively. The FLIM image pixels are coloured for *τ* (Cy3). Scale bar 20 µm. Microscopic FastFLIM images report the average photon arrival time, while FLIM lifetime values reported as numbers are derived from fitting the time-resolved fluorescence decay trace. **c**, Bar chart on the relative change in Cy3 lifetime following OS addition showing mean ± SD of n = 10; one-way ANOVA with Tukey’s multiple comparison test at ****P <0.0001. The relative change represents 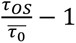. **d**, Scheme illustrating how fitted Cy3 lifetime can be converted via FRET efficiency and MM opening angle θ into indentation depth δ. **e**, Bar chart of MM-mediated indentation depths of indicated membranes as obtained by FLIM readout.

MM actuation pushes the cantilever tip into the adherent membrane (Fig. 1a, Actuation & Sensing). We surmised that the indentation degree depends on the membrane elasticity and associated energetic cost of deformation (function iii) resulting in a bigger tilt angle for a softer membrane and a smaller angle for a stiffer membrane (Fig. 1b). This should allow probing of local elasticity in a parallel fashion (Fig. 1b) and inside cells (Fig. 1c). MM’s design should provide sufficient energy for membrane indentation due to the deliberately chosen number of actuator strands and tip dimensions, using as rational design basis the Hertz’ biophysical model on indenting elastic membranes (Supplementary Note 1). Tip deformation is expected to be small, as its duplexes are predominantly aligned with the indentation axis and resist the load through axial compression.^66^ Furthermore, the footprint size of the MM tip of around 8 x 12 nm is sufficiently large and effects enough membrane lipids to interpret the molecular membrane indentation in terms of classical continuum-based elasticity^67^. To reflect the molecular nanoscale-underpinning of bilayer elasticity, an alternative term, microelasticity, may be used, analogous to microviscosity^68,69^.

To report on membrane indentation and elasticity (iv), we designed MM to convert the tilt angle into a fluorescence signal (Fig. 1a, Fluorescence readout). To measure the angle, the distance-reporting fluorophores Cy3 and Cy5 are positioned at MM’s base and beam, respectively, so that Förster resonance energy transfer (FRET) can track changes in fluorophore distance occurring during MM actuation (Fig. 1a, Fig. 2a, Supplementary Fig. 5). In the design, weak membrane indentation of a stiff membrane would keep Cy3 and Cy5 close, resulting in strong FRET and an enhanced Cy5 signal (Fig. 2a) while a stronger indentation of the soft membrane would lead to weak FRET and a stronger Cy3 signal^70^ according to a quantitative model which also includes associated changes in fluorescence lifetime (Supplementary Note 2). The order of the above four design points is not strictly followed in the following experimental account of MM fabrication, characterization, and application, as the separate functions cannot be completely disentangled in the multifunctional molecular machine.

We validated our design by fabricating MM via annealing a scaffold strand and staple strands (Supplementary Tables 1 and 2). Successful self-assembly was indicated by a single band in agarose gel electrophoresis (Supplementary Fig. 6). MM was subsequently purified from non-incorporated staple strands via PEG-mediated precipitation, and structurally analyzed by transmission electron microscopy (TEM) after staining. The TEM-derived dimensions of 39 x 9 x 30 nm (Supplementary Figs. 7-9) are in close agreement with the Cadnano design^65^ (40 x 12 x 30 nm) as well as the dimensions predicted by oxDNA coarse-grain simulations^71^ (40 x 8 x 31 nm) (Supplementary Fig. 10a).

The nanomechanical actuation of MM was confirmed by reversibly opening and closing the device. The experiments were conducted using MM without any membrane anchoring and membranes. Successful actuation was established with gel electrophoresis as the gel band of MM shifted upon addition of opening or closing strands (Fig. 2b). The shifts were maintained at temperatures from 25 °C to 45 °C (Supplementary Fig. 11a) and across multiple open-closing cycles (Supplementary Fig. 11b). Furthermore, in TEM analysis the MM’s tilt angle shifted for the closed MM from 6.3 ± 3.3 ° (n = 128) to the value of 30.1 ± 0.8 ° (n = 131) for the open structure (Fig. 2c,d, Supplementary Figs. 12-13). The open tilt angle matches closely the design and the value of 29° obtained from oxDNA simulations (Supplementary Fig. 10b). The simulations also provided the spring constant for the cantilever (Supplementary Figs. 10 and 14) from the simulations’ RMSD.^72^ The spring constants were 4.5 ± 0.9 and 4.8 ± 1.6 pN/mm for the closed and open MM, respectively (Fig. 2e, Supplementary Fig. 14b), which is close to the range of 6 – 20 pN/nm of AFM cantilevers used for analysis of suspended membranes^73^.

We furthermore established that triggered opening of MM can be kinetically tracked using the Cy3 and Cy5 distance markers at base and beam, respectively (Fig. 2a). When closed-state MM was mixed with opening strands (OS), Cy5 emission decreased (Fig. 2a,f) and the Cy3 signal increased (Supplementary Figs.15 and 16), as anticipated. The halftime for the Cy5 change was 16.9 ± 0.4 min which confirms that membrane indentation is a time-independent elastic process and does not relate to the time-sensitive viscosity. Reversible and triggered MM opening and closing were confirmed by the full reversal of the fluorescence changes for Cy3 and Cy5 (Fig. 2a, Supplementary Fig. 17).

With functional nanodevices at hand, we tested whether MM actuation and indentation can distinguish membranes of different elasticities. We equipped MM with 30 cholesterol membrane anchors (Supplementary Fig. 18, Supplementary Tables 7 and 8) and added the devices to vesicles composed of stiffer POPC and softer DOPC membranes with 200 nm diameter (Fig. 3a, Supplementary Fig. 19). Both types of lipids are zwitterionic, exhibiting near-neutral zeta potential values at pH 8 (Supplementary Fig. 20). The two membranes’ elastic Young’s moduli of ∼12.4 MPa and 9.7 MPa, respectively, differ as a DOPC’s fatty acid chain features an extra double bond (Supplementary Table 9). MM bound equally well to both POPC and DOPC vesicles with *K*_d_ values of 24.0 ± 3.0 µM and 22.0 ± 2.2 µM, respectively, as established with gel electrophoresis (Supplementary Fig. 21). The binding relies on cholesterol anchors, as their absence abolished binding, as shown with gel electrophoresis (Supplementary Fig. 22). To test whether MM indentation distinguishes membrane elasticity (Fig. 3a), MM carrying fluorophore distance markers were bound to vesicles. MM binding to lipid membranes did not alter fluorescence emission of the markers (Supplementary Fig. 23). Actuation of MM by adding the 12 opening strands (Fig. 3b) led to a decrease in Cy5 emission (Fig. 3c), indicating the successful opening of the membrane-bound MM. Strikingly, the fluorescence drop at 21.7 ± 1.4 % for POPC was smaller than for DOPC at 28.1 ± 1.9 % (Fig. 3c,d, all OS) in agreement with a smaller indentation degree for stiffer POPC membranes.

Membrane indentation depth was proportional to the force applied by MM opening, as demonstrated by using different sets of opening strands. Following its design, MM can be opened by six short, six long, or all twelve opening strands (Fig. 3b, short OS, long OS, all OS, Supplementary Figs. 3 and 4, Supplementary Table 5). When OS hybridize to actuator strands and form extended DNA duplexes, force for indentation is generated, which should scale with the length and the number of duplexes (Supplementary Note 3). In line with expectations, the Cy5 signal change and indentation increased from short OS, over long OS to all OS for soft DOPC and stiffer POPC vesicles (Fig. 3d), suggesting that MM follows Hooke’s law. The experimental data are in line with biophysical modelling of the predicted force for opening MM with six short, six long and 12 OS (Supplementary Fig. 45, Supplementary Note 3). While sufficient to indent the membrane, the force generated by MM is too small to pull the cholesterol-anchored MM base out of the membrane (Supplementary Note 3). MM also successfully probed the elasticity of other membrane systems such as DOPC:POPC vesicles with a decreasing content of DOPC and thereby increasing stiffness (Fig. 3e), DPPC membranes which are stiffer than POPC^74^ (Fig. 3e), and DOPC vesicles at 5°C which show higher stiffness compared to at room temperature^75^ (Supplementary Fig. 24). The relative elasticities of the above membranes scaled well with membrane fluidity as measured using the membrane dye Laurdan (Supplementary Fig. 24). The cantilever tip is not compressed or otherwise deformed during actuation in oxDNA simulations of MM at membrane planes of varying stiffnesses (Supplementary Fig. 25). The membrane-elasticity dependent indentation by the MM tip and correlated opening of MM is supported by the behaviour of an MM version without a tip. This tip-free MM exhibited a false-positive fluorescence signal for full MM opening on DPPC membranes, even though a complete MM with tip only poorly indents this stiff membrane (Supplementary Fig. 26).

Following successful bilayer indentation, we wondered whether elasticity differences can also be visually mapped onto membranes. Therefore, Cy3/Cy5-tagged purified MMs (Supplementary Figs. 27 and 28) were bound to micron-sized giant unilamellar vesicles (GUVs) composed of either POPC or DOPC in PBS buffer (Supplementary Figs. 29 and 30). MM-decorated vesicles were then imaged with fluorescence lifetime imaging microscopy (FLIM) to obtain for each image pixel the average fluorescence decay lifetime *τ* of Cy3 (Fig. 3a, Supplementary Fig. 31). This key parameter is expected to reveal MM’s Cy3-Cy5 distance, the underpinning MM tilt angle and membrane indentation, and ultimately membrane elasticity (Fig. 3a, Supplementary Fig. 31, Supplementary Note 2). As FLIM does not suffer from interference by laser power fluctuations or varying localized fluorophore concentrations,^76^ quantitative measurements are facilitated.

To ease visualizing membranes with different elasticities, FLIM images were colour-coded so that each image pixel reflects the lifetime *τ* for MM at a given membrane position (Supplementary Fig. 31). For example, in the FLIM image for membrane-bound and closed MMs *τ* was uniform at 2.3 ± 0.1 ns for POPC and DOPC vesicles, as indicated by blue/green hues (Fig. 4a,b, Supplementary Figs. 32 and 33, *τ* refers to FastFLIM lifetime values unless stated otherwise). The image in Fig. 4a shows membrane vesicles with approximately 1 million bound MM nanodevices (calculation in the legend to Supplementary Fig. 31). This highlights the ability of the molecular machine to probe membrane elasticity in a parallel manner. Upon MM actuation with opening strands, *τ* strikingly increased to 3.1 ± 0.1 ns for POPC and to 3.4 ± 0.1 ns for DOPC (Fig. 4b,c, Supplementary Figs. 32 and 33). These significant changes represent less deformability of the stiffer POPC than the softer and more deformable DOPC membranes. The changes to higher lifetimes were reverted upon adding closing strands to 2.7 ± 0.1 ns and 2.5 ± 0.2 ns for POPC and DOPC, respectively (Supplementary Figs. 32 and 33). As another experimental support for the lifetime changes representing MM cantilever tilting, we used a control MM-Cy3 lacking Cy5 to mimic a FRET-free and maximally tilt-opened MM. In agreement with expectations, the corresponding *τ* was longer than the lifetimes of the above membrane-bound and actuated nanodevices, with a value of 3.4 ± 0.1 ns (Supplementary Fig. 34).

To obtain further quantitative insight into elastic membrane deformation by MM, we used the fitted *τ* values to calculate via FRET efficiencies and MM opening angles the membrane indentation depths (Fig. 4d, Supplementary Note 2, Supplementary Table 10). The indentation depths for DOPC and POPC were at −1.0 ± 0.2 nm and −0.8 ± 0.1 nm (Fig. 4e). The indentation values were further converted into bending rigidities *κ_c_* of 3.4 ± 0.5 x 10^-20^ J and 4.3 ± 0.7 x 10^-20^ J, respectively. The bending rigidities are closely related to the Young’s moduli (Supplementary Note 2) and are slightly lower compared to values obtained with other techniques (Supplementary Table 11), possibly due to some flexibility in the anchoring of MM to membranes (Supplementary Fig. 18). The analytical resolution for determining bending rigidity with MM was obtained with a bootstrapping process (Supplementary Note 4). The resolution was Δ*κ_c_* ∼ 0.3 x 10^-21^ J, with lower and upper bounds for the analytical range between *κ_c_* ∼ 3.2 x 10^-20^ and 8.0 x 10^-20^ J, respectively, which captures most relevant membranes^29,77,78^. A higher indentation degree for DOPC than POPC was also found using the fit-free phasor analysis, which indicated less FRET and hence more opening of MM at the softer membrane (Supplementary Fig. 35). The differential indentation depths were independently confirmed by determining MM’s indentation effect on bilayer organisation using the mechanosensitive fluorescent probe FliptR, which reports on lipid tension^79^ (Supplementary Note 5). FliptR-tracked tension changes furthermore confirmed that MM indentation is reversible upon adding closing strands (Supplementary Note 5). Interpreting the molecular indentation of membranes in terms of classical continuum mechanical elasticity and bending rigidity is valid because the MM tip is in direct contact with more than 140 lipid molecules extending to a larger deformation cap of 450 lipids per leaflet (Supplementary Note 5), which is at and beyond the threshold of around 150 lipids/leaflet where the molecular description converges with the mechanical parameters^67^. Furthermore, the interpretation is valid as membrane indentation depths are considerably larger than the amplitude of the thermal membrane fluctuations (Supplementary Note 6), and the min-long MM-based measurements are much longer than the microsecond-short membrane fluctuations (Supplementary Note 6), thus averaging and cancelling out thermal effects. MM action also induces membrane bending, but not membrane thinning, and bending rigidity dominates over membrane tension (Supplementary Note 6).

Encouraged by visualizing nanomechanical membrane indentation, we attempted to map different membrane elasticities within single vesicles that mimic the structural heterogeneity of biological cell membranes. As a synthetic model, we used GUVs with softer DOPC-rich liquid-disordered (Ld) domains as well as stiffer DPPC/cholesterol liquid-ordered (Lo) domains (Supplementary Fig. 36a). Closed-state MM bound to both membrane regions and yielded a uniform *_τ_* value of 2.3 ± 0.1 ns (Fig. 4b), as in the above-studied mono-lipid vesicles. A slight preferential MM binding to L_d_ regions was confirmed by specific staining of L_d_ (Supplementary Fig. 36b,c) in line with other membrane-tethering DNA nanostructures^80^.

When actuated, MM differentiated stiffer L_o_ regions and softer L_d,_ with *τ* increases to 2.8 ± 0.2 ns and 3.2 ± 0.1 ns, respectively, in single vesicles (Fig. 4a-c, Supplementary Fig. 32c). The relative *τ* change for the L_o_ DPPC/cholesterol regions (2.3 ns to 2.8 ns) is lower than for the pure POPC vesicles (2.3 ns to 3.1 ns) as both cholesterol and chain saturation increase membrane rigidity. Similarly, the change for DOPC-rich L_d_ region (2.3 to 3.2 ns) is lower than that of pure DOPC GUVs (2.3 ns to 3.4 ns), since partial partitioning of cholesterol increases the stiffness of the L_d_ domains^81^, as supported by analysing membrane tension with FliptR (Supplementary Note 5). We note a non-linear relationship between the MM-derived membrane rigidity and FliptR-reported membrane tension (Supplementary Note 5, last figure) which reflects the correlated molecular interactions underpinning these two readouts^82^.

As a final test of MM’s utility for measuring bilayer nanomechanics, we probed live human cells to determine any elasticity differences between the plasma membrane and internal organelle membranes, after MM uptake by cells. When HeLa cells were incubated, MMs were rapidly internalized into late endosomes and lysosomes (co-staining see Supplementary Fig. 37) which is often found for other DNA nanostructures^83–86^. The addition of the endocytosis inhibitor Dynasore^87–90^ slowed down MM internalization to allow for a 40 min-long FLIM analysis (Supplementary Fig. 38). Details about the cell culture and imaging conditions are provided in the Methods section.

In FLIM images of cells with closed MM, signals were found for both the plasma membrane (PM) and the membranes of internal vesicles (INT) (Fig. 5a,b), with an average lifetime of 0.8 ± 0.1 ns uniformly across all membrane regions, which is lower than for synthetic vesicles. The FRET readout was unchanged at pH 4.8 compared with neutral conditions, indicating that acidic pH alone does not account for the altered intracellular lifetime (Supplementary Fig. 39). Nor is the MM function affected by typical intracellular or extracellular nucleases (Supplementary Fig. 40). We do not rule out that the fluorescent dyes of MM are influenced by the physicochemical environments at the cell membranes^91,92^, possible stacking interactions with cellular poteins, or partial cleavage of the fluorophores from the MM surface by other non-tested nucleases, something which can be prevented by adapting the MM design to position the dyes in a more protected environment, such as the nanospace between cantilever and base (Supplementary Fig. 4). When accounting for this lower contribution by fitting the multicomponent lifetime distribution (Supplementary Fig. 41) we obtained a longer *τ* value of 2.9 ± 0.2 ns.

**Fig. 5.**
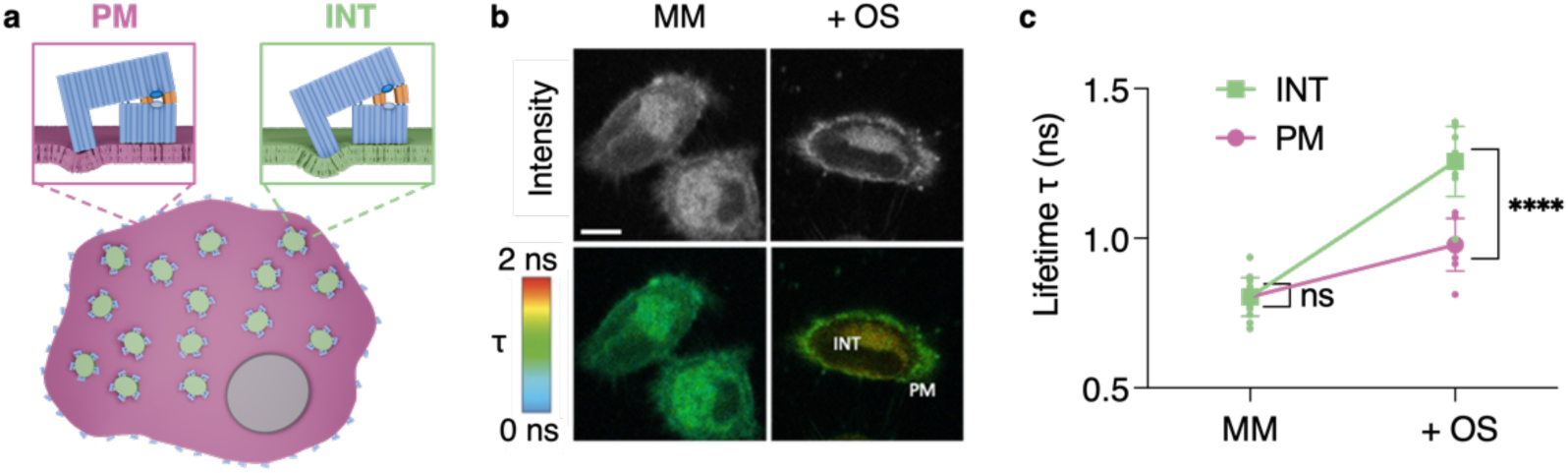
**FLIM analysis of the MM in live HeLa cells establishes bilayer regions of varying stiffness**. **a,** Schematic of a HeLa cell with bound MM to the stiffer outer plasma membrane (magenta) and softer internal membrane (green) as found in FLIM analysis. The cell nucleus is shown in dark grey. **b**, Intensity and Fast-FLIM images of MM in live HeLa cells and upon addition of opening strands (+OS). Plasma membrane (PM) and internal (INT) membrane regions are labeled. The pixels are coloured according to the lifetime (scale 0 - 2ns) and have a corresponding point in the phasor plot in Supplementary Fig. 40. Scale bar, 10 µm. **c**, The line plot displays the mean average lifetimes ± SD calculated by phasor analysis in the internal membrane ROI (INT) and the plasma membrane region of interest (PM). One-way ANOVA, Tukey’s multiple comparison test, ****P <0.0001; n = 10 cells per condition, across three independent experiments.

Actuation of cell-bound MM with opening strands uncovered average lifetime increases of 1.0 ± 0.1 ns for PM and 1.3 ± 0.1 ns for INT regions (Fig. 5a-c) which is below 1.5 ± 0.1 ns from control MM-Cy3 (Supplementary Fig. 42). When focusing on the long-lifetime component, PM membrane regions had a lifetime of 3.1 ± 0.2 ns and INT regions 3.2 ± 0.1 ns, which are in closer agreement with values obtained for synthetic membranes. The lower lifetime values at PM indicate that the plasma membrane is less elastic than the softer internal membranes^93,94^. Different elasticities at cellular membrane regions were also apparent in the FLIM decay curves and FLIM phasor plots showing more FRET efficiency and lower MM tilt angles at PM than INT membrane regions (Supplementary Figs. 42, 43, 44).

Top-down nanotechnology thrives by drawing inspiration from macroscopic principles and miniaturizing them, as demonstrated by mimicking the record player to devise AFM cantilevers. In this report, we have asked the simple question of whether the principle of indenting a compressible material with a tip can be miniaturized to the nanoscale and achieve scientific benefits over existing mechanoanalysis methods. We answered these questions by harnessing bottom-up DNA nanotechnology to build nanoscale tips and using them to determine the biologically and disease-relevant nanomechanics of lipid bilayer membranes. To be functional, the new 10 nm-sized tips were integrated into a molecular machine which binds to, actuates, senses, and reports on membrane elasticity. Compared to classical serial top-down mechanoanalysis, the molecular machines’ small size and self-contained multifunctional operation allow deploying millions of nanodevice copies for parallel analysis. Conventionally inaccessible membranes such as those of internal organelles are also probed, as demonstrated by the pioneering co-analysis of stiff plasma and soft internal cell membranes. The stiffness of the plasma membrane is likely related to its high content of cholesterol and saturated glycerophospholipids, while the more elastic internal membrane may reflect lower cholesterol levels and more unsaturated lipids^95–97^ , which alter membrane lipid order^94,98^. It is also possible that the stiffer PM membrane is interlinked to the underlying cytoskeleton which responds to external deforming force^99,100^. This may be explored in future studies by using the molecular machines’ indentation depth of around 1 nm to decouple the bilayer nanomechanics from the underlying protein cell cortex layer, which are usually probed together with classical top-down probes of greater indentation depth, even though bimodal AFM and other multifrequency techniques can achieve small amplitudes.

As another unique feature, our molecular machines’ minute-long opening kinetics measure elastic properties of the membrane rather than time-dependent viscosity, which is usually probed with second-to millisecond-fast oscillating AFM cantilevers. Faster actuation or self-oscillation may, however, be built into our molecular machine by harnessing recent advances in dynamic DNA nanotechnology^101–104^. The DNA-based machines for membrane mechanoanalysis complement other DNA nanodevices previously engineered for measuring biophysical cell parameters such as intracellular pH or ionic concentration.^105–109^ In future studies, the molecular machines’ nanomechanical analysis of external and internal cell membranes could provide insight into biologically relevant topics including endocytosis^110,111^ and cell division^112^ when the nanodevices are located at cell-cell interfaces; the latter are usually inaccessible for conventional top-down tools. Of equal interest is nanomechanically regulated cell fate^113^ , which controls cell division or apoptosis^114^, or stem cell differentiation.^115^ In cancer biology, membrane elasticity differs between cancerous and noncancerous cells^116,117^, and the molecular machines could help better understand the influence of elasticity on cancer cell motility and adhesion. A further application could be embryonic development, where mechanical properties help select viable embryos^118^. Current methods, such as AFM, require tissue removal, while invasive techniques like membrane microaspiration and ultraviolet laser cuts can disrupt embryo development^119^. Our molecular machines may provide an alternative opportunity to infer these properties in a non-invasive manner without disrupting the embryo’s development. In conclusion, our report mimics top-down nanotechnology with bottom-up nanotechnology to advance nanomechanobiology.

## METHODS

### Materials

DNA strands, 3’ labeled DNA-TEG-Chol and 5’ labeled DNA-Cy3 oligonucleotides as well as Cy3 and Cy5 labeled DNA oligonucleotides were purchased from Integrated DNA Technologies (UK) or Eurogentec (BE) with HPLC or PAGE purification. Phix174 scaffold was purchased from New England Biolabs (UK). Lipids 1-palmitoyl-2-oleoyl-glycero-3-phosphocholine (POPC), 1,2-dioleoyl-sn-glycero-3-phosphocholine (DOPC), 1,2-dipalmitoyl-sn-glycero-3-phosphocholine (DPPC) and cholesterol (plant-derived) were purchased dissolved in chloroform from Avanti Polar Lipids (UK). All other reagents were purchased from Sigma-Merck (UK) unless stated otherwise.

### MM fabrication

The DNA sequences and 2D connectivity map of the DNA MM are provided in Supplementary Table 2-6 and Supplementary Figure S1. To self-assemble MM at a concentration of 50 nM, Phix174 virion DNA scaffold (10 µL, 500 nM) was mixed with 6 x excess of 112 staple strands (32.5 µ, 100 µM; Supplementary Table 1), 5 x excess of cholesterol anchoring strands (6 µL, 100 µM; Supplementary Table 7), 1 x TAE 160 mM MgCl_2_ (10 µL) and topped up with deionized H_2_O to a total volume of 100 µL. For fluorescence experiments, staple strands main025 and main078 were replaced with Cy3 and Cy5-labeled versions at 3 x fold excess (15 µL, 10 µM). 12 opening strands (Supplementary Table 5) and 12 closing strands (Supplementary Table 6) were added in 5x excess (3.6 µL, 100 µM). The DNA nanostructure was folded in a Thermocycler (Biorad, US) by heating to 80 °C for 10 min following 10 min at 65 °C. Next, the mixture was cooled down for one degree every hour to 25 °C, and then cooled one degree every 10 min to 15 °C. Volumes were adopted accordingly for assembly at the 20 nM scale.

### MM purification

For purification by PEG precipitation, folded MM (100 µL, 20-50 nM) was mixed with 200 µL of buffer 1 (15% PEG 800, 1x TAE, 0.5 mM NaCl), vortexed for 3 s and centrifuged for 30 min at 10,000 r.c.f. at 25 °C. The orientation of the tube placed in the centrifuge was labeled to indicate the position of the forming pellet. The supernatant was removed after centrifugation and 100 µL of buffer 2 (1x TAE, 16 mM MgCl_2_) was added, vortexed for one min and incubated at 37 °C and 1000 rpm in a heating block (Biorad, US) for 20 h.

Purification by size exclusion chromatography (SEC) was conducted for microscopy experiments. Assembled MM was purified from excess staple strands using SEC via an ÄKTA purifier (Cytiva, US) with a Superdex 75 increase packed 10/300 GL column (GE Healthcare, US). The sample was eluted with degassed 16 mM MgCl_2_ buffer at a flow rate of 0.5 mL / min and by monitoring the absorbance at 260 and 280 nm. The eluent was collected on a 96-well plate and fractions containing MM were analysed via agarose gel electrophoresis.

### Membrane vesicle formation

To form small unilamellar vesicles (SUVs), a solution of POPC or DOPC lipids in chloroform (10 mM, 100 μL) was added to a 2 mL screw-top glass vial. The solvent was removed using argon gas, while rotating to yield a thin lipid film. The lipid was resuspended in SUV buffer (50 mM HEPES 500 mM NaCl, or 1 x TAE 16 mM MgCl_2_; 1 mL), the solution was vortexed for 30 s and then sonicated for 1 min at RT. The SUV vesicle solution was extruded 25 times through an Avanti Mini-Extruder (Avanti Polar Lipids, US) with 200 nm filter paper (Avanti Polar Lipids, US). Dynamic light scattering (DLS) was used to confirm the vesicle diameter via a Zetasizer Nano S (Malvern Panalytical, UK). The ζ-potentials of the vesicles were measured using a Zetasizer Nano S (Malvern Panalytical,). The SUVs were diluted to a final lipid concentration of 100 µM in buffer (50 mM HEPES, 500 mM NaCl, pH 8), and measurements were performed at 25 °C. The mean ζ-potential was calculated from three replicate measurements (n = 3).

To form giant unilamellar vesicles (GUV), a solution of POPC or DOPC lipids (36 µL, 10 mg / mL) in chloroform was placed within a rubber O-ring on an indium-tin-oxide (ITO) (Nanion Technologies, DE) coated glass slide and allowed to dry to form a lipid film. The ITO slide was inserted in a vesicle prep device (Nanion Technologies, DE). Sucrose (250 µL, 400 mM) was added to the dried lipid and another ITO glass slide was placed on top to close the chamber. The GUVs were formed via electroformation by using the voltage program with 10 Hz, 4V amplitude, 1 min rise, 100 min main time and 10 min fall time. The temperature was set at 70 °C for the mixed-phase GUVs and 27 °C for DOPC and POPC GUVs.

### Characterization of MM with agarose gel electrophoresis

A 1% agarose gel in 0.5 x TBE 10 mM MgCl_2_ with ethidium bromide (1:30000, 5 µL in 150 mL) was used to determine the assembly of MM. 5-10 µL of DNA (20 nM) were mixed with gel loading dye (purple 6x, no SDS; New England Biolabs, UK) and loaded onto the gel. The gel was run for 1-2 hours (depending on the degree of the desired separation) at 65 V and 4 °C. To distinguish between the opened and closed conformation of the DNA MM, a 2.5% agarose gel was run for 2.5 hours at 65 V and 4 °C. The bands were analysed using an Azure Biosystems C Series Imaging Systems (Azure Biosystems, US).

Agarose gel electrophoresis was also used to confirm the binding of MM to SUV membrane vesicles. Purified MM (7 μL, 20 nM) were incubated with cholesterol strands in 5x access (4.2 µL, 10 µM) for 15 min and mixed with SUVs (POPC or DOPC) (7 μL, 20-200 µM). The DNA-MM SUV mixture was incubated for 1 hr at RT. The mixture was analysed using 1% agarose gel with ethidium bromide (1:30000, 5 µL in 150 mL) in TBE (0.5 x TBE 10 mM MgCl_2_). The gel was run at 65 V for 90 min at 4 °C. The gel bands were visualized using an Azure Biosystems C Series Imaging Systems (Azure Biosystems, US).

### Transmission electron microscopy

Purified MM (6 µL, 3-5 nM) were placed on a glow-discharge treated carbon-coated Cu300 mesh grid and stained with 2 % uranyl formate. The images were obtained with a JEM 2100 electron microscope and Orius SC200 CCD camera at 200Kv (JEOL, JP).

### Spectrometric FRET analysis of MM

The fluorescence emission was measured with a Varian Eclipse fluorescence spectrophotometer (Agilent, UK). The excitation slit was set at 5 nm, the emission slit at 10 nm, the scan rate at 600 nm / min and excitation at 540 nm and 600 PMT voltage. 100 µL of the DNA MM was added to a quartz cuvette with a 10 mm path length (Hellma, DE). For kinetics, the excitation wavelength was set at 550 nm, the emissions were scanned between 570 and 670 nm. Opening or closing strands in 5x excess were added after 5 min. The data was normalised onto the average intensity before the addition of the strands and percentual change to the initial intensities recorded.

### oxDNA analysis

MM design files for the “open” and “closed” form were prepared in caDNAno^65^ and converted to oxDNA input files using tacoxDNA^120^. Simulations were performed using the oxDNA DNA2 model^121^ using 0.4 M salt equivalents, as a compromise between system stability and compute time.^122^ Structures were minimized for 2000 steps with a timestep of 0.005 before being relaxed and brought to 300 K using the Bussi-Donadio-Parrinello thermostat for a further 10^5^ steps, following protocols detailed in Matthies et al^123^. Both equilibrium and applied force simulations were then run with an Andersen-like thermostat for up to 10^7^ steps, with nucleotide velocities refreshed every 10^3^ steps and the diffusion coefficient set to 2.5. Analysis was performed using the oxDNA Analysis Tools package and visualisations in R; including ggplot^124^, VMD and Blender.

To simulate nuclease degradation, the simulations were conducted after removing a random selection of staple strands at 5% and 15% of the total from the caDNAno file. To determine the spring constants for these partial MM structures, the average RMSD of the oxDNA beads from the final ∼20% of the equilibrated trajectories was determined and compared to an idealised structure with all B-form DNA helices intact. Spring constants *k* were calculated as *k* = kB * T / x^2^, where kB is the Boltzmann constant in pN·nm/K, T is the temperature in Kelvin, and x is the averaged RMSD value. The error for this calculation was determined as the derivative of *k* with respect to x using dk/dx = −2 * kB * T / x^3^.

To simulate the actuation of the MM and to calculate the indentation depth and membrane-induced distortion of the cantilever tip, MM was in the closed conformation at the beginning of the simulation, with the opening strands partially attached to allow their rapid annealing to the actuator strands and cause actuation of MM. In the simulations, MM was placed atop a membrane-mimicking resistive plane of various stiffnesses. MM was tethered to the plane via 32 harmonic traps to mimic the 32 cholesterol anchors of MM. The stiffnesses of the plane varied from zero-force with no resistance against indentation to membrane planes with higher stiffness, including the “Soft” membrane with a stiffness value of 0.06 oxDNA units comparable to that of a DOPC membrane^125^, the “Stiff” membrane has an oxDNA stiffness of 1, and the “Stiffest” membrane is set to maximum resistance. In the simulations, MM-mediated indentation caused a greater energetic penalty the further the MM tip penetrated through the plane.

### GUV experiments with MM

MM (15 µL, 50 nM) was added to a solution of GUVs (15 µL) in 1x PBS buffer (150 µL) in a µ-Dish 35 mm, high (Ibidi, US) fitted with a 6.96 mm diameter insert. For FliptR FLIM experiments, the MM (15 µL, 50 nM) was added to a solution of GUVs in 1x PBS supplemented with FliptR (2 mM) under the same conditions. MM and GUVs were left for 15 min at RT and subjected to microscopy and lifetime imaging. For the MM actuating experiments, Fluorescent lifetime imaging microscopy (FLIM) measurements were carried out only after 10 min incubation with opening strands (OS) (0.9 µL, 8.3 µM) or closing strands (CS) (0.9 µL, 8.3 µM).

### Cell tissue experiments with MM

Wild-type HeLa cells (a gift from Prof Jeremy Carlton, King’s College London) were cultured in Dulbecco’s Modified Eagle Medium (DMEM) (Gibco, Life Technologies, US), supplemented with 10 % fetal bovine serum (FBS) and 1 % Penicillin–Streptomycin in T-75 flasks (Sigma-Merck (UK)). The cultures were maintained in a humidified atmosphere containing 5% CO_2_ at 37 °C and passaged every ∼3 days. Cells were plated at a density of 5,000 cells per well in a µ-Dish 35 mm, high (Ibidi, US) fitted with a 6.96 mm diameter insert. The following day, the cells were washed with 1x PBS and treated with Dynasore (50 µM) for 30 min to inhibit clathrin-mediated endocytosis. The cells were washed three times with 1x PBS and then incubated with MM in 1x PBS (5 nM) for 5 min at 37 °C. The cells were washed three times again with PBS to remove any non-bound DNA and subjected to microscopy and lifetime imaging. For the MM actuating experiments, FLIM measurements were carried out following 10 min incubation with opening strands (OS) (8.3 µM, 0.9 µL) after MM preincubation for 5 min. FLIM imaging of MM opening and closing was carried out within a 45 min timeframe after adding MM to avoid readings of low DNA signal and photon counts. Statistical analysis was conducted using Prism GraphPad (https://www.graphpad.com/features).

### Confocal laser scanning microscopy

Fluorescence microscopy images were taken using a 60x oil objective on the Zeiss Airyscan LSM 880 (University College London) or Nikon A1 (Nikon Imaging Centre, King’s College London). Both imaging systems are equipped with an environmental chamber to maintain cells at 37 °C and 5 % CO_2_. Images were taken using laser excitation of 532 nm (Cy3) to determine any DNA-membrane interaction. All images were taken using the same exposure and gain settings. Image analysis was carried out using ImageJ ((https://imagej.net)) and statistical analysis using Prism GraphPad.

### Fluorescence lifetime imaging microscopy

Multiphoton fluorescence lifetime imaging microscopy (FLIM) was conducted on the Leica TCS SP8-FALCON (FAst Lifetime CONtrast) system (Leica Microsystems GmbH, Wetzlar, DE) using a 40x air objective. Sensitive hybrid detectors (SMD, Leica Microsystems CMS GmbH, Mannheim, DE) allowed for photon detection, while a 910 nm pulsed laser (80 MHz) was used to excite FliptR and 760 nm to excite Cy3. FLIM images were acquired at 512 x 512 pixels. At least 10,000 photon events were recorded in each measurement. The photon events were time-binned into a histogram and a fluorescence decay curve was obtained. FastFLIM images (average photon arrival times per pixel) are shown. Regions of interest (ROI) were drawn to determine the lifetime of different vesicle phases or cell compartments. For FliptR FLIM fit analysis, the SP8 Falcon fitting model ‘n-exponential reconvolution’ was used for the biexponential fitting of the fluorescence decay curve, to extract the FliptR lifetimes *τ*_1_ and *τ*_2_. The parameter chi-squared (χ2) was used as an indicator of the quality of the curve fitting. The longest lifetime (*τ*_2_) was used to report on membrane tension, while *τ*_1_ only accounted for the minority of the signal due to a low photon number. Prism GraphPad was used to plot the average lifetime and conduct statistical analysis. For FRET-FLIM, phasor analysis, the data was analysed using an integrated FLIM-phasor analysis, enabling a global view of the fluorescence decay in each pixel of the measured image. FRET efficiency was determined using a FRET trajectory, which combines the phasor of the unquenched donor, background signal, and the experimental points in the phasor plot to determine the number of donors undergoing FRET and the corresponding FRET efficiency. This fit-free approach does not resolve the decay into exponential components.

## Notes

### Competing Interest Statement

The authors have declared no competing interest.

### Funding

- BBSRC: BB/X001342/1

## References

1. Atilla-Gokcumen, G. E. et al. Dividing cells regulate their lipid composition and localization. Cell 156, 428–439 (2014).

2. Fletcher, D. A. & Mullins, R. D. Cell mechanics and the cytoskeleton. Nature 463, 485–492 (2010).

3. Groves, J. T. Membrane mechanics in living cells. Dev. Cell 48, 15–16 (2019).

4. Shi, Z., Graber, Z. T., Baumgart, T., Stone, H. A. & Cohen, A. E. Cell membranes resist flow. Cell 175, 1769–1779.e13 (2018).

5. Chaudhuri, P. K., Low, B. C. & Lim, C. T. Mechanobiology of tumor growth. Chem. Rev. 118, 6499–6515 (2018).

6. Cross, S. E., Jin, Y. S., Rao, J. & Gimzewski, J. K. Nanomechanical analysis of cells from cancer patients. Nat. Nanotechnol. 2, 780–783 (2007).

7. Plodinec, M. et al. The nanomechanical signature of breast cancer. Nat. Nanotechnol. 7, 757–765 (2012).

8. Hamidi, H. & Ivaska, J. Every step of the way: Integrins in cancer progression and metastasis. Nat. Rev. Cancer 18, 533–548 (2018).

9. Mittelheisser, V. et al. Evidence and therapeutic implications of biomechanically regulated immunosurveillance in cancer and other diseases. Nat. Nanotechnol. 19, 281–297 (2024).

10. Kariuki, S. N. et al. Red blood cell tension protects against severe malaria in the dantu blood group. Nature 585, 579–583 (2020).

11. Arakawa, C. et al. Biophysical and biomolecular interactions of Malaria-infected erythrocytes in engineered human capillaries. Sci. Adv. 6, eaay7243 (2020).

12. Barabino, G. A., Platt, M. O. & Kaul, D. K. Sickle cell biomechanics. Annu. Rev. Biomed. Eng. 12, 345–367 (2010).

13. Cao, H. et al. Red blood cell mannoses as phagocytic ligands mediating both sickle cell anaemia and malaria resistance. Nat. Commun. 12, 1792 (2021).

14. Kasahara, K. et al. Spatiotemporal single-cell tracking analysis in 3D tissues to reveal heterogeneous cellular response to mechanical stimuli. Sci. Adv. 9, eadf9917 (2023).

15. Wolfenson, H., Yang, B. & Sheetz, M. P. Steps in mechanotransduction pathways that control cell morphology. Annu. Rev. Physiol. 81, 585–605 (2019).

16. Reuten, R. et al. Basement membrane stiffness determines metastases formation. Nat. Mater. 20, 892–903 (2021).

17. Nia, H. T. et al. Solid stress and elastic energy as measures of tumour mechanopathology. *Nat*. Biomed. Eng. 1, 1–11 (2016).

18. McCarthy, N. L. C., Ces, O., Law, R. V., Seddon, J. M. & Brooks, N. J. Separation of liquid domains in model membranes induced with high hydrostatic pressure. Chem. Commun. 51, 8675–8678 (2015).

19. Páez-Pérez, M., López-Duarte, I., Vyšniauskas, A., Brooks, N. J. & Kuimova, M. K. Imaging non-classical mechanical responses of lipid membranes using molecular rotors. Chem. Sci. 12, 2604–2613 (2021).

20. Ron, A. et al. Cell shape information is transduced through tension-independent mechanisms. Nat. Commun. 8, 2145 (2017).

21. Iskratsch, T., Wolfenson, H. & Sheetz, M. P. Appreciating force and shape — the rise of mechanotransduction in cell biology. Nat. Rev. Mol. Cell Biol. 15, 825–833 (2014).

22. van Meer, G., Voelker, D. R. & Feigenson, G. W. Membrane lipids: where they are and how they behave. Nat. Rev. Mol. Cell Biol. 9, 112–124 (2008).

23. Lekka, M. & Laidler, P. Applicability of AFM in cancer detection. Nat. Nanotechnol. 4, 72–72 (2009).

24. Guz, N. V., Patel, S. J., Dokukin, M. E., Clarkson, B. & Sokolov, I. AFM study shows prominent physical changes in elasticity and pericellular layer in human acute leukemic cells due to inadequate cell-cell communication. Nanotechnology 27, 494005 (2016).

25. Gavara, N. & Chadwick, R. S. Determination of the elastic moduli of thin samples and adherent cells using conical atomic force microscope tips. Nat. Nanotechnol. 7, 733–736 (2012).

26. Barak, P., Rai, A., Rai, P. & Mallik, R. Quantitative optical trapping on single organelles in cell extract. Nat. Methods 10, 68–70 (2013).

27. Neuman, K. C. & Nagy, A. Single-molecule force spectroscopy: optical tweezers, magnetic tweezers, and atomic force microscopy. Nat. Methods 5, 491–505 (2008).

28. Jensen, L. E. et al. Membrane curvature sensing and stabilization by the autophagic LC3 lipidation machinery. Sci. Adv. 8, eadd1436 (2022).

29. Sayem Karal, M. A., Masum Billah, M., Ahmed, M. & Kabir Ahamed, M. A review on the measurement of the bending rigidity of lipid membranes. Soft Matter 19, 8285–8304 (2023).

30. Pu, H. et al. Micropipette aspiration of single cells for both mechanical and electrical characterization. IEEE Trans. Biomed. Eng. 66, 3185–3191 (2019).

31. Pereno, V. et al. Electroformation of giant unilamellar vesicles on stainless steel electrodes. ACS Omega 2, 994–1002 (2017).

32. Lüchtefeld, I. et al. Dissecting cell membrane tension dynamics and its effect on Piezo1-mediated cellular mechanosensitivity using force-controlled nanopipettes. Nat. Methods 21, 1063–1073 (2024).

33. Lekka, M. Discrimination between normal and cancerous cells using AFM. BioNanoScience 6, 65–80 (2016).

34. Rothemund, P. W. K. Folding DNA to create nanoscale shapes and patterns. Nature 440, 297–302 (2006).

35. Douglas, S. M. et al. Self-Assembly of DNA into nanoscale three-dimensional shapes. Nature 459, 414–418 (2009).

36. Jabbari, H., Aminpour, M. & Montemagno, C. Computational approaches to nucleic acid origami. ACS Comb. Sci. 17, 535–547 (2015).

37. Pfeifer, W. & Saccà, B. From nano to macro through hierarchical self-assembly: The DNA paradigm. ChemBioChem 17, 1063–1080 (2016).

38. Blanchard, A. T. & Salaita, K. Emerging uses of DNA mechanical devices. Science 365, 1080–1081 (2019).

39. Yao, G. et al. Meta-DNA structures. Nat. Chem. 12, 1067–1075 (2020).

40. Dey, S. et al. DNA origami. Nat. Rev. Methods Primer 1, 13 (2021).

41. Gür, F. N. et al. Double-to single-strand transition induces forces and motion in DNA origami nanostructures. Adv. Mater. 33, 2101986 (2021).

42. Zhang, L. et al. High-performance organohydrogel artificial muscle with compartmentalized anisotropic actuation under microdomain confinement. Adv. Mater. 35, 2202193 (2023).

43. Sato, Y. et al. Environment-dependent self-assembly of DNA origami lattices on phase-separated lipid membranes. Adv. Mater. Interfaces 5, 1–6 (2018).

44. Rubio-Sánchez, R., Barker, S. E., Walczak, M., Cicuta, P. & Michele, L. D. A modular, dynamic, DNA-based platform for regulating cargo distribution and transport between lipid domains. Nano Lett. 21, 2800–2808 (2021).

45. Maingi, V. et al. Digital nanoreactors to control absolute stoichiometry and spatiotemporal behavior of DNA receptors within lipid bilayers. Nat. Commun. 14, 1532 (2023).

46. Du, Y. et al. Membrane-anchored DNA nanojunctions enable closer antigen-presenting cell–T-cell contact in elevated T-cell receptor triggering. Nat. Nanotechnol. 18, 818–827 (2023).

47. Yu, Z. et al. Compliant DNA origami nanoactuators as size-selective nanopores. Adv. Mater. 2405104 (2024).

48. Xing, Y., Rottensteiner, A., Ciccone, J. & Howorka, S. Functional nanopores enabled with DNA. Angew. Chem. Int. Ed. 62, e202303103 (2023).

49. Diederichs, T. et al. Principles of small-molecule transport through synthetic nanopores. ACS Nano 15, 16194–16206 (2021).

50. Howorka, S. Building membrane nanopores. Nat. Nanotechnol. 12, 619–630 (2017).

51. Burns, J. R., Seifert, A., Fertig, N. & Howorka, S. A biomimetic DNA-based channel for the ligand-controlled transport of charged molecular cargo across a biological membrane. Nat. Nanotechnol. 11, 152–156 (2016).

52. Langecker, M. et al. Synthetic lipid membrane channels formed by designed DNA nanostructures. Science 338, 932–936 (2012).

53. Franquelim, H. G., Khmelinskaia, A., Sobczak, J. P., Dietz, H. & Schwille, P. Membrane sculpting by curved DNA origami scaffolds. Nat. Commun. 9, 811 (2018).

54. Kurokawa, C. et al. DNA cytoskeleton for stabilizing artificial cells. Proc. Natl. Acad. Sci. U. S. A. 114, 7228–7233 (2017).

55. Franquelim, H. G., Dietz, H. & Schwille, P. Reversible membrane deformations by straight DNA origami filaments. Soft Matter 17, 276–287 (2021).

56. Grome, M. W., Zhang, Z., Pincet, F. & Lin, C. Vesicle tubulation with self-assembling DNA nanosprings. Angew. Chem. Int. Ed. 57, 5330–5334 (2018).

57. Grome, M. W., Zhang, Z. & Lin, C. Stiffness and membrane anchor density modulate DNA-nanospring-induced vesicle tubulation. ACS Appl. Mater. Interfaces 11, 22987–22992 (2019).

58. Baumann, K. N. et al. Coating and stabilization of liposomes by clathrin-inspired DNA self-assembly. ACS Nano 14, 2316–2323 (2020).

59. Marras, A. E., Zhou, L., Su, H. J. & Castro, C. E. Programmable motion of DNA origami mechanisms. Proc. Natl. Acad. Sci. U. S. A. 112, 713–718 (2015).

60. Paez-Perez, M., Russell, I. A., Cicuta, P. & Di Michele, L. Modulating membrane fusion through the design of fusogenic DNA circuits and bilayer composition. Soft Matter 18, 7035–7044 (2022).

61. Stengel, G., Zahn, R. & Höök, F. DNA-induced programmable fusion of phospholipid vesicles. J Am Chem Soc 129, 9584–9585 (2007).

62. Löffler, P. M. G., et al. A DNA-programmed liposome fusion cascade. Angew. Chem. Int. Ed. 56, 13228–13231 (2017).

63. Jahn, R., Cafiso, D. C. & Tamm, L. K. Mechanisms of SNARE proteins in membrane fusion. Nat. Rev. Mol. Cell Biol. 25, 101–118 (2024).

64. Jahnke, K., Huth, V., Mersdorf, U., Liu, N. & Göpfrich, K. Bottom-Up assembly of synthetic cells with a DNA cytoskeleton. ACS Nano 16, 7233–7241 (2021).

65. Douglas, S. M. et al. Rapid prototyping of 3D DNA-Origami shapes with caDNAno. Nucleic Acids Res. 37, 5001–5006 (2009).

66. Wilton, D. J., Ghosh, M., Chary, K. V. A., Akasaka, K. & Williamson, M. P. Structural change in a B-DNA helix with hydrostatic pressure. Nucleic Acids Res. 36, 4032– 4037 (2008).

67. Li, J., Thomson, S. & Monje, V. Impact of System Size on the Biophysical Properties of Multicomponent Lipid Bilayers: A Systematic Study Using Molecular Dynamics Simulations. J. Chem. Inf. Model. 65, 12464–12474 (2025).

68. Paez-Perez, M. & Kuimova, M. K. Molecular rotors: Fluorescent sensors for microviscosity and conformation of biomolecules. *Angew. Chem. Int. Ed*. 63, e202311233 (2024).

69. Polita, A., Toliautas, S., Žvirblis, R. & Vyšniauskas, A. The effect of solvent polarity and macromolecular crowding on the viscosity sensitivity of a molecular rotor BODIPY-C10. Phys. Chem. Chem. Phys. 22, 8296–8303 (2020).

70. Jepsen, M. D. E. et al. Single molecule analysis of structural fluctuations in DNA nanostructures. Nanoscale 11, 18475–18482 (2019).

71. Poppleton, E., Romero, R., Mallya, A., Rovigatti, L. & Šulc, P. OxDNA.org: A public webserver for coarse-grained simulations of DNA and RNA nanostructures. Nucleic Acids Res. 49, W491–W498 (2021).

72. Yoo, J. & Aksimentiev, A. In situ structure and dynamics of DNA origami determined through molecular dynamics simulations. Proc. Natl. Acad. Sci. 110, 20099–20104 (2013).

73. Steltenkamp, S. et al. Mechanical properties of pore-spanning lipid bilayers probed by atomic force microscopy. Biophys. J. 91, 217–226 (2006).

74. Nanda, H. et al. Relaxation dynamics of saturated and unsaturated oriented lipid bilayers. Soft Matter 14, 6119–6127 (2018).

75. Pan, J., Tristram-Nagle, S., Kučerka, N. & Nagle, J. F. Temperature dependence of structure, bending rigidity, and bilayer interactions of dioleoylphosphatidylcholine bilayers. Biophys. J. 94, 117–124 (2008).

76. Periasamy, A., Mazumder, N., Sun, Y., Christopher, K. G. & Day, R. N. FRET microscopy: Basics, issues and advantages of FLIM-FRET imaging. in Advanced Time-Correlated Single Photon Counting Applications vol. 111 249–276 (Springer International Publishing, Cham, 2015).

77. Himbert, S. et al. The bending rigidity of the red blood cell cytoplasmic membrane. PLoS ONE 17, e0269619 (2022).

78. Gracià, R. S., Bezlyepkina, N., Knorr, R. L., Lipowsky, R. & Dimova, R. Effect of cholesterol on the rigidity of saturated and unsaturated membranes: fluctuation and electrodeformation analysis of giant vesicles. Soft Matter 6, 1472 (2010).

79. Colom, A. et al. A fluorescent membrane tension probe. Nat. Chem. 10, 1118– 1125 (2018).

80. Bennett, I. D., Burns, J. R., Ryadnov, M. G., Howorka, S. & Pyne, A. L. B. Lipidated DNA nanostructures target and rupture bacterial membranes. Small **n/a**, 2207585.

81. Ma, Y. et al. Cholesterol partition and condensing effect in phase-separated ternary mixture lipid multilayers. Biophys. J. 110, 1355–1366 (2016).

82. Takechi-Haraya, Y. et al. Atomic Force Microscopic Analysis of the Effect of Lipid Composition on Liposome Membrane Rigidity. Langmuir 32, 6074–6082 (2016).

83. Li, J. et al. Self-assembled multivalent DNA nanostructures for noninvasive intracellular delivery of immunostimulatory CpG oligonucleotides. ACS Nano 5, 8783–8789 (2011).

84. Hung, Y. H., Chen, L. M. W., Yang, J. Y. & Yuan Yang, W. Spatiotemporally controlled induction of autophagy-mediated lysosome turnover. Nat. Commun. 4, 2111 (2013).

85. Liang, L., et al. Single-particle tracking and modulation of cell entry pathways of a tetrahedral DNA nanostructure in live cells. *Angew. Chem. Int. Ed*. 53, 7745–7750 (2014).

86. Balakrishnan, D., Wilkens, G. D. & Heddle, J. G. Delivering DNA origami to cells. Nanomed. 14, 911–925 (2019).

87. Macia, E. et al. Dynasore, a cell-permeable inhibitor of dynamin. Dev. Cell 10, 839–850 (2006).

88. Kirchhausen, T., Macia, E. & Pelish, H. E. Use of dynasore, the small molecule inhibitor of dynamin, in the regulation of endocytosis. in Methods in Enzymology vol. 438 77–93 (2008).

89. Ferguson, S. M. & De Camilli, P. Dynamin, a membrane-remodelling GTPase. Nat. Rev. Mol. Cell Biol. 13, 75–88 (2012).

90. Hivare, P., Rajwar, A., Gupta, S. & Bhatia, D. Spatiotemporal dynamics of endocytic pathways adapted by small DNA nanocages in model neuroblastoma cell-derived differentiated neurons. ACS Appl. Bio Mater. 4, 3350–3359 (2021).

91. Kafle, B. P. Molecular luminescence spectroscopy. in Chemical analysis and material characterization by spectrophotometry 269–296 (Elsevier, Cambridge, 2020).

92. Pazos, E., Vázquez, O., Mascareñas, J. L. & Vázquez, M. E. Peptide-based fluorescent biosensors. Chem. Soc. Rev. 38, 3348–3359 (2009).

93. Santinho, A., Carpentier, M., Lopes Sampaio, J., Omrane, M. & Thiam, A. R. Giant organelle vesicles to uncover intracellular membrane mechanics and plasticity. Nat. Commun. 15, 3767 (2024).

94. López-Duarte, I., Truc Vu, T., Izquierdo, M. A., Bull, J. A. & Kuimova, M. K. A molecular rotor for measuring viscosity in plasma membranes of live cells. Chem. Commun. 50, 5282–5284 (2014).

95. Sharpe, H. J., Stevens, T. J. & Munro, S. A comprehensive comparison of transmembrane domains reveals organelle-specific properties. Cell 142, 158–169 (2010).

96. Antonny, B., Vanni, S., Shindou, H. & Ferreira, T. From zero to six double bonds: Phospholipid unsaturation and organelle function. Trends Cell Biol. 25, 427–436 (2015).

97. Harayama, T. & Riezman, H. Understanding the diversity of membrane lipid composition. Nat. Rev. Mol. Cell Biol. 19, 281–296 (2018).

98. Niko, Y., Didier, P., Mely, Y., Konishi, G. I. & Klymchenko, A. S. Bright and photostable push-pull pyrene dye visualizes lipid order variation between plasma and intracellular membranes. Sci. Rep. 6, 18870 (2016).

99. Guolla, L., Bertrand, M., Haase, K. & Pelling, A. E. Force transduction and strain dynamics in actin stress fibres in response to nanonewton forces. J. Cell Sci. 125, 603–613 (2012).

100. Yap, B. & Kamm, R. D. Cytoskeletal remodeling and cellular activation during deformation of neutrophils into narrow channels. J. Appl. Physiol. 99, 2323–2330 (2005).

101. Centola, M. et al. A rhythmically pulsing leaf-spring DNA-origami nanoengine that drives a passive follower. Nat. Nanotechnol. 19, 226–236 (2024).

102. Hu, Y. et al. Quantifying T cell receptor mechanics at membrane junctions using DNA origami tension sensors. Nat. Nanotechnol. 19, 1674–1685 (2024).

103. Reinhardt, S. C. M. et al. Ångström-resolution fluorescence microscopy. Nature 617, 711–716 (2023).

104. Huang, J. et al. A modular DNA origami nanocompartment for engineering a cell-free, protein unfolding and degradation pathway. Nat. Nanotechnol. 19, 1521– 1531 (2024).

105. Saminathan, A. et al. A DNA-based voltmeter for organelles. Nat. Nanotechnol. 16, 96–103 (2021).

106. Cui, C. et al. A lysosome-targeted DNA nanodevice selectively targets macrophages to attenuate tumours. Nat. Nanotechnol. 16, 1394–1402 (2021).

107. Modi, S. et al. A DNA nanomachine that maps spatial and temporal pH changes inside living cells. Nat. Nanotechnol. 4, 325–330 (2009).

108. Zou, J. et al. A DNA nanodevice for mapping sodium at single-organelle resolution. Nat. Biotechnol. 42, 1075–1083 (2024).

109. Hu, Y. et al. DNA-based ForceChrono probes for deciphering single-molecule force dynamics in living cells. Cell 187, 3445–3459.e15 (2024).

110. Mercier, V. et al. Endosomal membrane tension regulates ESCRT-III-dependent intra-lumenal vesicle formation. Nat. Cell Biol. 22, 947–959 (2020).

111. Boucrot, E. & Kirchhausen, T. Endosomal recycling controls plasma membrane area during mitosis. Proc. Natl. Acad. Sci. 104, 7939–7944 (2007).

112. Lafaurie-Janvore, J. et al. ESCRT-III assembly and cytokinetic abscission are induced by tension release in the intercellular bridge. Science 339, 1625–1629 (2013).

113. De Belly, H., Paluch, E. K. & Chalut, K. J. Interplay between mechanics and signalling in regulating cell fate. Nat. Rev. Mol. Cell Biol. 23, 465–480 (2022).

114. Gudipaty, S. A. et al. Mechanical stretch triggers rapid epithelial cell division through Piezo1. Nature 543, 118–121 (2017).

115. Hu, D., Dong, Z., Li, B., Lu, F. & Li, Y. Mechanical force directs proliferation and differentiation of stem cells. Tissue Eng. Part B Rev. 29, 141–150 (2023).

116. Nguyen, H. L. et al. Elastic moduli of normal and cancer cell membranes revealed by molecular dynamics simulations. Phys. Chem. Chem. Phys. 24, 6225– 6237 (2022).

117. Corbin, E. A., Kong, F., Lim, C. T., King, W. P. & Bashir, R. Biophysical properties of human breast cancer cells measured using silicon MEMS resonators and atomic force microscopy. Lab. Chip 15, 839–847 (2015).

118. Firmin, J. et al. Mechanics of human embryo compaction. Nature 629, 646– 651 (2024).

119. Eisenstein, M. Push and pull: how to measure the forces that sculpt embryos. Nature 630, 780–782 (2024).

120. Suma, A. et al. TacoxDNA: A user-friendly web server for simulations of complex DNA structures, from single strands to origami. J. Comput. Chem. 40, 2586–2595 (2019).

121. Poppleton, E. et al. Design, optimization and analysis of large DNA and RNA nanostructures through interactive visualization, editing and molecular simulation. Nucleic Acids Res. 48, e72 (2020).

122. Snodin, B. E. K. et al. Introducing improved structural properties and salt dependence into a coarse-grained model of DNA. J. Chem. Phys. 142, 234901 (2015).

123. Matthies, M., Sample, M. & Sulc, P. Coarse-Grained Simulations of DNA and RNA Systems with oxDNA and oxRNA Models: Introductory Tutorial. (2023). doi:10.48550/arXiv.2308.01455.

124. Wickham, H. *Ggplot2: Elegant Graphics for Data Analysis*. (Springer, New York, NY, 2009). doi:10.1007/978-0-387-98141-3.

125. Griffo, A. et al. Mechanics of biomimetic free-standing lipid membranes: insights into the elasticity of complex lipid compositions. RSC Adv. 14, 13044– 13052 (2024).

