## Supporting Information for "Molecular machines measure the nanomechanics of biological membranes"

### 1. Supplementary Figures

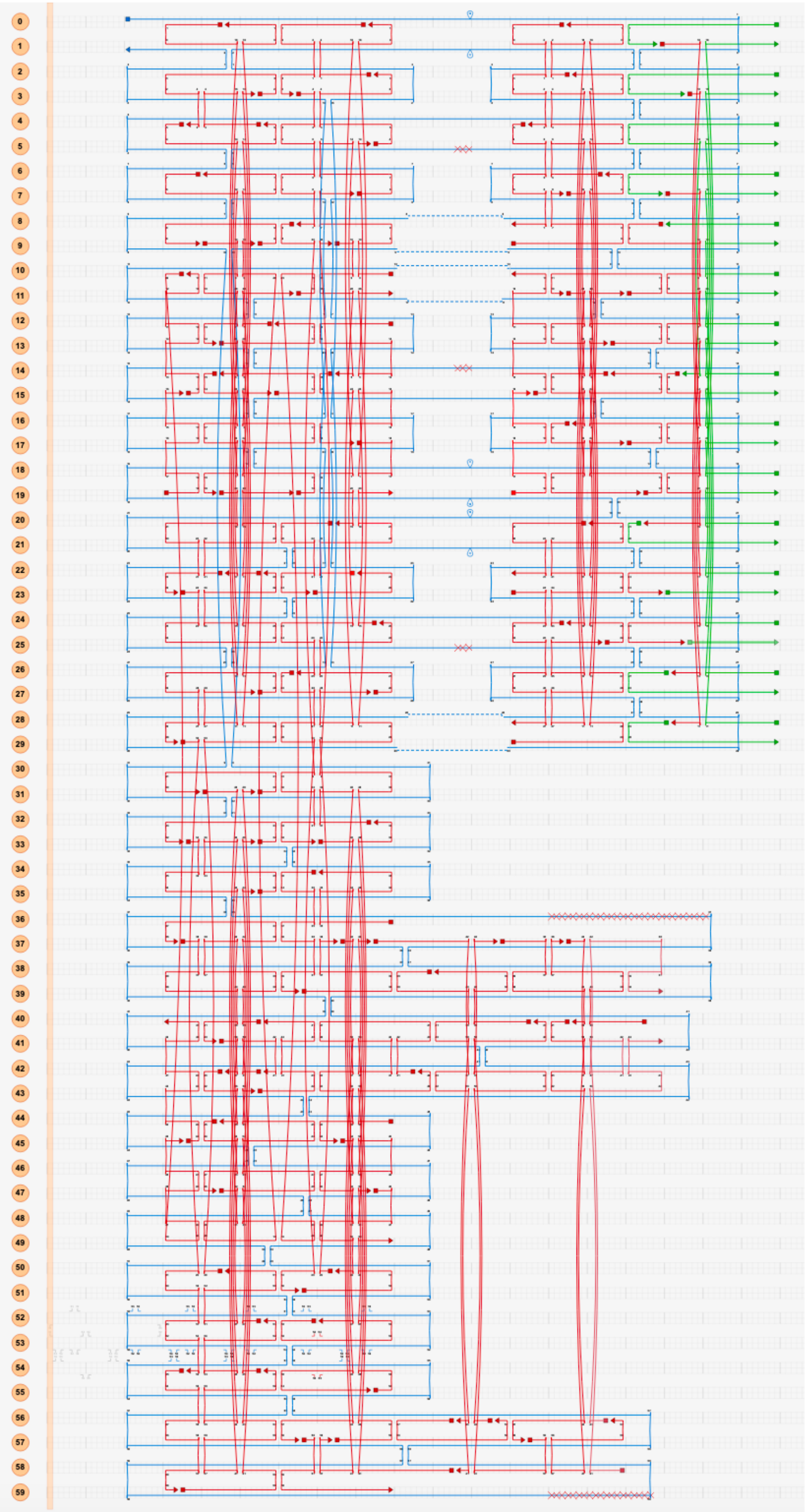

**Supplementary Figure 1: 2D connectivity map of the MM designed with CaDNAno.** The scaffold is depicted in blue, staple strands are indicated in red, and cholesterol anchoring strands in green. 5' and 3' termini of the DNA strands are represented by a square and triangle, respectively. The duplexes are numbered on the left side.

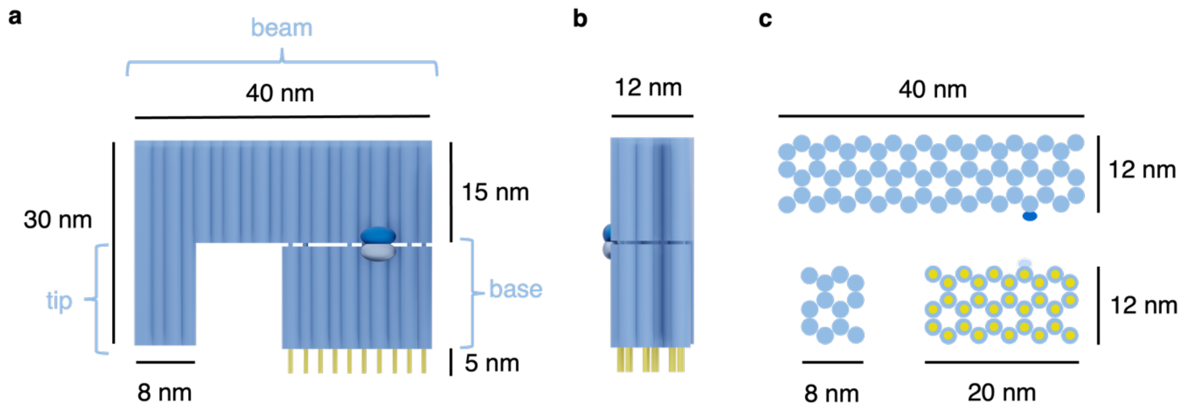

**Supplementary Figure 2. 3D model of the MM and its dimensions.** The 3D model was generated with Blender and is shown in views from the **a)** front **b)** side **c)** top and bottom. MM consists of a base, a beam, and a tip. The blue cylinders represent DNA duplexes, which are interconnected by single strands (not shown). Yellow cylinders indicate cholesterol anchors for membrane binding of MM. The dark blue and the light blue dots annotate the location for the incorporated Cy5 and Cy3 fluorophores, respectively.

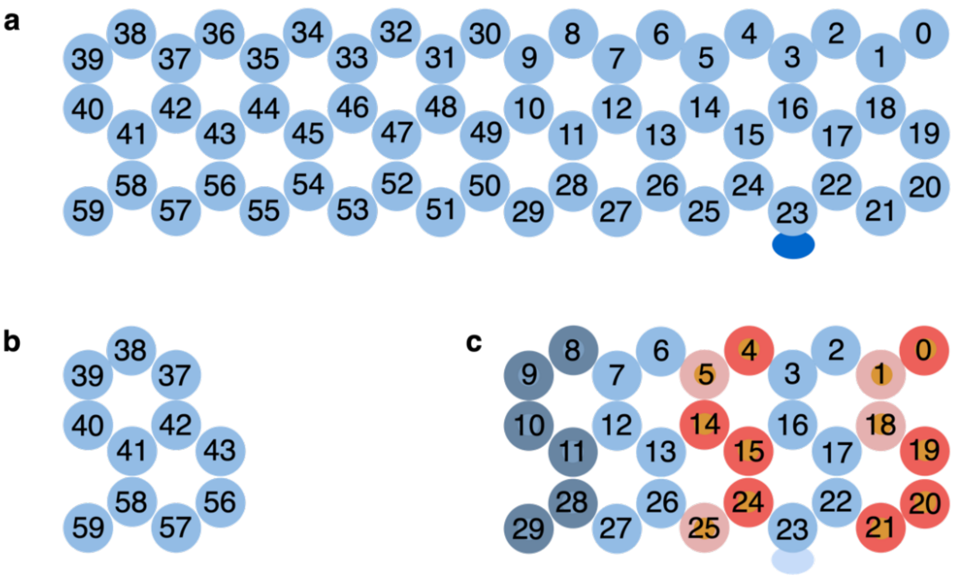

**Supplementary Figure 3. 2D duplex map of MM, its tip and base with numbered component duplexes in top view.** Each circle represents a double helix within the MM. The numbering of duplexes is the same as in Supplementary Figure 1. **a)** Top view of the beam of MM. The dark blue oval on duplex 23 indicates the Cy3 dye. **b)** Horizontally cut section of the MM tip. **c)** Top view of the MM base. The red and light red circles indicate the locations of the hairpin and single-stranded actuator sequences, respectively, and grey-blue shows the position of the single-stranded hinge sequences, respectively. Actuator and hinge sequences connect the beam and the base (not shown). Actuator sequences in positions 4, 5,14, 15, 24, and 25 are shorter in length than those of position 0, 1, 18, 19, 20, and 21. The former hybridize with short opening strands 7-12 while the latter bind to long opening strands 1-6 (Supplementary Table 5).

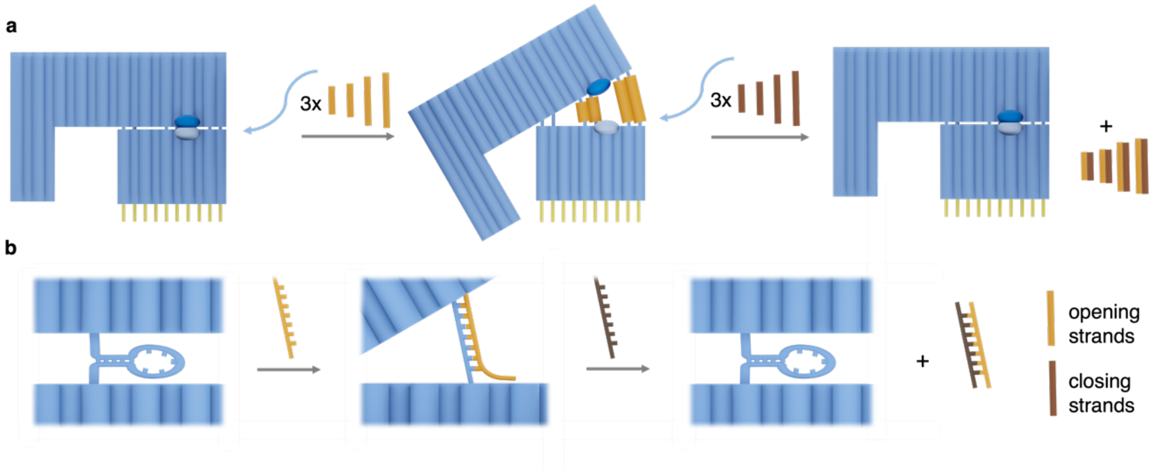

**Supplementary Figure 4. Scheme on the reversible opening and closing of MM. a)** The addition of 12 oligonucleotide opening strands (orange) opens MM while the addition of 12 closing oligonucleotides (brown) closes the MM to yield side product duplexes. **b)** Detailed mechanism of the DNA-triggered opening and closure of MM exemplified by a single actuator hairpin located between beam (top) and base (bottom). The single-stranded opening strand binds to the dumbbell-like hairpin of complementary sequence to unzip the hairpin and form an energetically favourable duplex which pushes the beam and base apart to open MM. To revert the opening of MM to the closed state, a closing strand binds to a single-stranded toehold section of the MM-bound opening strand to unzip it from the hairpin sequence. This reforms the hairpin and reverts MM to the closed conformation. A new duplex between complementary opening and closing strands is formed as a side product.

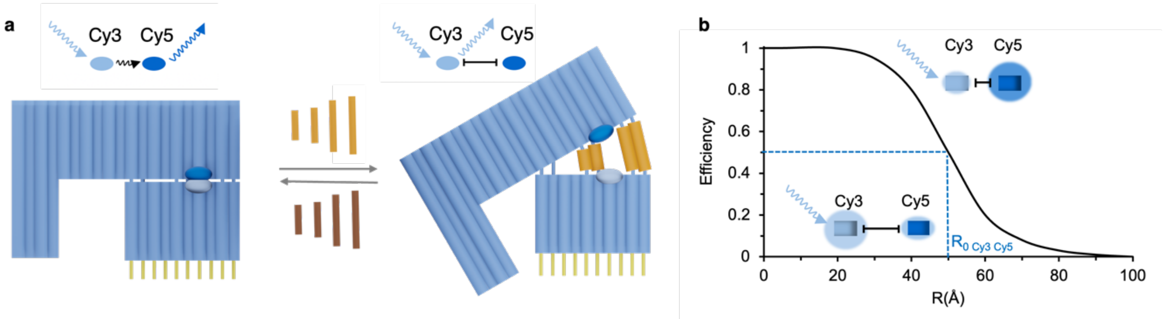

**Supplementary Figure 5. Triggered reversible opening of the MM is tracked by fluorescence resonance energy transfer (FRET). a)** Scheme of MM carrying FRET donor dye Cy3 (light blue) and acceptor dye Cy5 (dark blue) at the base and beam, respectively as distance markers. MM opening and closing by the addition of opening and closing strands alters the distance between Cy3 and Cy5 and causes FRET-induced changes in fluorescence emission. In the closed conformation, both fluorophores are within proximity, and excitation of Cy3 leads to FRET and Cy5 emission. In the open state, the two fluorophores are apart and FRET between Cy3 and Cy5 is weakened as the dye distance exceeds the Förster distance $R_0$. **b)** Plot displaying the distance dependency of the FRET efficiency for Cy3 and Cy5. An increase in the distance between the two fluorophores leads to a decrease in FRET efficiency. The Förster distance $R_0$ is defined as the distance between two fluorophores to allow 50% FRET efficiency and is 5 nm for the FRET pair Cy3 Cy5. Illustration adapted from Roy et al.[1]

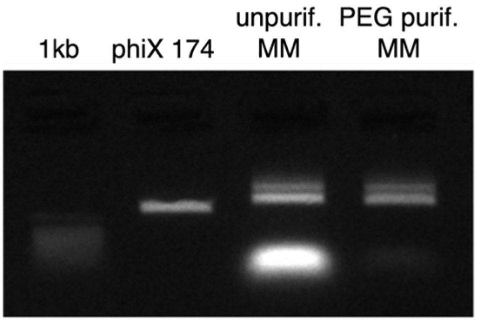

**Supplementary Figure 6. Gel electrophoretic analysis on the formation and purification of MM.** 1% agarose gel in 0.5x TAE, 10 mM $MgCl_2$ at 65 V for 1 h separating PhiX174 scaffold, unpurified folded MM and PEG-purified MM. The upward mobility shift for the assembled MM band relative to the scaffold band indicates successful assembly; a higher-migrating band represent a dimeric MM. The fast migrating, intense band of small staple strands which had been added in 6x excess for MM assembly, were removed by precipitation with PEG, as confirmed by the disappearance of the gel band for staple strands.

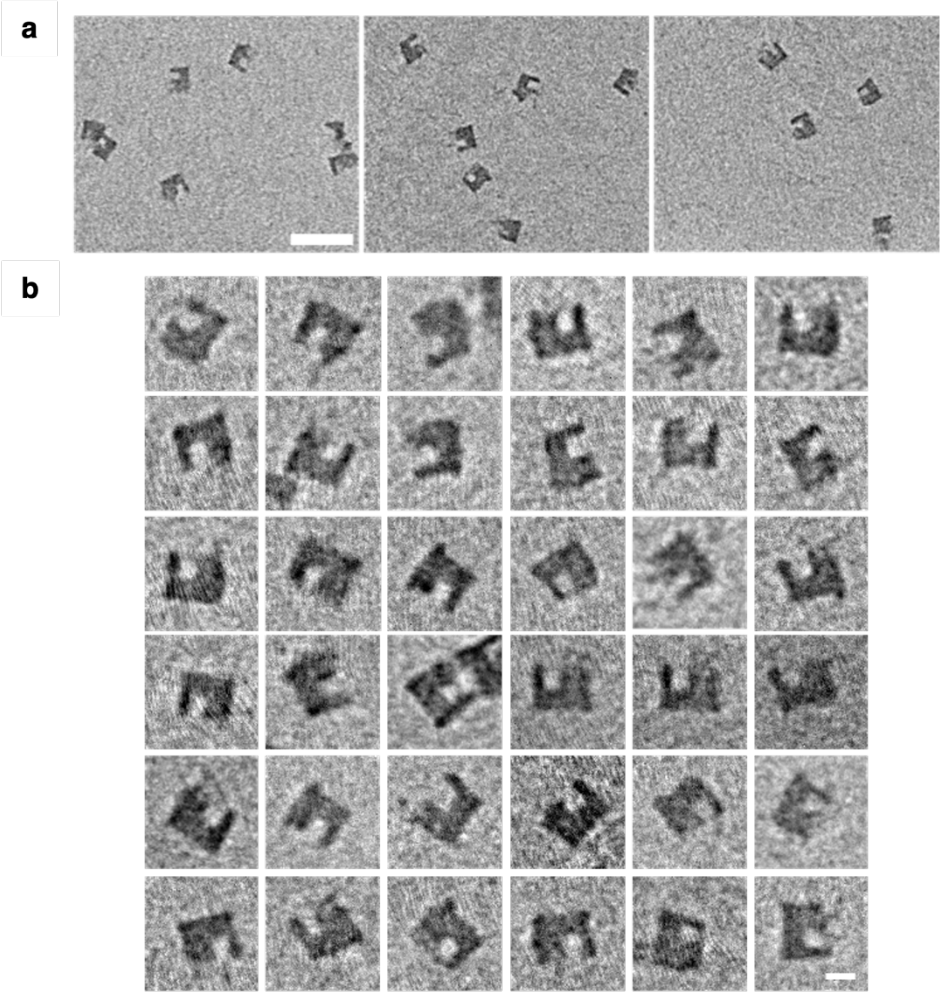

**Supplementary Figure 7. Transmission electron microscopy images of negatively stained MM in the closed state. a)** Low-magnification view. Scale bar, 100 nm. Of the 200 particles analysed, 83% were properly folded monomers. **b)** High-magnification view of representative properly formed particles selected for tilt-angle analysis. Scale bar, 20 nm.

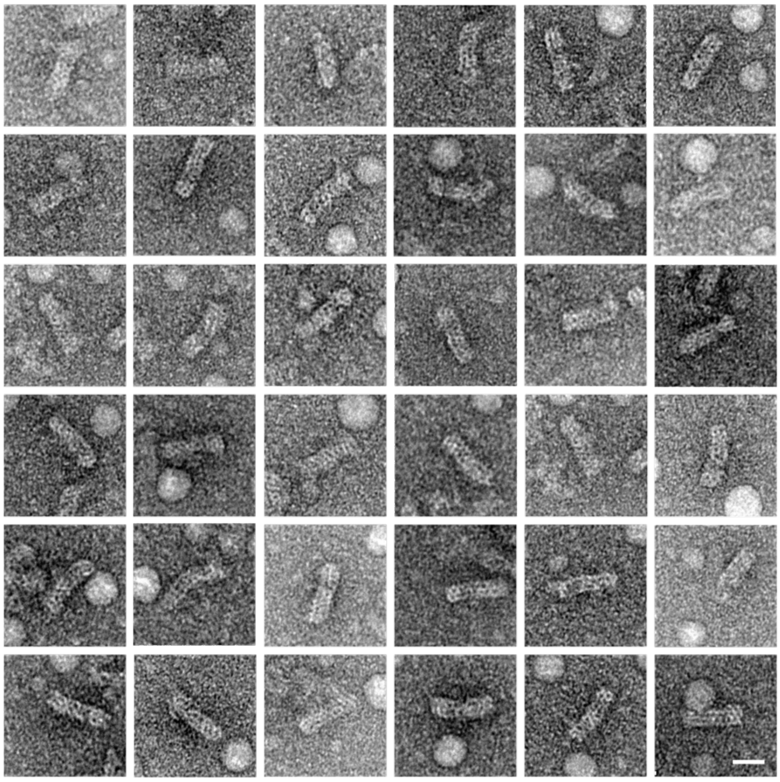

**Supplementary Figure 8. TEM analysis of negatively stained MM in the closed conformation in top view.** Scalebar, 20 nm.

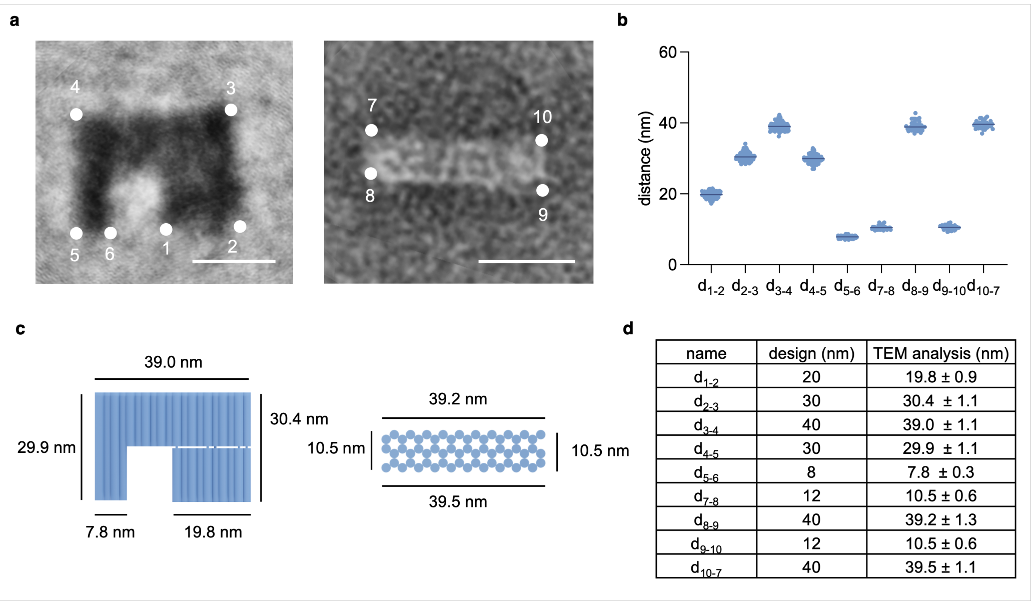

| name | design (nm) | TEM analysis (nm) |
|---|---|---|
| $d_{1\text{-}2}$ | 20 | 19.8 ± 0.9 |
| $d_{2\text{-}3}$ | 30 | 30.4 ± 1.1 |
| $d_{3\text{-}4}$ | 40 | 39.0 ± 1.1 |
| $d_{4\text{-}5}$ | 30 | 29.9 ± 1.1 |
| $d_{5\text{-}6}$ | 8 | 7.8 ± 0.3 |
| $d_{7\text{-}8}$ | 12 | 10.5 ± 0.6 |
| $d_{8\text{-}9}$ | 40 | 39.2 ± 1.3 |
| $d_{9\text{-}10}$ | 12 | 10.5 ± 0.6 |
| $d_{10\text{-}7}$ | 40 | 39.5 ± 1.1 |

**Supplementary Figure 9. Dimensions of MM obtained by analysis of TEM images**. **a)** Mean TEM images after MM alignment from the front and top view with marked points for dimension measurements. **b)** Plot displaying the measured dimensions for MM. Lowercase numbers indicate the two points used for distance measurements, n = 128 for side view, n = 36 for top view. **c)** Model of MM in front and top view with the mean dimensions obtained by TEM analysis. **d)** Table summarising nominal dimensions from the MM design and dimensions obtained from TEM analysis (n = 128 for side view, n = 20 for top view; ± SD).

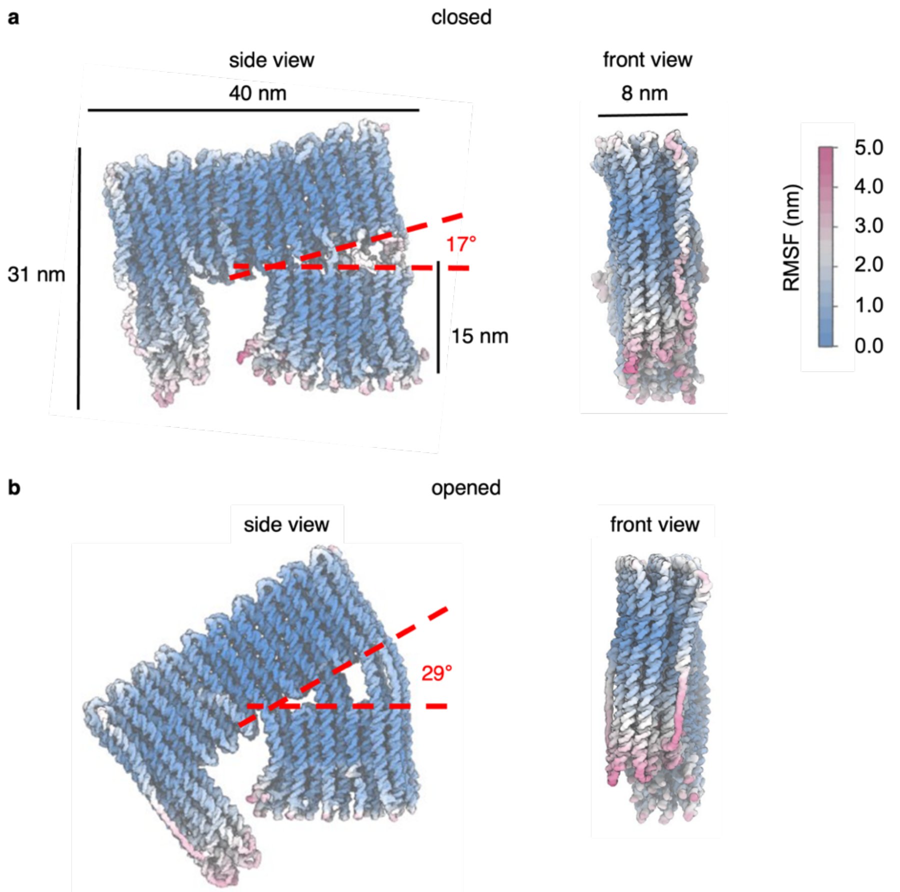

**Supplementary Figure 10. oxDNA simulations of MM. a)** Closed and **b)** opened MM shown in front (left) and side view (right) with oxDNA-derived dimensions, the angle between the beam and base is annotated in red. The colours in the MM structures represent the RMSF noise as indicated by the scale.

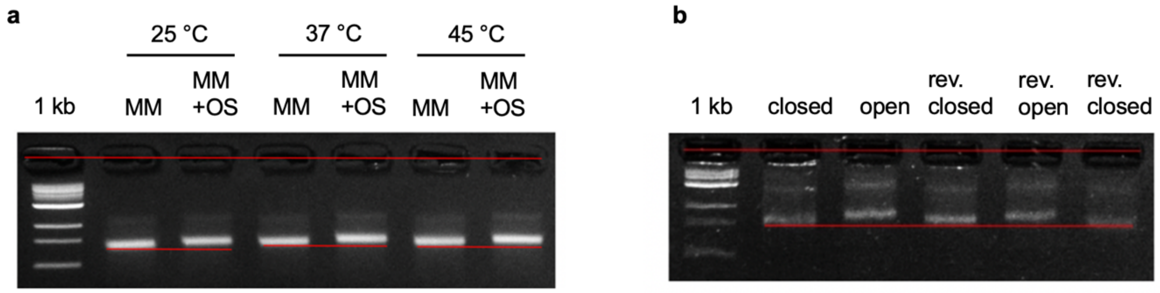

**Supplementary Figure 11. Gel electrophoretic analysis of MM opening at various temperatures and reversible MM opening and closing. a)** 2% Agarose gel run in 0.5 x TBE, 10 mM $MgCl_2$ for 2.5 h at 65 V confirms the opening of MM at temperatures from 25 °C to 45 °C, as indicated by the gel band shift occurring for closed MM when mixed with opening strands (OS). **b)** 2.5% Agarose gel for 3 h at 65 V on the reversible opening and closing of MM in 1 x PBS as indicated by repeated upward and downward gel shifts. Some MM aggregation is promoted at high agarose gel concentration. A red line parallel to the wells aids as a visual guide to highlight the gel shift. A 1 kb DNA ladder at the left is a size reference.

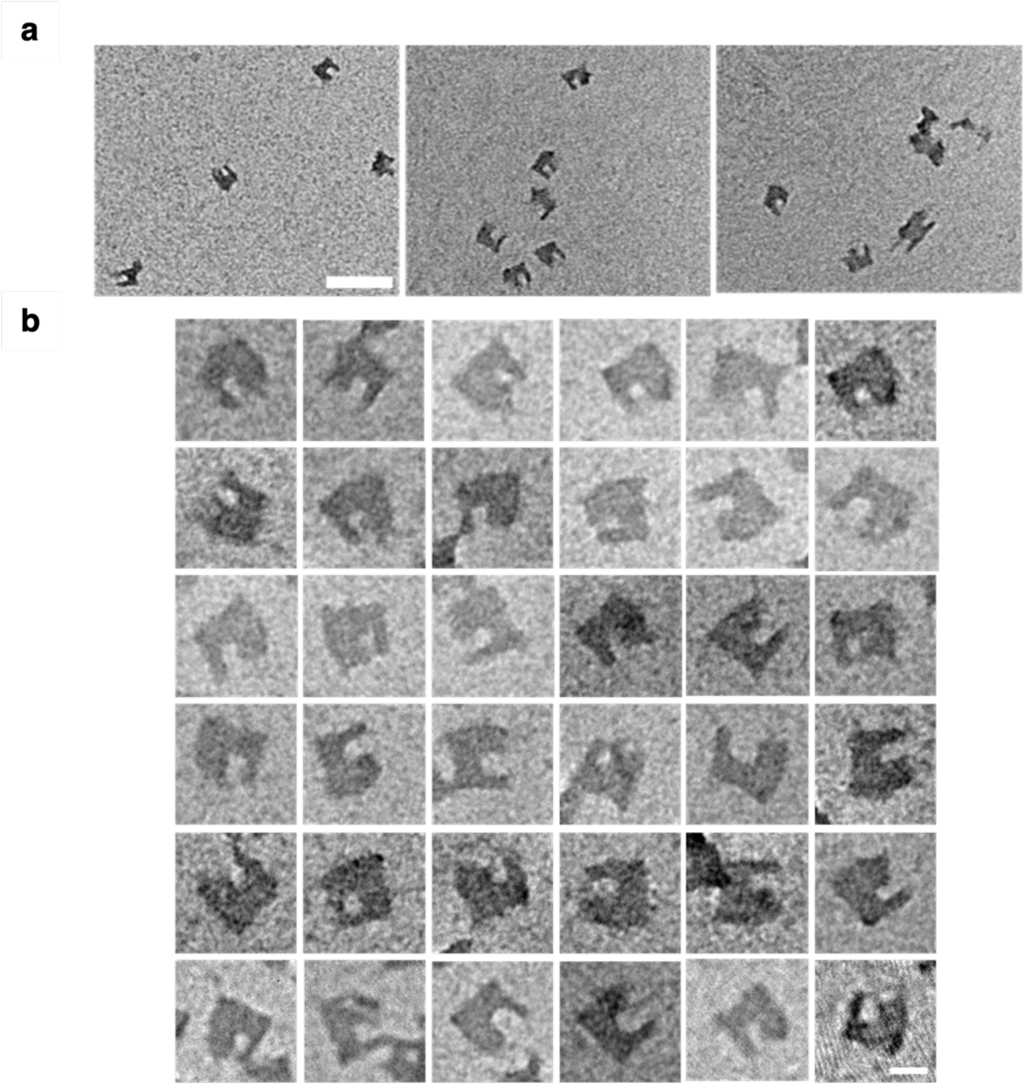

**Supplementary Figure 12. TEM images of negatively stained MM in the open state. a)** Low-magnification view. Scale bar, 100 nm. Of the 200 particles analysed, 81% were properly folded monomers. **b)** High-magnification view of representative properly formed particles selected for tilt-angle analysis. Scale bar, 20 nm.

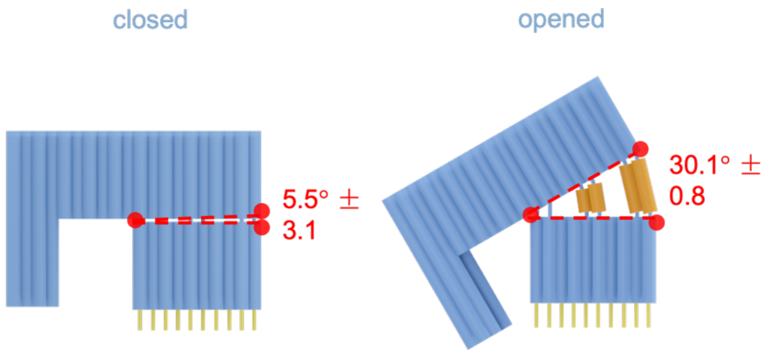

**Supplementary Figure 13. Analysis of MM opening angle from TEM images.** Front view of the 3D model of MM in closed and opened confirmations, indicating the corresponding opening angles (red) as obtained by TEM analysis. Corresponding scatter plot of the opening angles between base and beam obtained from TEM images of MM in closed and opened state is in Figure 2d of the main manuscript.

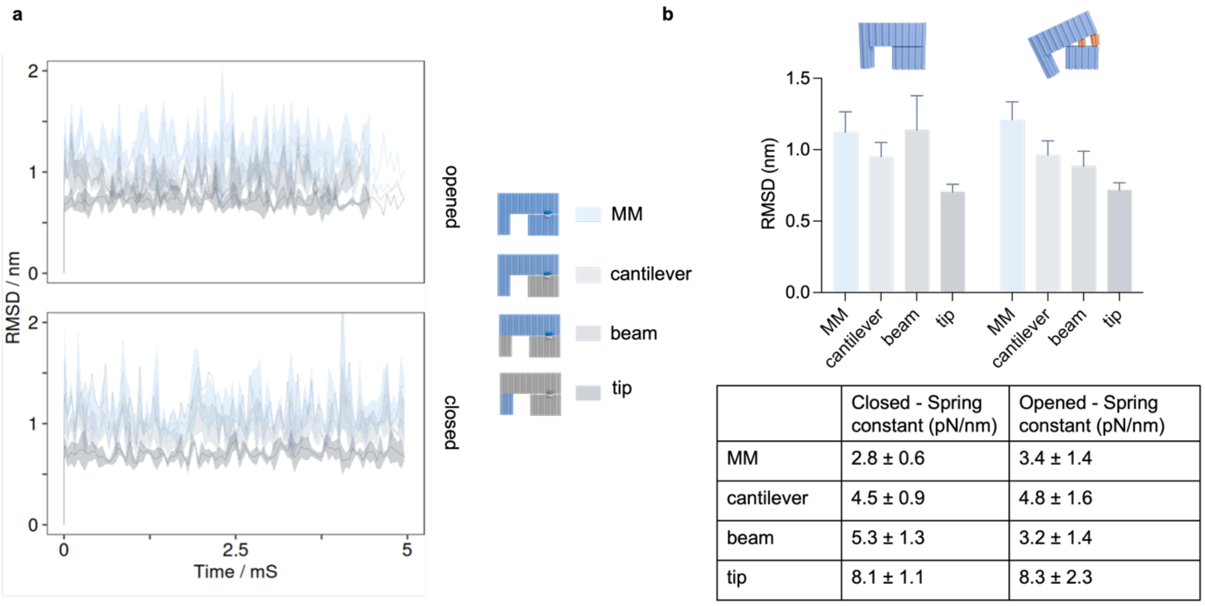

| | Closed - Spring constant (pN/nm) | Opened - Spring constant (pN/nm) |
|---|---|---|
| MM | 2.8 ± 0.6 | 3.4 ± 1.4 |
| cantilever | 4.5 ± 0.9 | 4.8 ± 1.6 |
| beam | 5.3 ± 1.3 | 3.2 ± 1.4 |
| tip | 8.1 ± 1.1 | 8.3 ± 2.3 |

**Supplementary Figure 14. OxDNA simulations of MM yield the time-dependent root mean squared displacement (RMSD) to calculate the spring constant of MM's cantilever**. **a)** Each kinetic RMSD trace shows the average of 3 independent oxDNA simulations and the coloured ribbon shows the standard deviation. RMSD data were analyzed for MM, the cantilever, the beam and the tip. **b)** Mean values for RMSD calculations from a) as bar diagram (top) and as table with spring constant (k) which were derived from the RMSD using the formula $k = \frac{k_b t}{x^2}$ where $k_b$ is the Boltzmann constant, $t$ is the temperature and $x$ is the RMSD. The error of the spring constant was calculated using Taylor series expansion.

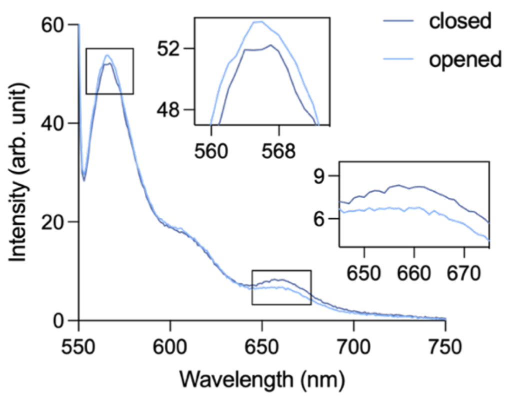

**Supplementary Figure 15. Fluorescent emission spectra of Cy3/Cy5-labelled MM with FRET-dependent signatures for the open and closed state.** Spectra of closed MM (brown) and opened MM (orange) using Cy3 excitation at 540 nm show emission peaks at 574 nm (Cy3) and 664 nm (Cy5). Increased emission at 664 nm is a signature for the closed MM state while increased emission at 574 nm is characteristic for the open MM state.

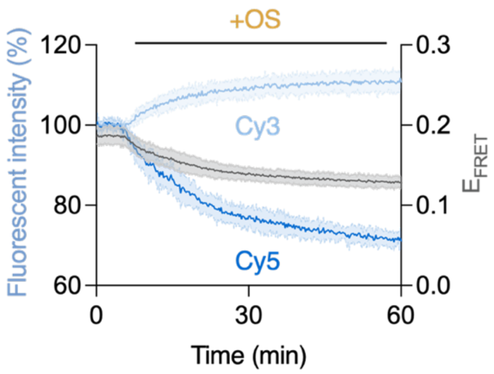

**Supplementary Figure 16. Mean kinetic traces of relative Cy3 and Cy5 fluorescence changes, and FRET efficiency ($E_{FRET}$) changes upon addition of OS .** MM opening was monitored for 60 min after addition of OS at 5 min. Traces were smoothed via 2nd order smoothing (4 neighbours) integrated within GraphPad Prism. Mean trace ± SD from n = 11. $E_{FRET}$ was calculated via $E_{FRET} = \frac{F_{Cy5}}{F_{Cy3}+F_{Cy5}}$.

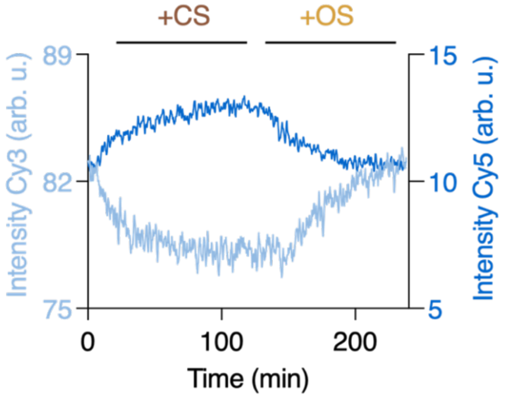

**Supplementary Figure 17. Kinetic FRET traces showing reversibility of subsequential closing and opening of MM.** Cy3 and Cy5 kinetic emission traces tracking MM closing and opening after MM assembly with OS. MM fluorescent traces were monitored over 240 min; CS was added at 5 min and CS at 130 min.

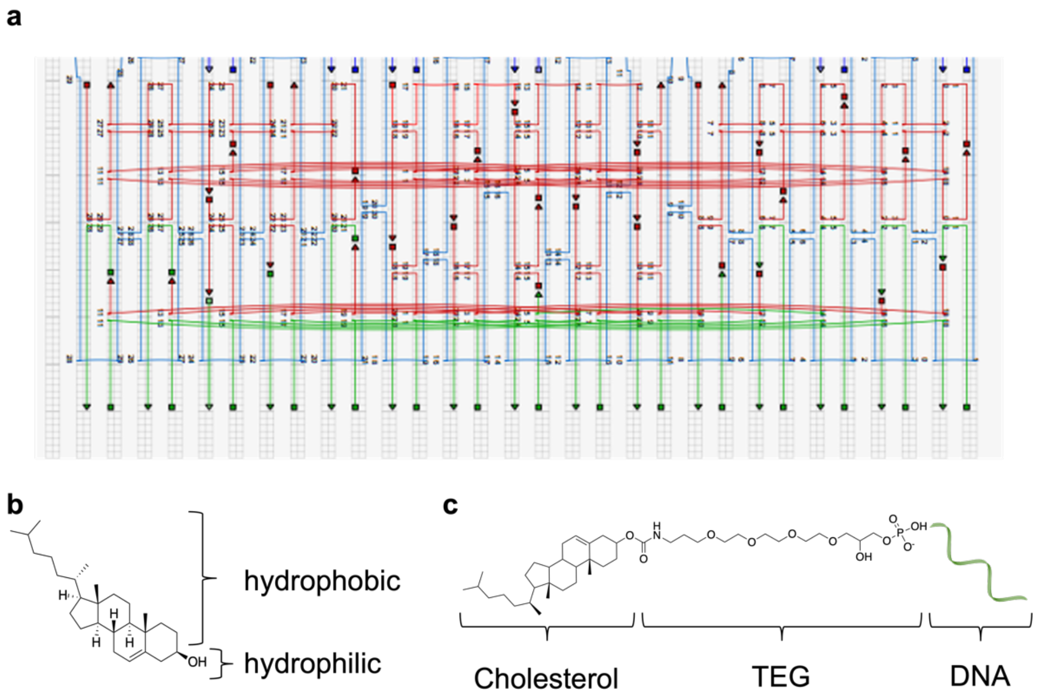

**Supplementary Figure 18. MM position and chemical structure of the cholesterol-tag used for membrane anchoring of MM. a)** Part of the 2D DNA connectivity map of MM's base unit. Blue strands represent the scaffold strand, red the staple strands, and green the lipid anchoring strands. The arrows indicate the 3' and square the 5' terminus of staple and anchoring strands. **b)** Chemical diagram of cholesterol with hydrophobic (red) and hydrophilic (blue) parts. **c)** Lipid anchoring strand comprising cholesterol attached via a flexible tetra(ethylene glycol) (TEG) linker to the DNA sequence.

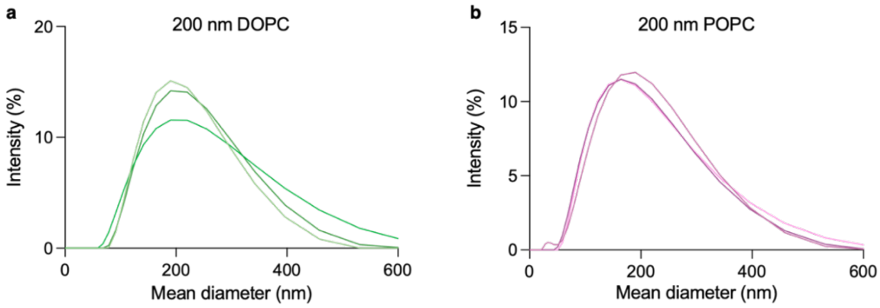

**Supplementary Figure 19. DLS analysis on the size distribution of DOPC and POPC small unilamellar vesicles extruded at 200 nm**.

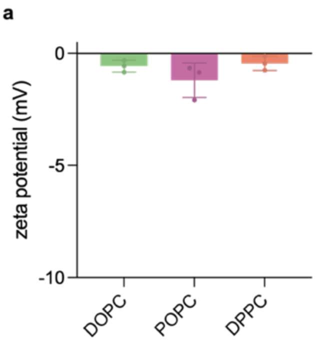

**b**

| lipid | zeta potential (mV) |
|---|---|
| DOPC | -0.57 ± 0.26 |
| POPC | -1.20 ± 0.77 |
| DPPC | -0.45 ± 0.32 |

**Supplementary Figure 20. Zeta potential for different lipid vesicles. a)** Zeta potential for DOPC, POPC and DPPC vesicles (200 nm, 100 μM lipid concentration) at pH 8, n=3. **b)** Table of zeta potential values ± SD, n=3.

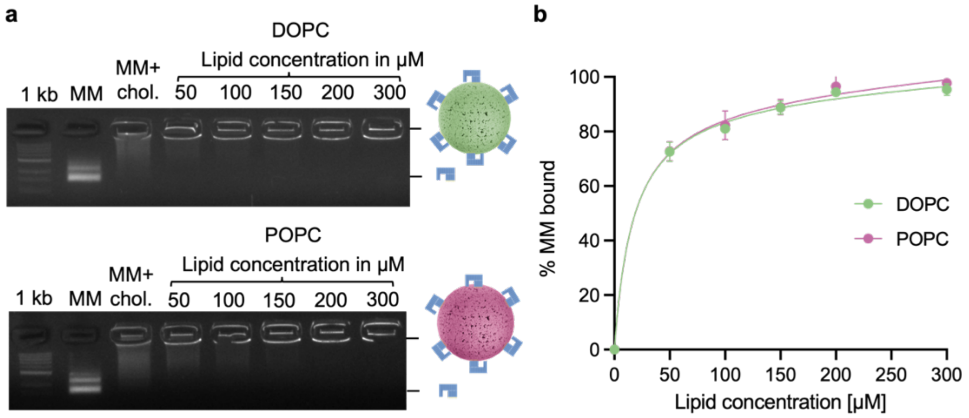

**Supplementary Figure 21. Gel electrophoretic analysis of MM binding to unilamellar vesicles. a)** 1% Agarose gel (0.5 x TAE 10 mM $MgCl_2$) of MM without and with cholesterol-membrane anchors; MM+chol (2.5 μM) was incubated with small unilamellar vesicles (SUVs, average diameter 200 nm) made with 50 to 300 μM DOPC or POPC final concentration. The gel band upshift of MM+chol is caused by the interaction of cholesterol tags with the gel matrix while the upshift of MM+chol incubated with SUVs is caused by successful MM binding to SUVs which are too large to migrate into the gel. **b)** Plot showing MM binding extent in dependence of SUVs concentration measured as lipid concentration. The data summarise the binding gel binding analysis as in panel a (n = 3). Fitting the graph with a Langmuir isotherm yielded $K_d$ values for MM binding of 22.0 ± 2.2 μM for DOPC and 24.0 ± 3.0 μM for POPC vesicles. A 1 kb DNA ladder was included as a reference.

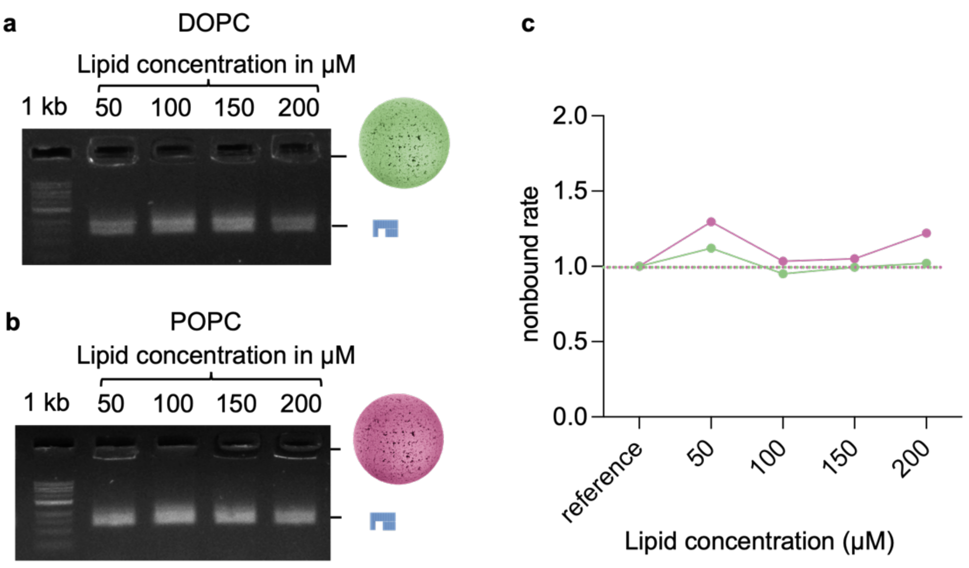

**Supplementary Figure 22. Gel electrophoretic analysis showing no membrane binding of non-cholesterol tagged MM. a)** 1% Agarose gel (0.5 x TAE 10 mM $MgCl_2$) of MM without cholesterol-membrane anchors incubated with SUVs of 200 nm average diameter formed from 50 to 300 µM DOPC or **b)** POPC. The presence of the gelectrophoretic MM band upon addition of up to 200 µM lipid concentration confirms negligible nonspecific binding of non-cholesterol MM to membrane vesicles. **c)** Plot quantifying minimal binding of MM to SUV. Mean-pixel intensities of the wells were measured via ImageJ and compared to a reference well without MM, set as reference to value 1.

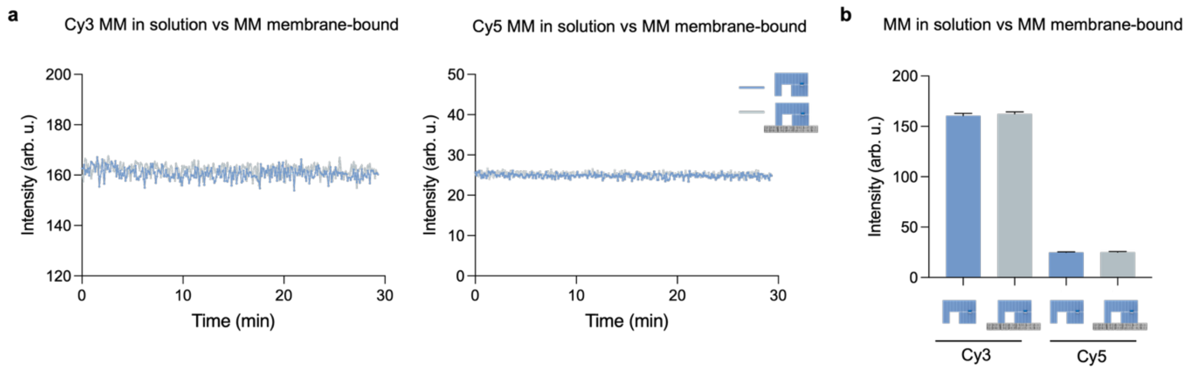

**Supplementary Figure 23. Membrane binding does not affect MM fluorescence emission. a)** Kinetic Cy3 and Cy5 emission traces of MM in solution (blue) and membrane-bound MM (grey). **b)** Mean Cy3 and Cy5 emission intensities of MM in solution and membrane-bound MM. Membrane-bound measurements were performed with DOPC vesicles (200 nm). To account for dilution upon vesicle addition, an equivalent volume of buffer was added to MM in solution.

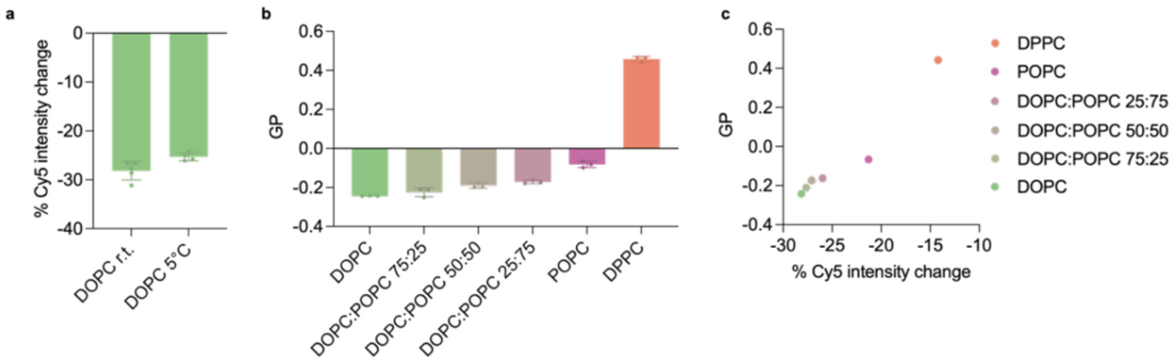

**Supplementary Figure 24. MM measurements detect temperature-induced elasticity differences and scale with membrane fluidity. a)** Percentage Cy5 emission change for MM-mediated indentation of DOPC vesicles at room temperature and 5°C. The values represent the relative change measured after 60 min incubation of MM with OS; mean ± SD (n=4). **b)** Membrane fluidity of indicated membrane vesicles as determined from membrane dye Laurdan and expressed as $GP = \frac{I_{435\pm2} - I_{490\pm2}}{I_{435\pm2} + I_{490\pm2}}$, with $I_{435\pm2}$ and $I_{435\pm2}$ being obtained from the Laurdan experiments; mean ± SD (n=3) **c)** Plot of mean GP against percentage Cy5 emission change from MM-mediated indentation.

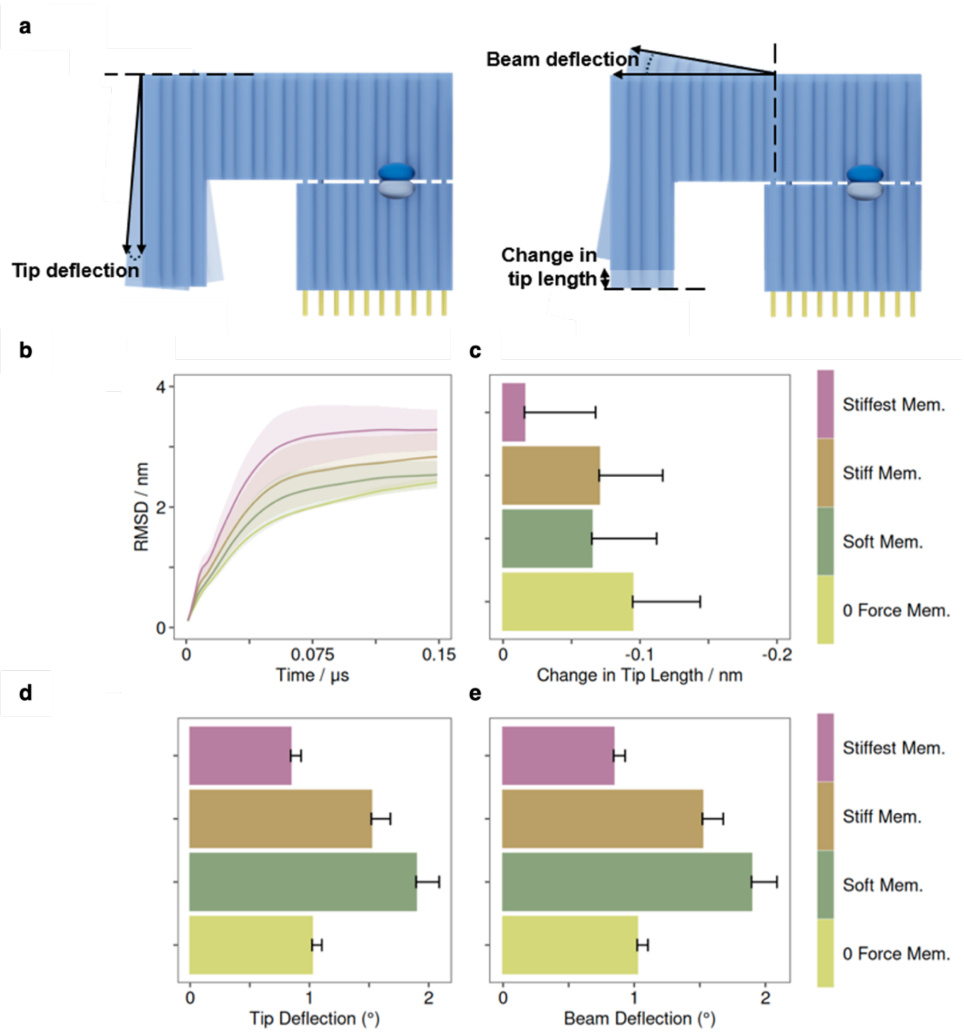

**Supplementary Figure 25. Membranes indentation does not deform the MM cantilever tip. a)** Schematic showing three potential deformation modes, involving tip or beam bending, and tip compression. **b)** Time evolution of the Root Mean Squared Deviations (RMSD) of oxDNA-simulated MM placed atop a membrane-mimicking resistive plane of various stiffness. **c)** Relative compression of the MM cantilever upon moving its tip towards the resistive plane. The compression was determined by measuring the distance between 6 oxDNA beads at the bottom of the cantilever tip versus 6 oxDNA beads at the top of cantilever, as averaged across the 6 bead pairs. The distance was corrected by any changes that occurred to the tip of the MM when simulated in the absence of the opening strands. For panel b), each trace is the average of 3 to 6 trajectories, and the standard deviation is indicated by the coloured ribbon. Data in panel c were obtained by averages averaged across the entire trajectories to account for tip-compression that occurs during actuation and membrane penetration. **d)** Tip deflection relative to the beam. Two reference vectors – along the tip longest axis and along the half of the beam closest to the tip – where calculated, to define the tip^beam angle. Tip deflection was defined as the change of this angle compared to the initial one at t = 0μs (90°). **e)** Beam deflection or flexing. Two parallel, reference vectors of equal length were defined along the beam's main axis, each spanning one half of the beam length, to define the $beam_{SectionA}$^$beam_{SectionB}$ angle. Beam deflection was defined as the change of this angle compared to the initial one at t = 0μs (180°). The "Soft" membrane has a stiffness value of 0.06 oxDNA units comparable to that of a DOPC membrane, the "Stiff" membrane has an oxDNA stiffness of 1, and the "Stiffest" membrane is set to maximum resistance.

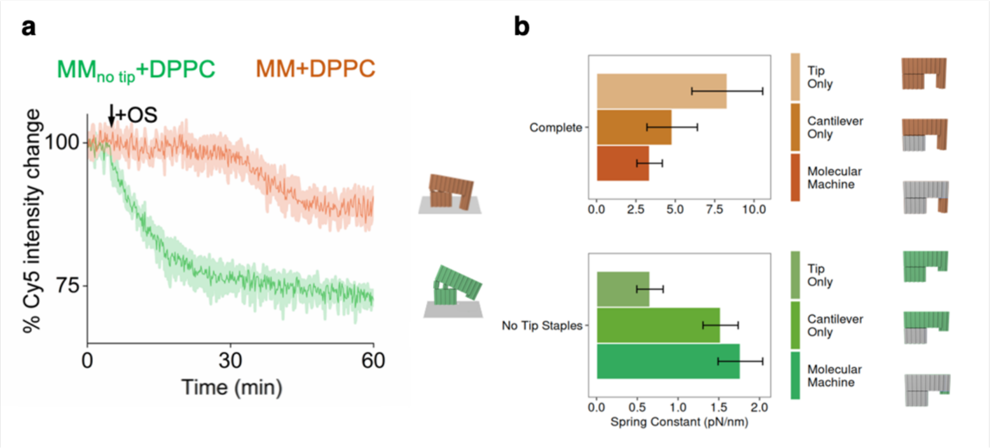

**Supplementary Figure 26. A tip-truncated MM is unable to indent membranes. a)** Fluorescence traces of $MM_{no\ tip}$ (green) and functional MM (red) bound to gel-phase DPPC membrane vesicles of high bending rigidity. $MM_{no\ tip}$ lacks the staple strands which are required for forming the rigid tip and is therefore unable to indent the membrane. $MM_{no\ tip}$ exhibits a false-positive fluorescence signal consistent with full MM opening, despite the absence of membrane indentation. In contrast, the functional MM exhibits only partial opening, as the rigid tip resists compression against the stiff gel-phase DPPC membrane. The data represent the means and standard deviations of four independent repeats. **b)** The spring constants for the tip, cantilever, and entire structure of full MM and $MM_{no\ tip}$. The spring constants were derived from oxDNA simulations and the RMSD as described in the Methods section.

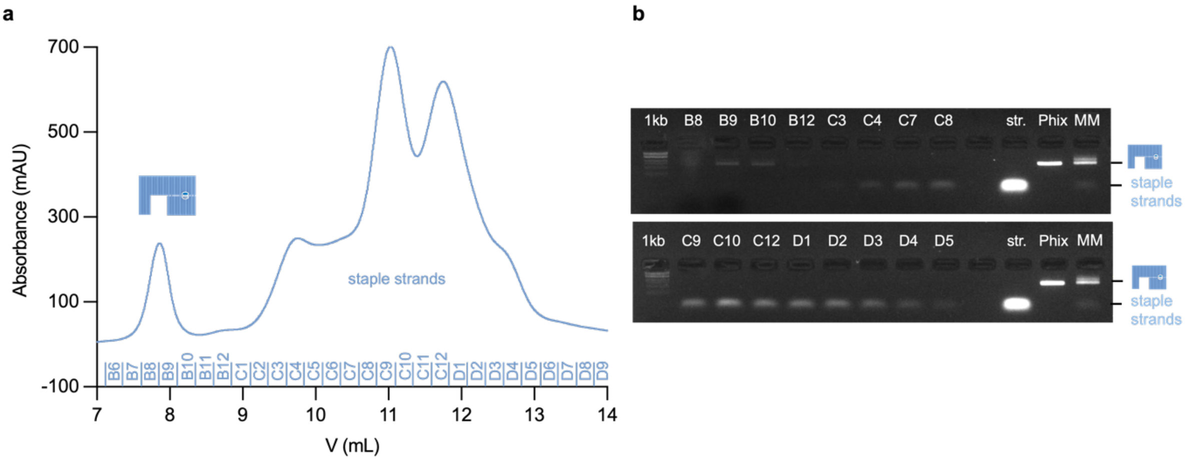

**Supplementary Figure 27. Purification of MM by size exclusion chromatography (SEC) and gel electrophoresis analysis of SEC fractions. a)** SEC elution profile for MM and component staple strands using absorption at 260 nm. The x-axis provides the names of the SEC fractions. **b)** 1% Agarose gel run for 1 h at 65 V showing the SEC fractions from panel a, and as a reference, scaffold strand PhiX174 and PEG-purified MM. Fractions B9 and B10 contain purified MM without staple strands.

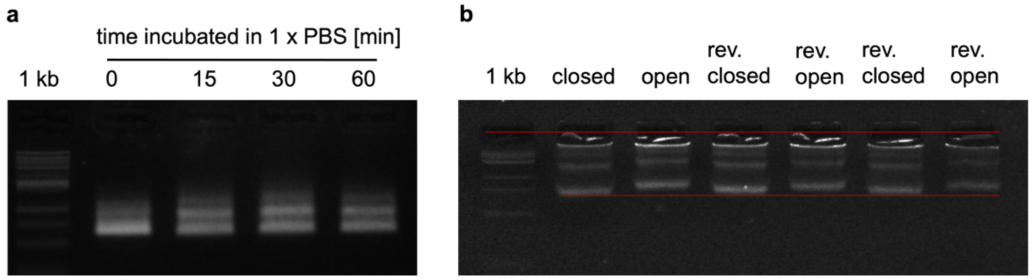

**Supplementary Figure 28. Gel electrophoresis analysis of the stability and reversible opening of MM in 1 x PBS. a)** 1% Agarose gel run (65 V, 1 h) of MM incubated in 1x PBS for up to 60 min. **b)** 2.5% Agarose gel (65 V, 3 h) on the reversible opening and closing of MM in 1 x PBS. Reversible opening and closing for two cycles are indicated by upward and downward shifts. The visible aggregation of the MM in the wells is due to a high agarose concentration.

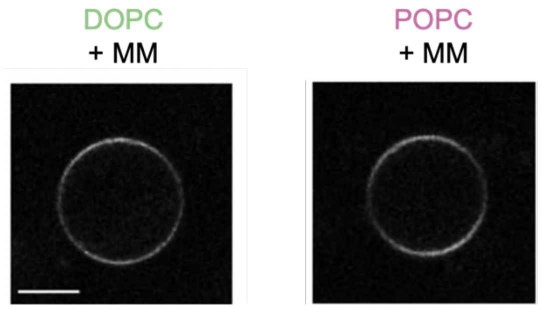

**Supplementary Figure 29. Confocal microscopy images of fluorescent-tagged MM bound to GUVs composed of DOPC and POPC imaged under the Cy5 channel.** Scale bar, 20 µm.

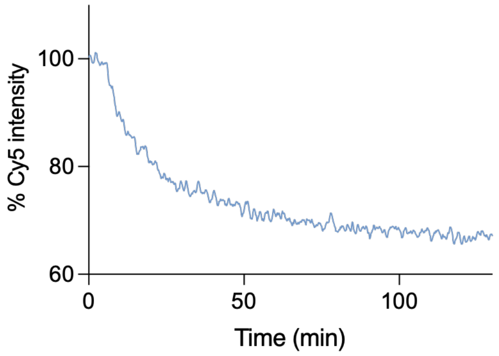

**Supplementary Figure 30. Kinetic trace of Cy5 fluorescence emission on MM opening in 1x PBS.** Normalised fluorescent Cy5 trace showing the opening of MM in 1x PBS buffer by adding to closed MM opening strands in 10x excess at 5 min., 2nd order smoothing (4 neighbours) integrated within graphpad prism.

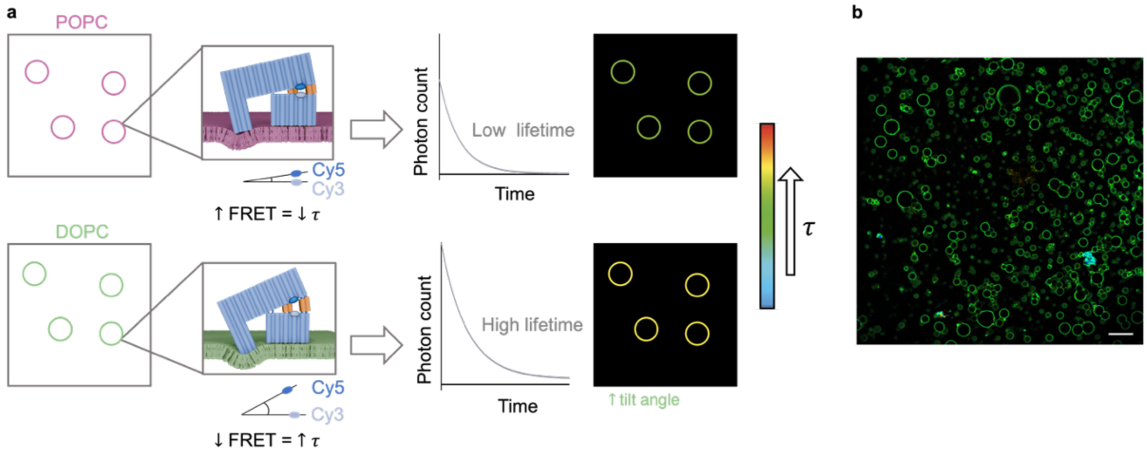

**Supplementary Figure 31. Scheme for measuring membrane elasticities by FRET and FLIM and estimation on the number of MM bound to GUVs**. **a)** POPC vesicles with stiff membrane results in a low degree of indentation and MM opening, hence proximity of the fluorophore pair Cy3/Cy5, high FRET efficiency, and low lifetime ($\tau$) which can be measured by FLIM. By contrast, DOPC vesicles with soft membrane allows for more membrane indentation, a higher degree of MM opening, greater separation of the Cy3/Cy5 pair, low FRET efficiency, and a high lifetime. The different lifetimes $\tau$ of the cells are being measured and then represented in a colourmap with red/yellow indicating high lifetimes and green/blue low lifetimes. **b)** FastFLIM microscopic image of DOPC GUVs decorated with closed MM showing a field of view of 590 x 590 µm$^2$. To the GUVs, 9*10$^{10}$ MM (30 µL, 5 nM, diluted to 120 µL) were added. Assuming complete binding and a z-section of approximately 300 nm height yields 6*10$^4$ MM per field of view. The foreground pixels account for 8% of the image leading to a final concentration of 8* 10$^5$ MM on the membranes in the field of view. This calculation underestimates the amount of bound MM, as out-of-focus MM are also binding to the membranes and MM distribution is not homogenous.

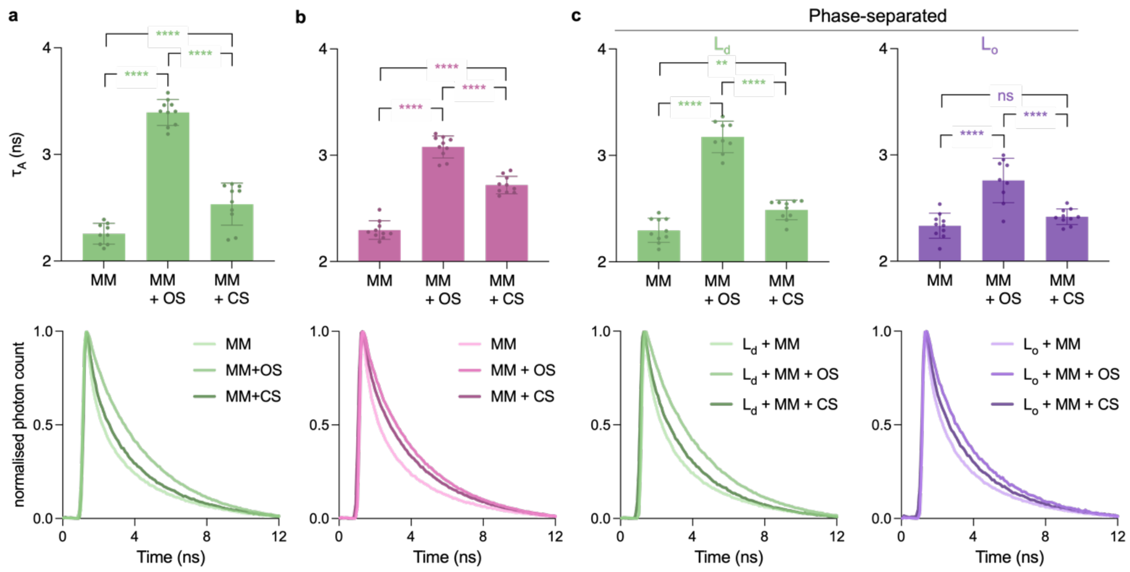

**Supplementary Figure 32. Average Cy3 lifetime and lifetime decay curves of MM bound to GUVs.** Average Cy3 lifetimes ± SD (top panel) of **a)** Cy3/Cy5-MM bound to DOPC GUVs, **b)** POPC GUVs, and **c)** biphasic GUVs with disordered DOPC and ordered DPPC CL domains, and with opening strands (+OS), and after sequential addition of closing strands (+CS). The bottom panels show the lifetime decay curves with lifetime components $\tau_1$ and $\tau_2$. The average lifetime is derived from $A_1 * \tau_1$ and $A_2 * \tau_2$ where $A_1$ and $A_2$ are weighting factors in the lifetime distribution. n = 10 GUVs analysed per condition, across three independent experiments. One-way ANOVA, Tukey's multiple comparison test, *****P < 0.0001; **P = 0.0055; *p = 0.0211; ns = not significant. *

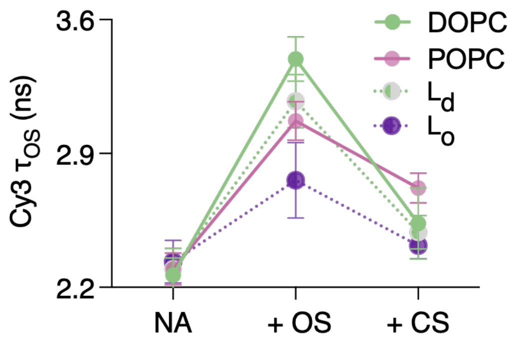

**Supplementary Figure 33**. **Cy3 lifetime changes of GUV-bound MM upon actuation reveal differences in membrane elasticities.** Plot on the average Cy3 lifetime changes for MM actuation from Supplementary Figure 26. Datapoints are from n = 10 GUVs per condition, across three independent experiments, and show the average and SD.

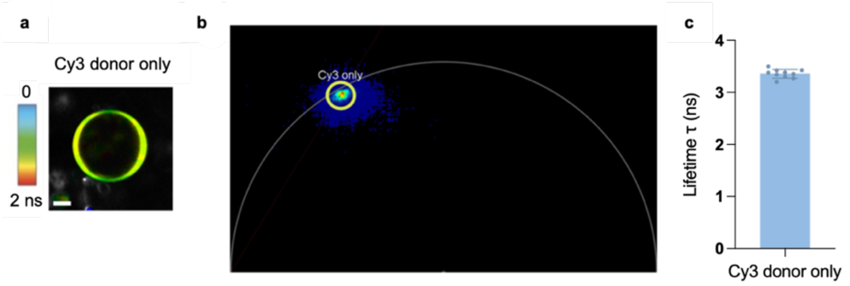

**Supplementary Figure 34. FLIM analysis Cy3-labeled MM bound to DOPC GUVs a)** Fast-FLIM image of a DOPC GUV incubated with Cy3-MM. The pixels are color-coded according to lifetime (scale 0-2 ns) and have a corresponding point in the phasor plot. Scale bar, 10 μm. **b)** Phasor plot of Cy3-MM, overlay of n = 10 fields of view. **c)** The average lifetime of Cy3-MM from phasor analysis is 3.4 ns ± 0.1, n= 10.

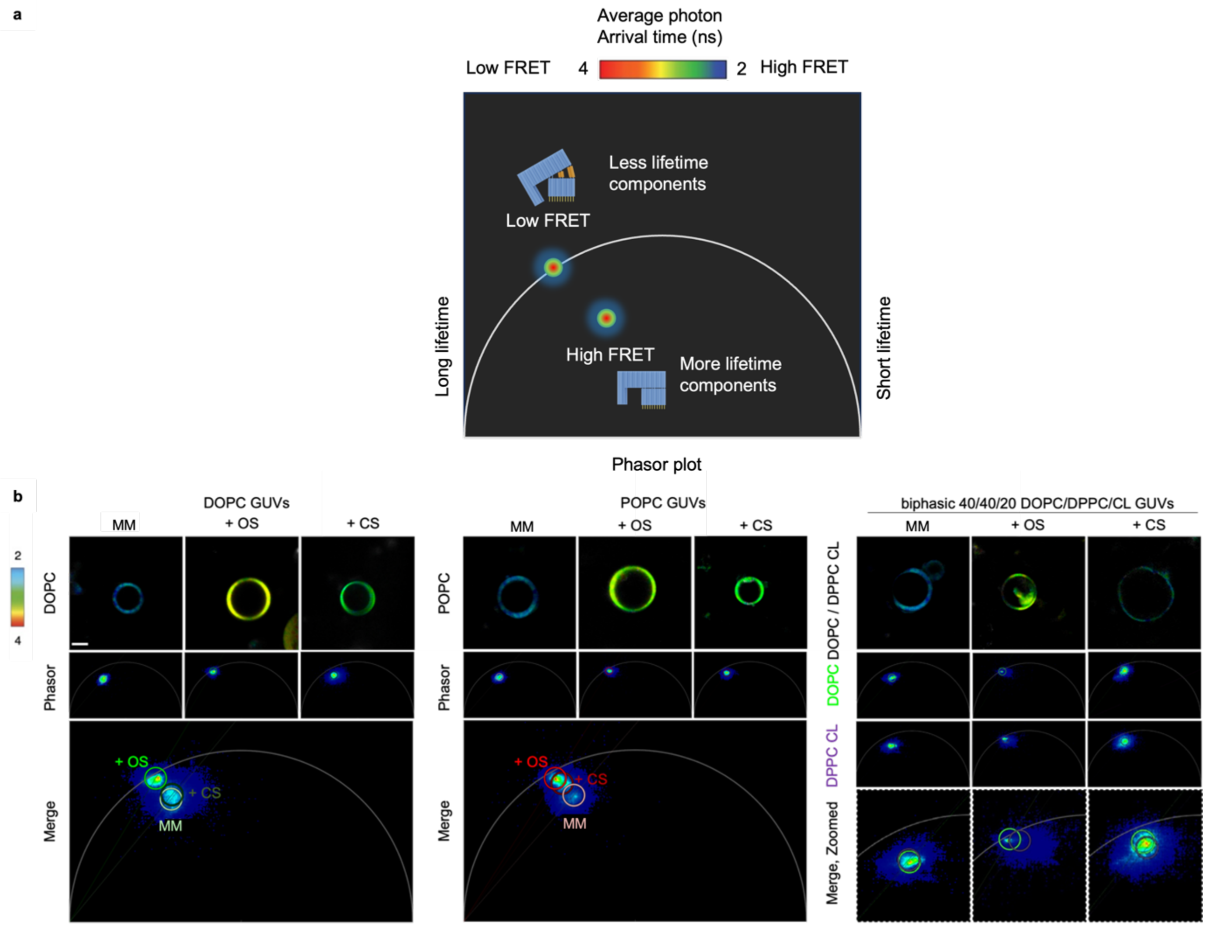

**Supplementary Figure 35. Phasor analysis of FLIM read-out of Cy3/Cy5-MM bound to vesicles. a)** Schematic diagrams of the phasor plot showing that the phasor dot position on the universal circle line indicates the fluorescence decay lifetime. Furthermore, the position of the phasor dot on the universal circle line indicates fewer lifetime components and no FRET, while a position towards the centre indicates more lifetime components and FRET. A component point in the phasor plot represents an image pixel in the FLIM image. **b)** Cy3 lifetime of MM in GUVs composed of DOPC, POPC and DOPC/DPPC CL. The data is shown for MM, the addition of opening strands (+OS), and closing strands (+CS). Panels show graphical 2D phasor FLIM distributions of n = 10 GUVs (overlaid). The bottom panels for DOPC and POPC show the merge phasor plots for MM, + OS, + CS. The bottom panels in biphasic illustrate phasor points of DOPC and DPPC CL phases merged.

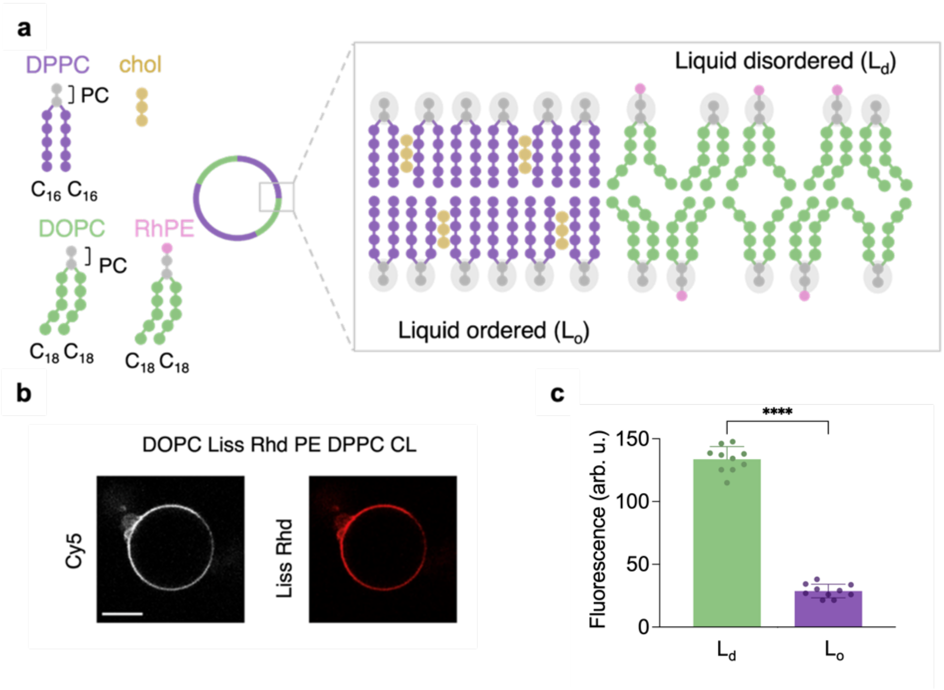

**Supplementary Figure 36. Microscopic analysis of MM preferentially binding to liquid disordered rich region. a)** Schematic for biphasic vesicles. DOPC and Liss Rhodamine PE form the liquid-disordered phase, DPPC and cholesterol form the liquid-ordered phase. **b)** Confocal microscopy images of Cy5-tagged MM bound to GUVs composed of DOPC/RhPE/DPPC/CL GUV (1% Rh), imaged under the Cy5 and Rhodamine channels, respectively. **c)** Bar chart shows mean membrane fluorescence intensity ± SD of DOPC and DPPC CL phases. N > 10 GUVs analysed across three independent experiments. Unpaired t-test, ****P <0.0001.

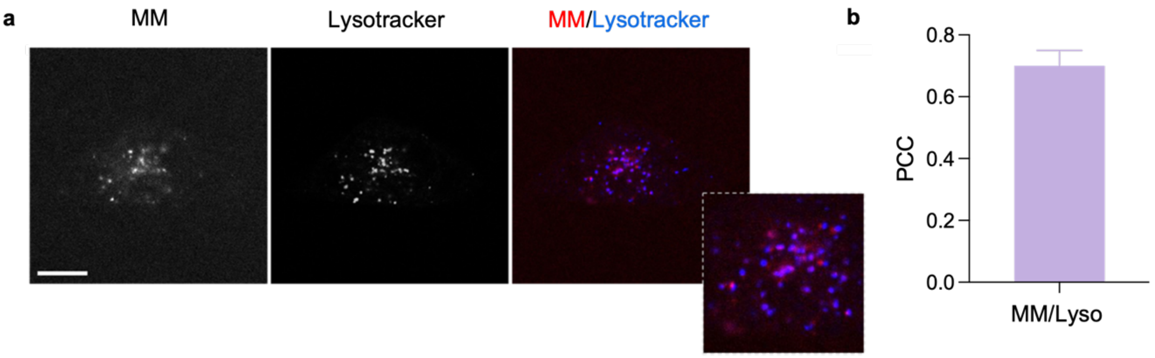

**Supplementary Figure 37. Rapid endocytosis of MM in live HeLa cells confirmed by co-staining with lysosomes and late endosomes. a)** Fluorescence confocal microscopy images of Cy3-tagged MM in live HeLa cells stained with green LysoTracker MM. For the MM/Lysotracker image, the MM is displayed in red, lysotracker is shown in blue. Purple indicates an overlay of MM and LysoTracker. Scale bar, 10 μm. **b)** Average Pearson correlation coefficient (PCC) ± SD of MM to Lysotracker.

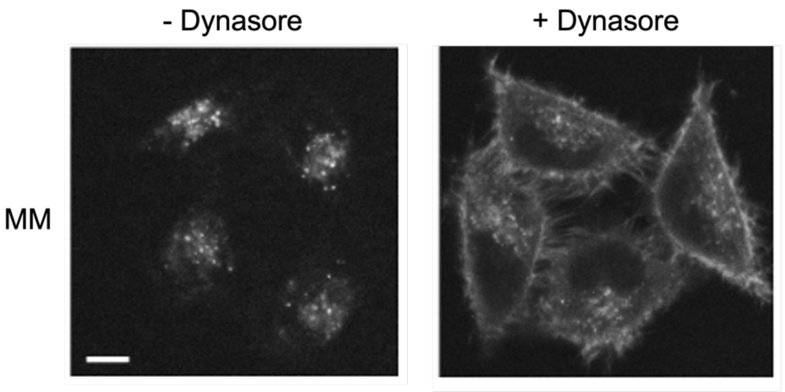

**Supplementary Figure 38. Dynamin inhibitor Dynasore increases the dwell time of MM at the plasma membrane.** Fluorescence confocal microscopy images of Cy3-tagged MM without the addition of Dynasore and after 30 min incubation with Dynasore (50 μM). Scale bar, 10 μm.

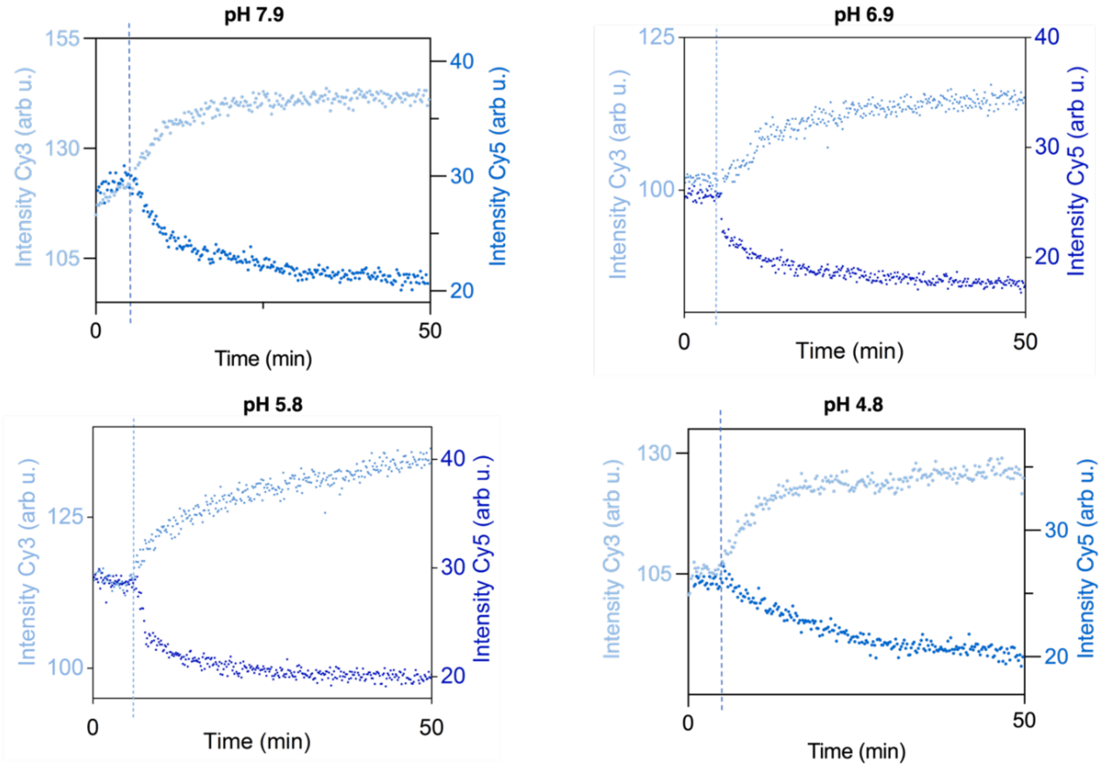

**Supplementary Figure 39. pH stability of FRET readout at a range of neutral to low pH.** Cy3 and Cy5 kinetic emission traces tracking MM opening at pH 7.9, 6.9, 5.8 and 4.8. Opening strands were added after 5 min (dashed line). pH was changed by supplementing the 16 mM $MgCl_2$ 1x TAE buffer with 1 mM HCl.

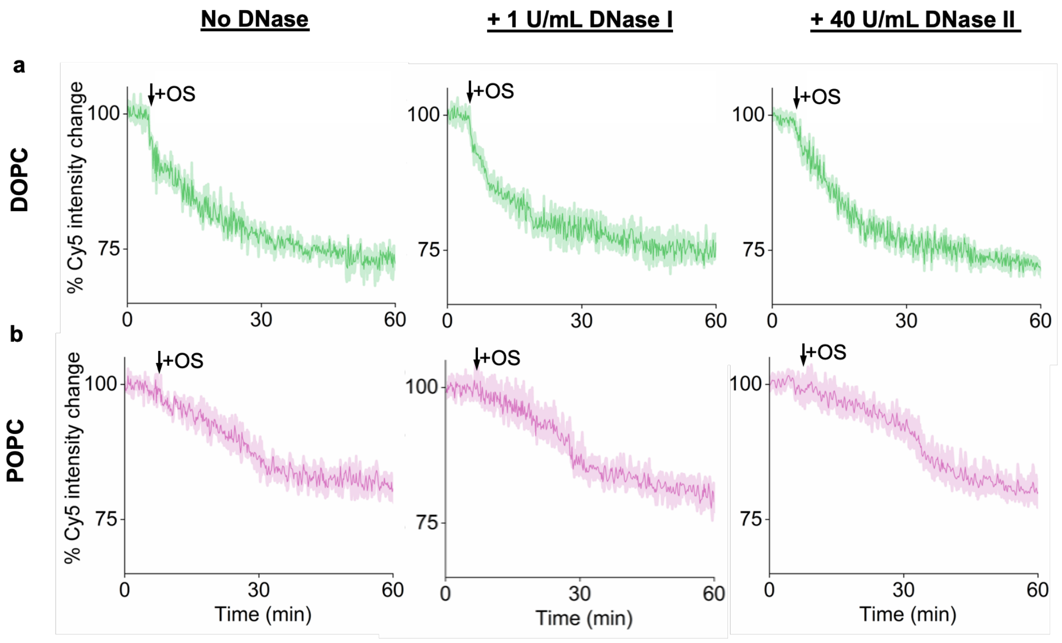

**Supplementary Figure 40. DNase treatment does not affect MM function**. To mimic an extracellular and intracellular nuclease challenge, MM was incubated with DNase I (1 U/mL) and DNase II (40 U/mL) at pH 6, respectively. After 15 min incubation, cholesterol strands were added and the mixtures were introduced to vesicles for binding of MM to membranes. The fluorescence Cy5 emission change was recorded shortly before and after OS addition (indicated by arrow). **a, b)** Cy5 emission traces for MM bound to DOPC (a) and POPC (b) in the absence of DNase (left) and in the presence of DNase I (middle) and DNase II (right). The data represent the means and standard deviation of four independent repeats.

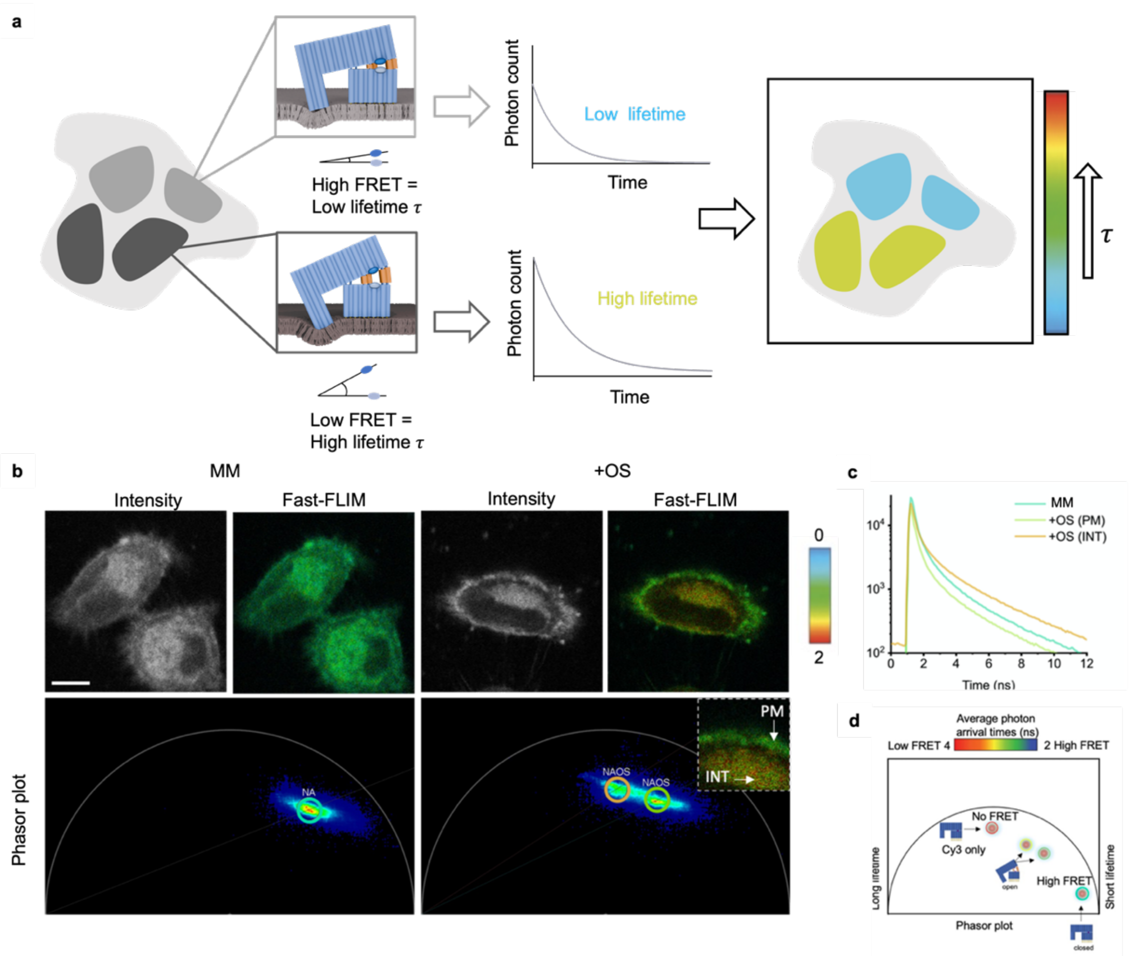

**Supplementary Figure 41. Fast-FLIM images, corresponding phasor plots, and lifetime decays of MM bound to live HeLa cells reveal distinct lifetimes from the plasma membrane (PM) and internal vesicles (INT). a)** Scheme for measuring membrane elasticities of different membrane regions via Fast-FLIM. **b)** Fast-FLIM images from main paper Figure 4 (scale 0-2) and corresponding phasor plots for MM + opening strands obtained from drawing ROIs on the PM and in INT of cells; overlay of n = 10 fields of view for each condition. Scale bar, 10 μm. **b)** Lifetime decay curves (average of n = 10 from three independent experiments) of closed MM and following the addition of OS in PM and INT regions. **c)** Schematic diagram of phasor plot demonstrating how the phasor point moves with FRET and the addition of OS.

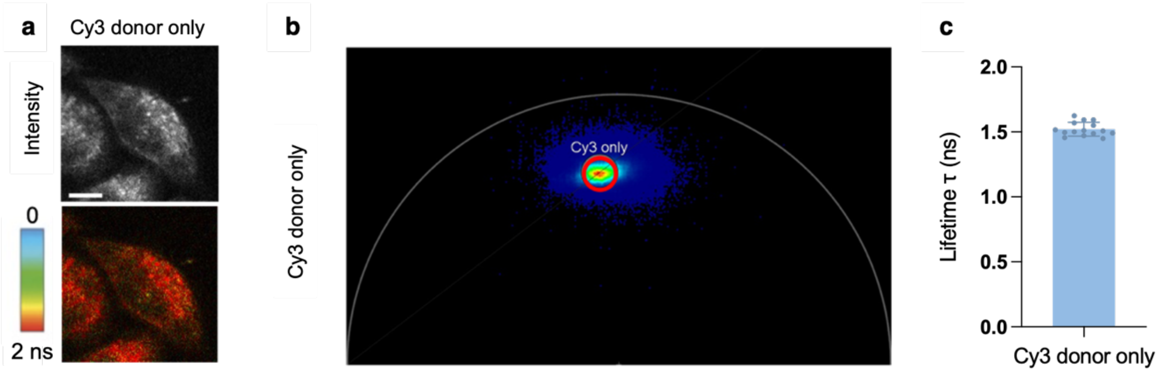

**Supplementary Figure 42. FLIM lifetime analysis of MM-Cy3 bound to cells. a)** Fast-FLIM image of live HeLa cells incubated with MM carrying only Cy3 but no Cy5. The pixels are coloured according to lifetime (scale 2-4 ns) and have a corresponding point in the phasor plot. Scale bar, 10 μm. **b)** Phasor plot of MM-Cy3, overlay of n = 15 fields of view. **c)** Lifetime of MM-Cy3 from phasor analysis.

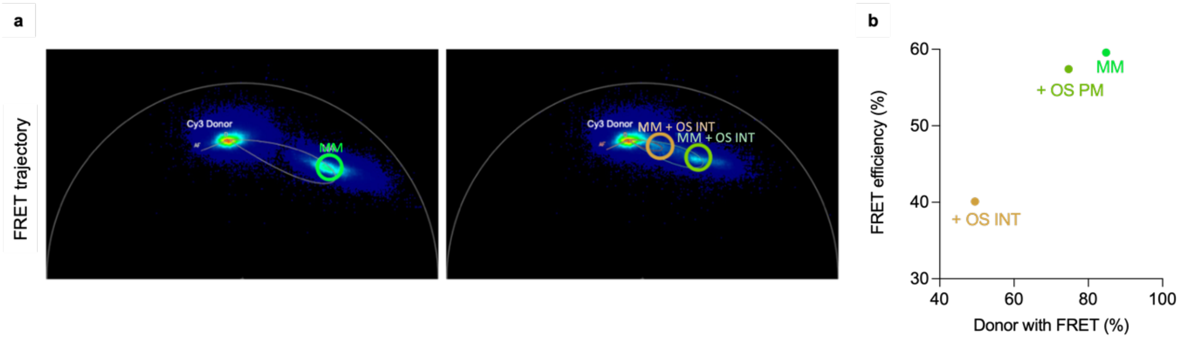

**Supplementary Figure 43. FRET trajectories reveal a higher FRET efficiency and number of FRET donors for MM bound at the plasma membrane (PM) than internal (INT) cell membranes.** FRET trajectories drawn from the MM-Cy3 data to the quenched and FRET-active MM data and the MM + opening strands data in the PM and INT. A circle is placed on each data to determine the number of donors in FRET and the FRET efficiency. Autofluorescence is considered. Scatter plot shows results from the FRET trajectory indicating the FRET efficiency and the number of donors with FRET (%). Data shows the overlay of n = 10 fields of view for each condition from three independent experiments.

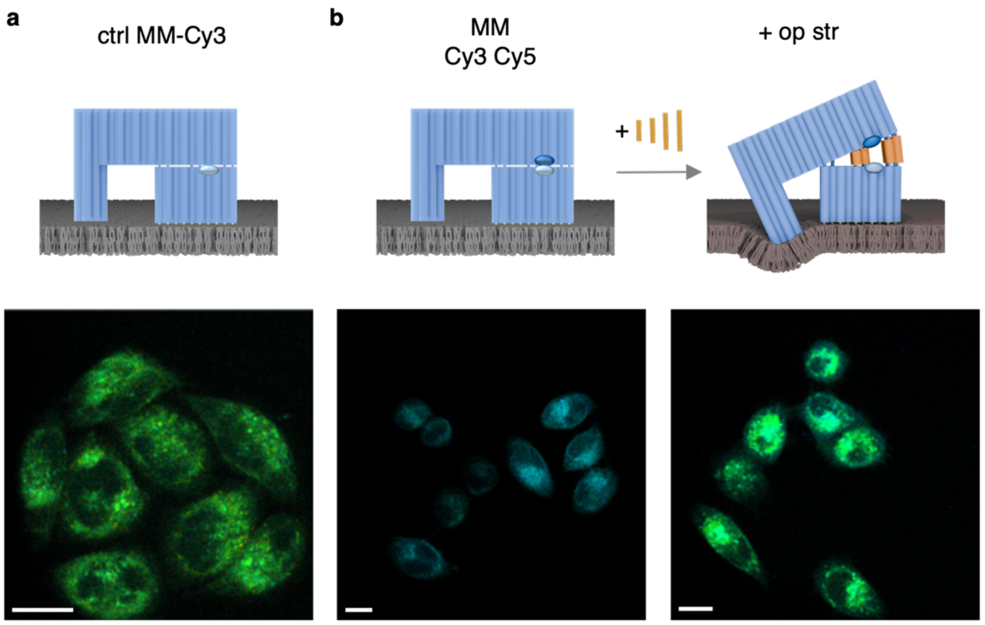

**Supplementary Figure 44. FLIM lifetime images of MM-bound to HeLa cells. a)** FLIM lifetime images of MM-Cy3 with Cy3 only as a control to determine the maximum unquenched lifetime. **b)** FLIM lifetime images of closed Cy3 Cy5 plus addition of opening strands (orange). Scalebar, 20μm.

### 2. Supplementary Tables

**Supplementary Table 1. Sequence of phiX174 scaffold strand**

| Sequence 5' → 3' |
|---|
| GAGTTTTATCGCTTCCATGACGCAGAAGTTAACACTTTCGGATATTTCTGATGAGTCGAAAAAT<br>TATCTTGATAAAGCAGGAATTACTACTGCTTGTTTACGAATTAAATCGAAGTGGACTGCTGGCG<br>GAAAATGAGAAAATTCGACCTATCCTTGCGCAGCTCGAGAAGCTCTTACTTTGCGACCTTTCG<br>CCATCAACTAACGATTCTGTCAAAAACTGACGCGTTGGATGAGGAGAAGTGGCTTAATATGCT<br>TGGCACGTTCGTCAAGGACTGGTTTAGATATGAGTCACATTTTGTTCATGGTAGAGATTCTCTT<br>GTTGACATTTTAAAAGAGCGTGGATTACTATCTGAGTCCGATGCTGTTCAACCACTAATAGGTA<br>AGAAATCATGAGTCAAGTTACTGAACAATCCGTACGTTTCCAGACCGCTTTGGCCTCTATTAAG<br>CTCATTCAGGCTTCTGCCGTTTTGGATTTAACCGAAGATGATTTCGATTTTCTGACGAGTAACA<br>AAGTTTGGATTGCTACTGACCGCTCTCGTGCTCGTCGCTGCGTTGAGGCTTGCGTTTATGGTA<br>CGCTGGACTTTGTAGGATACCCTCGCTTTCCTGCTCCTGTTGAGTTTATTGCTGCCGTCATTG<br>CTTATTATGTTCATCCCGTCAACATTCAAACGGCCTGTCTCATCATGGAAGGCGCTGAATTTAC<br>GGAAAACATTATTAATGGCGTCGAGCGTCCGGTTAAAGCCGCTGAATTGTTCGCGTTTACCTT<br>GCGTGTACGCGCAGGAAACACTGACGTTCTTACTGACGCAGAAGAAAACGTGCGTCAAAAATT<br>ACGTGCAGAAGGAGTGATGTAATGTCTAAAGGTAAAAAACGTTCTGGCGCTCGCCCTGGTCGT<br>CCGCAGCCGTTGCGAGGTACTAAAGGCAAGCGTAAAGGCGCTCGTCTTTGGTATGTAGGTGG<br>TCAACAATTTTAATTGCAGGGGCTTCGGCCCCTTACTTGAGGATAAATTATGTCTAATATTCAA<br>ACTGGCGCCGAGCGTATGCCGCATGACCTTTCCCATCTTGGCTTCCTTGCTGGTCAGATTGGT<br>CGTCTTATTACCATTTCAACTACTCCGGTTATCGCTGGCGACTCCTTCGAGATGGACGCCGTT<br>GGCGCTCTCCGTCTTTCTCCATTGCGTCGTGGCCTTGCTATTGACTCTACTGTAGACATTTTTA<br>CTTTTTATGTCCCTCATCGTCACGTTTATGGTGAACAGTGGATTAAGTTCATGAAGGATGGTGT<br>TAATGCCACTCCTCTCCCGACTGTTAACACTACTGGTTATATTGACCATGCCGCTTTTCTTGGC<br>ACGATTAACCCTGATACCAATAAAATCCCTAAGCATTTGTTTCAGGGTTATTTGAATATCTATAA<br>CAACTATTTTAAAGCGCCGTGGATGCCTGACCGTACCGAGGCTAACCCTAATGAGCTTAATCA<br>AGATGATGCTCGTTATGGTTTCCGTTGCTGCCATCTCAAAAACATTTGGACTGCTCCGCTTCCT<br>CCTGAGACTGAGCTTTCTCGCCAAATGACGACTTCTACCACATCTATTGACATTATGGGTCTGC<br>AAGCTGCTTATGCTAATTTGCATACTGACCAAGAACGTGATTACTTCATGCAGCGTTACCATGA<br>TGTTATTTCTTCATTTGGAGGTAAAACCTCTTATGACGCTGACAACCGTCCTTTACTTGTCATG<br>CGCTCTAATCTCTGGGCATCTGGCTATGATGTTGATGGAACTGACCAAACGTCGTTAGGCCAG<br>TTTTCTGGTCGTGTTCAACAGACCTATAAACATTCTGTGCCGCGTTTCTTTGTTCCTGAGCATG<br>GCACTATGTTTACTCTTGCGCTTGTTCGTTTTCCGCCTACTGCGACTAAAGAGATTCAGTACCT<br>TAACGCTAAAGGTGCTTTGACTTATACCGATATTGCTGGCGACCCTGTTTTGTATGGCAACTTG<br>CCGCCGCGTGAAATTTCTATGAAGGATGTTTTCCGTTCTGGTGATTCGTCTAAGAAGTTTAAGA<br>TTGCTGAGGGTCAGTGGTATCGTTATGCGCCTTCGTATGTTTCTCCTGCTTATCACCTTCTTGA<br>AGGCTTCCCATTCATTCAGGAACCGCCTTCTGGTGATTTGCAAGAACGCGTACTTATTCGCCA<br>CCATGATTATGACCAGTGTTTCCAGTCCGTTCAGTTGTTGCAGTGGAATAGTCAGGTTAAATTT<br>AATGTGACCGTTTATCGCAATCTGCCGACCACTCGCGATTCAATCATGACTTCGTGATAAAAGA<br>TTGAGTGTGAGGTTATAACGCCGAAGCGGTAAAAATTTTAATTTTTGCCGCTGAGGGGTTGAC<br>CAAGCGAAGCGCGGTAGGTTTTCTGCTTAGGAGTTTAATCATGTTTCAGACTTTTATTTCTCGC<br>CATAATTCAAACTTTTTTTCTGATAAGCTGGTTCTCACTTCTGTTACTCCAGCTTCTTCGGCACC<br>TGTTTTACAGACACCTAAAGCTACATCGTCAACGTTATATTTTGATAGTTTGACGGTTAATGCTG<br>GTAATGGTGGTTTTCTTCATTGCATTCAGATGGATACATCTGTCAACGCCGCTAATCAGGTTGT<br>TTCTGTTGGTGCTGATATTGCTTTTGATGCCGACCCTAAATTTTTTGCCTGTTTGGTTCGCTTTG<br>AGTCTTCTTCGGTTCCGACTACCCTCCCGACTGCCTATGATGTTTATCCTTTGGATGGTCGCC<br>ATGATGGTGGTTATTATACCGTCAAGGACTGTGTGACTATTGACGTCCTTCCCCGTACGCCGG<br>GCAATAATGTTTATGTTGGTTTCATGGTTTGGTCTAACTTTACCGCTACTAAATGCCGCGGATT<br>GGTTTCGCTGAATCAGGTTATTAAAGAGATTATTTGTCTCCAGCCACTTAAGTGAGGTGATTTA<br>TGTTTGGTGCTATTGCTGGCGGTATTGCTTCTGCTCTTGCTGGTGGCGCCATGTCTAAATTGTT<br>TGGAGGCGGTCAAAAAGCCGCCTCCGGTGGCATTCAAGGTGATGTGCTTGCTACCGATAACA<br>ATACTGTAGGCATGGGTGATGCTGGTATTAAATCTGCCATTCAAGGCTCTAATGTTCCTAACCC<br>TGATGAGGCCGTCCCTAGTTTTGTTTCTGGTGCTATGGCTAAAGCTGGTAAAGGACTTCTTGA<br>AGGTACGTTGCAGGCTGGCACTTCTGCCGTTTCTGATAAGTTGCTTGATTTGGTTGGACTTGG<br>TGGCAAGTCTGCCGCTGATAAAGGAAAGGATACTCGTGATTATCTTGCTGCTGCATTTCCTGA<br>GCTTAATGCTTGGGAGCGTGCTGGTGCTGATGCTTCCTCTGCTGGTATGGTTGACGCCGGAT<br>TTGAGAATCAAAAAGAGCTTACTAAAATGCAACTGGACAATCAGAAAGAGATTGCCGAGATGC |

AAAATGAGACTCAAAAAGAGATTGCTGGCATTCAGTCGGCGACTTCACGCCAGAATACGAAAG
ACCAGGTATATGCACAAAATGAGATGCTTGCTTATCAACAGAAGGAGTCTACTGCTCGCGTTG
CGTCTATTATGGAAAACACCAATCTTTCCAAGCAACAGCAGGTTTCCGAGATTATGCGCCAAAT
GCTTACTCAAGCTCAAACGGCTGGTCAGTATTTTACCAATGACCAAATCAAAGAAATGACTCGC
AAGGTTAGTGCTGAGGTTGACTTAGTTCATCAGCAAACGCAGAATCAGCGGTATGGCTCTTCT
CATATTGGCGCTACTGCAAAGGATATTTCTAATGTCGTCACTGATGCTGCTTCTGGTGTGGTTG
ATATTTTTCATGGTATTGATAAAGCTGTTGCCGATACTTGGAACAATTTCTGGAAAGACGGTAA
AGCTGATGGTATTGGCTCTAATTTGTCTAGGAAATAACCGTCAGGATTGACACCCTCCCAATTG
TATGTTTTCATGCCTCCAAATCTTGGAGGCTTTTTTATGGTTCGTTCTTATTACCCTTCTGAATG
TCACGCTGATTATTTTGACTTTGAGCGTATCGAGGCTCTTAAACCTGCTATTGAGGCTTGTGGC
ATTTCTACTCTTTCTCAATCCCCAATGCTTGGCTTCCATAAGCAGATGGATAACCGCATCAAGC
TCTTGGAAGAGATTCTGTCTTTTCGTATGCAGGGCGTTGAGTTCGATAATGGTGATATGTATGT
TGACGGCCATAAGGCTGCTTCTGACGTTCGTGATGAGTTTGTATCTGTTACTGAGAAGTTAAT
GGATGAATTGGCACAATGCTACAATGTGCTCCCCCAACTTGATATTAATAACACTATAGACCAC
CGCCCCGAAGGGGACGAAAAATGGTTTTTAGAGAACGAGAAGACGGTTACGCAGTTTTGCCG
CAAGCTGGCTGCTGAACGCCCTCTTAAGGATATTCGCGATGAGTATAATTACCCCAAAAAGAA
AGGTATTAAGGATGAGTGTTCAAGATTGCTGGAGGCCTCCACTATGAAATCGCGTAGAGGCTT
TGCTATTCAGCGTTTGATGAATGCAATGCGACAGGCTCATGCTGATGGTTGGTTTATCGTTTTT
GACACTCTCACGTTGGCTGACGACCGATTAGAGGCGTTTTATGATAATCCCAATGCTTTGCGT
GACTATTTTCGTGATATTGGTCGTATGGTTCTTGCTGCCGAGGGTCGCAAGGCTAATGATTCA
CACGCCGACTGCTATCAGTATTTTTGTGTGCCTGAGTATGGTACAGCTAATGGCCGTCTTCATT
TCCATGCGGTGCACTTTATGCGGACACTTCCTACAGGTAGCGTTGACCCTAATTTTGGTCGTC
GGGTACGCAATCGCCGCCAGTTAAATAGCTTGCAAAATACGTGGCCTTATGGTTACAGTATGC
CCATCGCAGTTCGCTACACGCAGGACGCTTTTTCACGTTCTGGTTGGTTGTGGCCTGTTGATG
CTAAAGGTGAGCCGCTTAAAGCTACCAGTTATATGGCTGTTGGTTTCTATGTGGCTAAATACGT
TAACAAAAAGTCAGATATGGACCTTGCTGCTAAAGGTCTAGGAGCTAAAGAATGGAACAACTC
ACTAAAAACCAAGCTGTCGCTACTTCCCAAGAAGCTGTTCAGAATCAGAATGAGCCGCAACTT
CGGGATGAAAATGCTCACAATGACAAATCTGTCCACGGAGTGCTTAATCCAACTTACCAAGCT
GGGTTACGACGCGACGCCGTTCAACCAGATATTGAAGCAGAACGCAAAAAGAGAGATGAGAT
TGAGGCTGGGAAAAGTTACTGTAGCCGACGTTTTGGCGGCGCAACCTGTGACGACAAATCTG
CTCAAATTTATGCGCGCTTCGATAAAAATGATTGGCGTATCCAACCTGCA

**Supplementary Table 2. Names and sequence of the DNA oligonucleotides used as staple strands to form MM.** Oligonucleotide main025 carries a 3' Cy3 modification and main078 carries an internal Cy5 modification.

| ID | Sequence 5' → 3' |
| --- | --- |
| main001 | CTGGCGTGAAGCAGAAGTATTTAGTAGCGGTAAGA |
| main002 | CCTTGAAGAGCTTTGCAGTCAACCATTATCAAATTG |
| main003 | AACCGAAGAAGATCAGCAAGCATTAACCGTCA |
| main004 | TACGTCCTGCGTGTAGCATCAACAGTTGCGG |
| main005 | AAGAGTCTCATTGCATATACCTGGTAGCAGTAATT |
| main006 | GGTCAAATCTGAAATTTGAATCTTATGGAACGCCCATGG |
| main007 | ACATATTCAAAATATAACGGTAATAAAGTTAGCTT |
| main008 | GCAAATCGACTGGAACTGTTCACCATAAGTAGGGT |
| main009 | GTTACTCAATCTTCTAAAAACCATTGGGC |
| main010 | ATTTGCGTCGCGTGCACCGCATGTTATGTAGG |
| main011 | GTTCTTTAATATCCCATACCGCTGATTGCCTGAATTTC |
| main012 | AGCAACAATTTCGCGAGTGATAAACGGTCACACTG |
| main013 | ACTAACGTACAGGAAAGCGAGGACCTACGGCAGCA |
| main014 | GGCAGTCGGGACTGATTAATGAAGAAAACCACTTT |
| main015 | CAGTAGTAATCAATATCCATAAAAATCGGTCACCA |
| main016 | ATAGCCAGATTTTTTTTCTGACTAGTTTGC |

| | |
|---|---|
| main017 | CTTCAATGTGGTAGAAGTAAAGCTCTAAGAACCGC |
| main018 | TATCCGAAAGTGTTAACGGCGTGGGCACACTCA |
| main019 | AGCTTCAGCGGCTTTAAACGTAATATGGAGA |
| main020 | CACTGGTCATAATCATGGTACAACACCATCCTTCACGG |
| main021 | ACCTATCAGGTATAACCTAACAGTTGAACTT |
| main022 | TAGTCATCATGACCAGCACCTTTCCTTTATCATGC |
| main023 | TTCAGCTTTACAATCCTAACATACTAGCAGGTAGAAAT |
| main024 | CGATGTAGCCATTACCCCAACAGAAACAACGGG |
| main025 | TAAAAAGCTGTAACCGACGAGCGCTGCGGGGCGAGCCACTCCT |
| main026 | TCATCCTTTGGGGTAATTACAGGCGCATAAA |
| main027 | ATACCAGCATCACCCATTATCGGTCGGCTTT |
| main028 | GTTGTGTTTTCTCTGTTGCATTTTGTTTGCATTCT |
| main029 | GAAACCAGAACAATCTTAACGATAGCAGGAGAAG |
| main030 | TTCTCATTCTTCGGTTCGTCAGTCAGTAGCAAT |
| main031 | GAATCTCCACCTTTCCAGCAATTGCCATCATA |
| main032 | TTTTCAGCAACGGAAATACAATTAAAATTGTTGCT |
| main033 | ACGACAGCGTGGCGAACCAAACAGGCAAAAAAAAAGTCAATGCCAGAT T |
| main034 | TAGCCTCCAGGCATCCACGGCACCCTGAGCAT |
| main035 | GGACGCTCGACGCCATTAATATTTAGACGCTAGTA |
| main036 | ATCAGAAACGGTCTGCAGCAGCAAGATAATCACAA |
| main037 | CAAAAAAGCCCCGCGGCGCCAGCCTGCAACGCTTT |
| main038 | AAGTGTCTCCATTCTTAGCCTAGAAA |
| main039 | CCAAATGCAGCAAGGTCCTTTGCGTTCTGCGGCTC |
| main040 | AGTCGATTAGAGCGCATGGTTGTCAAATAACACAT |
| main041 | TCAGCGGGCACCAACACCAGCGAATGGC |
| main042 | AACGCGGTTGGGAAATTTTCATTTGTCATTGG |
| main043 | GCATGGTCAAGTTAATCTCTGACCAAGCACTGGCA |
| main044 | AGCATGAGCTTCCAAAAAAACGCAAGCCTGCGAGTATC |
| main045 | TCCTAGACAATTCCAAGATACGCTCGCATTGGGGA |
| main046 | AGGTGTCACACCCAACATAAGTCCAACCAAATCGA |
| main047 | ATTTCCAGCAATCTTGTAACGA |
| main048 | GGTAGCACATAATTCATCCATTAATACC |
| main049 | ATTAGCAAGGGATAACCGAAGGAGGGAGAGCCGCGTTC |
| main050 | GAGACCATCTCGGAGTAGCGCAACGGCCTTTACGC |
| main051 | CCAAGTGCCAAGTCAAAAACGATAAGTCAGCCGGA |
| main052 | CTCGCGTCACTTAAGTGGCAAATTAGAGCCAAATA |
| main053 | GATTGTCCAGTTGCATTTTTTTTGAGCCTCGATTT |
| main054 | GCAAACCAAACCATGAAAAGTCCTTGACGGAGGTTGA |
| main055 | CACCAAACATTATTGCCCGAAGGACTCCAAAGATA |
| main056 | TGATTCAAAACTCCACCGCGCTTCGCTT |
| main057 | TCCTCATAGCACAAATAAAGCAATACAAAAATCTTC |
| main058 | TTGCTGCGATGCCAAGCGGTCTGGACATAAGACGA |
| main059 | ACGACAGAATCGATGCGGTTATCCAGAAAGAGTTT |
| main060 | GTACGATGAACCACTAACTAAAATAGAGCTTGGCT |

| | |
|---|---|
| main061 | AATGGTAATGATCGGACTCAGAGTGGAGTGGAGCG |
| main062 | TGCTACGCAAGGTAAACGTTCTTCTCGATAGCAAG |
| main063 | TTGGAGCAGCATCAGTGCACATAAATCACCGCAGT |
| main064 | AAGCCAATTTAGGGTCGCCATTATCGAACTCCCGT |
| main065 | AACGATTACATGCCAGAACGTTTTTAGTACCTTTGA |
| main066 | ACCATACTCATGAATCATTTAGCTACGAACACATC |
| main067 | ATATCTGGCTCGGCGCCAGTAGTAGATATAGC |
| main068 | TTGATCTGCTTTGCATACGAAAAGAACGTCAGGCA |
| main069 | ACGTTGAACGAACTGGCGGCGACAGCCCCTGATTT |
| main070 | ATCCACGCTACCATGAGGCGAATAAGTAGCC |
| main071 | GCAGCTAGAATGGGAAACATAGCGTCAGAGTCTCAGGA |
| main072 | AAAGCCTCTACTCCCAGCCGGCAAGTATCGGTATG |
| main073 | AACTACAGCATCAAAAGCAATACTCAAAACATTCAAAG |
| main074 | GGCGTTATAAATTGAATAGACATGCCGGAGGAGCA |
| main075 | CATCAGCGCGACCAGTCAATAGTCTGTAAAACAGGGAATGCAGCGGC GT |
| main076 | GTATCGCTCCCAAGCATTGATATAATAACCACCGG |
| main077 | AAATTGCATTCATCAAACTCGAAATGAAGACGCTGT |
| main078 | GTACCCGACGACCAATCAACGC |
| main079 | TTATACGAAAACAGCTTCCACAGAATGTTTCTTTG |
| main080 | GGAGAAGCAGCTATGGCGAGAACTTCTCAGTAATC |
| main081 | GACGCAACAATTAAATTAGTCATGCCTCACATTACCCC |
| main082 | AATCAAAAATACTTTGCGCCGCCAATAACTTTGCG |
| main083 | AAAGTATCAAGTTGGGGGGGTCTATAACAGCTCAC |
| main084 | CTCATAATAATAGAGGCCAGTAACGTGAAGCATTGGGA |
| main085 | AGATTTTGCCTAGGGACGGCCTCATCAGGAGCCTTAAG |
| main086 | ACGATAACCTGACCAGCTTGCGGCATCTTCTC |
| main087 | TCTGTTGAACACGATAGCTTTACCAACC |
| main088 | GAAACCTAGACCTTTAAATAGCAGTCTTCTGCGTC |
| main089 | CACTGATGAGGGATACTCATCGCGAGCGTTCA |
| main090 | TACCTATGTTAACGTATTTAGCCATCAAAGCGCAA |
| main091 | ACGTGTTAAAATGTGACTCATAGAATCTCTCTTTTTGATGA |
| main092 | CGTCAATCTCTAGTAAGCTCTTTTTGATTCT |
| main093 | TGAAGACGGTTGACTCCTCATAATAGACGCAAAAT |
| main094 | ATCATTTCACAATTAGCTAATCACGTTCTTGCCT |
| main095 | ATCGCGGAGCGCACGTTCGAACAAACATCACCTTA |
| main096 | TCTCGCTGAATTCAAATAGCTTTAATAAAAATACGT |
| main097 | CATATTAGAACAGCATTTCTTTAATAA |
| main098 | CAGATCATTGGCTTGCGAGTCATTTCTTCTGCGTT |
| main099 | ATCCAAGTATGAAACCTAGTAAGCATTTGGCGAT |
| main100 | GGAACTCAAGTTTTTGAATATTAGAACAGTAGA |
| main101 | TATCAGAAAAAAAGCATGATTGCGAAACCTGG |
| main102 | ATTAAGCTCATTCTCACCATAACGAGCATCATGCGAAC |
| main103 | TTACTATCTGATTCTGAATAGTCACTCGTAAACAAG |
| main104 | GTCAAATAAAAAGAATAGTTGTAACAC |

| | |
|---|---|
| main105 | GCCAGGCGGTTAATGCCAGCGACGACATTAGAATC |
| main106 | GAAGATAAGCAAATGTCAATAGAGGGGCCGAAGTATGCAGCTTCA |
| main107 | CAGATCCCGAAGGCCACAAAGCGGCTCACCTTAAATAGG |
| main108 | CTCAGGAAAGCCGACTGAAAATAATACCATAACAT |
| main109 | ATGGAAGCGATTGTCGTCACAGGGAGTCAAAGTTT |
| main110 | AGACCAGAAACAAAAATCGCGTACCTTCAAGAAG |
| main111 | AAGAGCGTCATAAGAGGTGCCCAGAGCAGTAGGCG |
| main112 | AACGACCAGAACGACGCATTTTGACAGTCCAAATG |

**Supplementary Table 3. Sequence of hinge sections with the phiX174 sequence**

| on 2D map | Sequence 5' → 3' |
|---|---|
| 8 | GGATT |
| 9 | TT |
| 10 | AA |
| 11 | GTCTT |
| 28 | GCAGA |
| 29 | GG |

**Supplementary Table 4. Sequences of actuator sections within phiX174 sequence and their position within the 2D duplex map in Supplementary Figure 3**. Actuator sequences form either a hairpin or a simple single-stranded loop, as indicated. The bold letters represent the bases which are hybridising to form the hairpin's stem.

| on 2D map | Sequence of hairpin structures 5' → 3' | Hairpin formation |
|---|---|---|
| 0 | ATGAGTCGAAAAATTAT**CTT**GATA**AAG**CAGG | Yes |
| 19 | CTTTGTTCCTGAG**CATG**GCAC**TATG**TTTACT | Yes |
| 20 | ACT**GAC**CAAAC**GTC**GTCGTTAGGC**CAG**TTTT**CTG** | Yes |
| 1 | TCGTATGGTTCTTGCTGCCGAGGGTCGC | No |
| 18 | CAACTCACTAAAAACCAAGCTGTCGCTA | No |
| 21 | **CAGT**TATATG**GCTG**TTGGTTTCTATGTG | Yes |
| 4 | TGC**GAC**AGGCTCATG**CTG**CTGA | Yes |
| 15 | TTA**CCA**AGC**TGG**GTTACGA | Yes |
| 24 | T**GC**AAAATACGTG**GC**CTTA | Yes |
| 5 | CAATAAAATCCCTAAG | No |
| 14 | CTTTC**CCA**TCT**TGG**CT | Yes |
| 25 | TTGGTATGTAGGTGGT | No |

**Supplementary Table 5. Names and sequences of the opening strand oligonucleotides.** Strands 01-06 are long OS and strands 07-12 are short opening strands.

| ID | Sequence 5' → 3' |
|---|---|
| opening strand 01 | CTACCATGGCCCTGCTTTATCAAGATAATTTTTCGACTCAT |
| opening strand 02 | CGTAAAGCTCAGTAAACATAGTGCCATGCTCAGGAACAAAG |
| opening strand 03 | CACGCGCATCCAGAAAACTGGCCTAACGACGTTTGGTCAGT |
| opening strand 04 | AGGACAGTATGCGACCCTCGGCAGCAAGAACCATACGA |
| opening strand 05 | TCAGAGCATA TAGCGACAGCTTGGTTTTTAGTGAGTTG |
| opening strand 06 | TAGTCGACGTCACATAGAAACCAACAGCCATATAACTG |
| opening strand 07 | GTAACACTTGTCAGCATGAGCCTGTCGCA |
| opening strand 08 | CTGTCTCATTTCGTAACCCAGCTTGGTAA |
| opening strand 09 | CAGTTGATTCTAAGGCCACGTATTTTGCA |
| opening strand 10 | TAGCGTTGGTCTTAGGGATTTTATTG |
| opening strand 11 | CAAGACTATCAGCCAAGATGGGAAAG |
| opening strand 12 | CTCTGACGGCACCACCTACATACCAA |

**Supplementary Table 6. Names and sequences of the closing strand oligonucleotides**

| ID | Sequence 5' → 3' |
|---|---|
| closing strand 01 | ATGAGTCGAAAAATTATCTTGATAAAGCAGGGCCATGGTAG |
| closing strand 02 | CTTTGTTCCTGAGCATGGCACTATGTTTACTGAGCTTTACG |
| closing strand 03 | ACTGACCAAACGTCGTTAGGCCAGTTTTCTGGATGCGCGTG |
| closing strand 04 | TCGTATGGTTCTTGCTGCCGAGGGTCGCATACTGTCCT |
| closing strand 05 | CAACTCACTAAAAACCAAGCTGTCGCTATATGCTCTGA |
| closing strand 06 | CAGTTATATGGCTGTTGGTTTCTATGTGACGTCGACTA |
| closing strand 07 | TGCGACAGGCTCATGCTGACAAGTGTTAC |
| closing strand 08 | TTACCAAGCTGGGTTACGAAATGAGACAG |
| closing strand 09 | TGCAAAATACGTGGCCTTAGAATCAACTG |
| closing strand 10 | CAATAAAATCCCTAAGACCAACGCTA |
| closing strand 11 | CTTTCCCATCTTGGCTGATAGTCTTG |
| closing strand 12 | TTGGTATGTAGGTGGTGCCGTCAGAG |

**Supplementary Table 7. Names and sequences of anchoring strand oligonucleotides which bind to cholesterol-TEG-modified DNA oligonucleotides.** The binding position to duplexes within the 2D map of Supplementary Fig. 1 is also shown. The lowercase letters in the sequence indicate the sections to which cholesterol-TEG-modified oligonucleotides bind.

| on 2D map | ID | Sequence 5' → 3' |
|---|---|---|
| 23 | cholesterol anchor01 | CTACAAAGTCCAGCtcatcttcacactact |
| 8 | cholesterol anchor02 | cctacgagtcagtcctCCAGTCCTTGACGA |
| 0 | cholesterol anchor03 | cctacgagtcagtcctAGCAGTCCACTTCGATTTAATGCAATT |
| 6 | cholesterol anchor04 | cctacgagtcagtcct TCTCCTCATCCAACGCGTCAACACCAAG |

| | | |
|---|---|---|
| 21 | cholesterol anchor05 | CCATACAACGCAGCGACGAGCACGtcatcttcacactact |
| 9, 10 | cholesterol anchor06 | cctacgagtcagtcctCAACAAGATCTAAAtcatcttcacactact |
| 1, 18 | cholesterol anchor07 | cctacgagtcagtcctAAATCATTTCCGCCtcatcttcacactact |
| 22, 17 | cholesterol anchor08 | cctacgagtcagtcctGTACCATCGGCAGAtcatcttcacactact |
| 28, 11 | cholesterol anchor09 | cctacgagtcagtcctCCTTCCAAAAATGTtcatcttcacactact |
| 7, 12 | cholesterol anchor10 | cctacgagtcagtcctAGTGGTTAGCCACTtcatcttcacactact |
| 4 | cholesterol anchor11 | cctacgagtcagtcct AGTTGATGGCGAAAGGTCGGTGAAGTTTTTGACAGCTTG |
| 25 | cholesterol anchor12 | ATAAACTCAAtcatcttcacactact |
| 24, 15 | cholesterol anchor13 | cctacgagtcagtcctCAGGAGCGGATTGTtcatcttcacactact |
| 26, 13 | cholesterol anchor14 | cctacgagtcagtcctATGAACAACCTATTtcatcttcacactact |
| 20, 19 | cholesterol anchor15 | cctacgagtcagtcctAGAGCGGAAAATCGtcatcttcacactact |
| 5, 14 | cholesterol anchor16 | cctacgagtcagtcctTCAGTAAAATCGTTtcatcttcacactact |
| 29 | cholesterol anchor17 | GACAGGCCATGTTTTCCGTAAATTCAGCGtcatcttcacactact |
| 2 | cholesterol anchor18 | cctacgagtcagtcctGCTGCGCAAGGATAGGTCGAAGACAAAGTAAG |
| 3, 16 | cholesterol anchor19 | cctacgagtcagtcctAGCCTGATTCTCGA tcatcttcacactact |
| 27 | cholesterol anchor20 | GCAATGTTTACCGTTTGAATGTTGACGGGtcatcttcacactact |

**Supplementary Table 8. Names and sequences of the cholesterol-TEG-modified DNA oligonucleotides which bind to the anchoring strands listed in Supplementary Table 7.** Oligonucleotide 3' chol was procured from IDT and 5' chol from Eurogentec.

| ID | Sequence 5' → 3' |
|---|---|
| 3' chol | GGA CTG ACT CGT AGG /3CholTEG/ |
| 5' chol | /5CholTEG/ AGT AGT GTG AAG ATG |

**Supplementary Table 9. Names, structure, and phase transition temperatures of lipids used in the study.**

| Lipid name | Chemical structure | $T_m$ (°C) |
|---|---|---|
| **DOPC**<br>18:1 (Δ9-Cis) PC<br>1,2-dioleoyl-sn-glycero-3-phosphocholine | 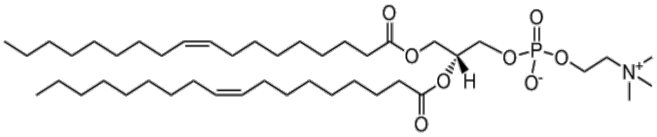 | -17 |
| **POPC**<br>16:0-18:1 PC<br>1-palmitoyl-2-oleoyl-glycero-3-phosphocholine | 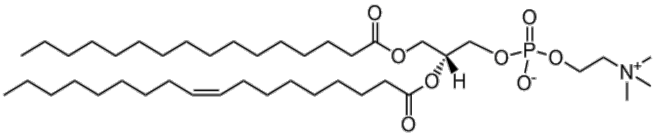 | -2 |
| **DPPC**<br>16:0 PC<br>1,2-dipalmitoyl-sn-glycero-3-phosphocholine | 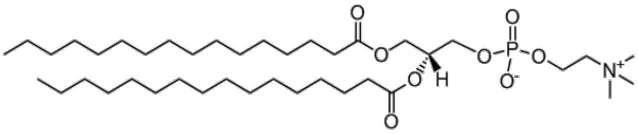 | 41 |
| **Liss Rhod PE 16:0**<br>1,2-dipalmitoyl-sn-glycero-3-phosphoethanolamine-N- (lissamine rhodamine B sulfonyl) (ammonium salt) | 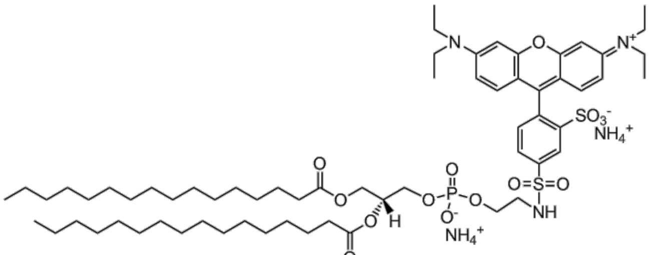 | N/A |
| **Cholesterol**<br>(plant derived) | 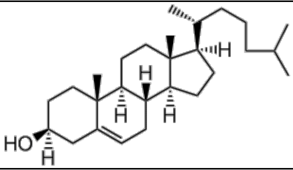 | N/A |

*Phase transition temperatures values obtained from Avanti polar lipids (https://avantilipids.com)

### 3. Supplementary Notes

#### Supplementary Note 1: Rational design of MM informed by free energy calculations

The design of the DNA-based molecular machine is defined by the relationship between the free energy gained by DNA hybridization $\Delta G_{opening}$, driving the indentation process, and the energy penalty of deforming a lipid membrane of given thickness $H$ and bending rigidity $\kappa_c$. We estimate the maximum deformation (indentation) that could be imposed on the membrane before it would break and form a toroidal pore around the DNA structure. Based on Hertz's theory describing the elastic indentation of a 2D half-space by an axisymmetric flat indenter, we assumed MM's tip to be cylindrical, and thus described the indentation force as in Acerbi et al.[2]:

$$F_I = 2RE^*\delta = 2R\left(\frac{1-\nu_{membrane}^2}{E_{membrane}} + \frac{1-\nu_{DNA}^2}{E_{DNA}}\right)^{-1} \delta \approx 2R\frac{E_{membrane}}{1-\nu_{membrane}^2}\delta$$

*Equation (1.1)*

in terms of Young's elastic modulus (*E*), Poisson ratio (*v*), indenter radius (*R*), indentation depth ($\delta$) and applied force (*F*).

Because the stiffness of a closely packed DNA nanostructure is much stiffer than lipid bilayers (i.e. $E_{DNA} \gg E_{membrane}$), the indentation force can be expressed as:

$$F_I \approx 2R\frac{E_{membrane}}{1-\nu_{membrane}^2}\delta$$

*Equation (1.2)*

The indentation force will be associated to a mechanical energy of the form:

$$U_I = \int_{\delta_0}^{\delta} F_I(l)\,dl = \int_{\delta_0}^{\delta} 2R\frac{E_{membrane}}{1-\nu_{membrane}^2}l\,dl = 2R\frac{E_{membrane}}{1-\nu_{membrane}^2}(\delta^2 - \delta_0^2)$$

*Equation (1.3)*

As we can consider the membrane to be incompressible, it follows that $\nu_{membrane}$ = 0.5. We also assume no indentation at time zero, so that $\delta_0$ = 0. Then, we end up with the expression:

$$U_I \approx \frac{8}{3}RE_{membrane}\delta^2$$

*Equation (1.4)*

According to thin-plate theory, the membrane's elastic modulus can be related to its bending rigidity ($\kappa_c$) by means of the following identity:[3]

$$\kappa_c = E_{membrane}H^3/12(1-\nu_{membrane}^2) \approx \frac{1}{9}E_{membrane}H^3$$

*Equation (1.5)*

Therefore, the indentation energy can be expressed as a function of the membrane's bending rigidity as:

$$U_I \approx \frac{8}{3}R\frac{9\kappa_c}{H^3}\delta^2 = \frac{24R\kappa_c}{H^3}\delta^2$$

*Equation (1.6)*

On the other hand, for a sufficient deformation the membrane would eventually be punctured, and a hydrophilic toroidal pore will be formed, in which lipid headgroups will rearrange to face the lumen. The energy cost for such a process is:[4]

$$U_P(R) = 2\pi\gamma_L R - \pi\gamma_S R^2$$

*Equation (1.7)*

Where *R* is the pore radius, $\gamma_L$ is the membrane's pore line tension and $\gamma_S$ is the membrane's surface tension. For lipid membranes where no external tension is applied, the squared term can be disregarded; so the energy cost for opening a pore grows linearly with *R*.[5]

According to *Wohlert et al.*[5] line tension can be compared to the bending rigidity $\kappa_c$ so that the energy for pore formation is simplified as:

$$U_P(R) \approx \frac{\kappa_c \pi^2 R}{H}$$

*Equation (1.8)*

Therefore, we can calculate the ratio $\varphi$ between the energy cost for both processes as:

$$\varphi = \frac{U_I}{U_P} = \frac{{24R\kappa_c\delta^2}/{H^3}}{{\kappa_c\pi^2 R}/{H}} = \frac{24}{(\pi H)^2}\delta^2$$

*Equation (1.9)*

From this relation, it then follows that indentation depth and membrane thickness would be the only parameters determining whether the bilayer will be punctured. Indeed, penetration would occur if $U_I > U_P$ (i.e. $\varphi > 1$). Hence, a critical indentation value $\delta_c$ can be calculated as:

$$\delta_c \coloneqq \delta \setminus \varphi = 1 \rightarrow \delta_c = \frac{\sqrt{6}\pi H}{12}$$

*Equation (1.10)*

Assuming a characteristic membrane thickness $H \sim$ 4 nm, the maximum allowed indentation is calculated to be $\delta_c \sim$ 2.6 nm. Considering a typical bending rigidity $\kappa_c \sim 10^{-19}$ J and $H \sim$ 4 nm, Eq. 1.10 can be used to relate the indentation energy with the tip radius, which for the above values is:

$$U_I(R) \sim 2.5 * 10^{-10} R$$

*Equation (1.11)*

On the other hand, the hybridization free energy for a DNA duplex of length 15-30 bp is about $10^{-19}$ J/duplex, hence the number of duplexes required to actuate the molecular machine and create an indentation $\delta_c$ on lipid membranes can be expressed as:

$$N_{OPduplex} \sim 2.5\ nm^{-1}\ R$$

*Equation (1.12)*

To lower the device's complexity, $N_{OPduplex}$ should be kept to a minimum. The lower bound for *R* is limited by the honeycomb lattice used in the DNA origami design, i.e. a single hexagonal arrangement of the DNA duplexes with an equivalent radius of *R* ~ 3 nm. Based on design considerations, however, we decided to extend the tip's cross-section (Supplementary Fig. 2c) to increase the rigidity of the DNA nanostructure, to an equivalent radius $R_{eq}$.

$$R_{eq} = \sqrt{ab/\pi}$$

*Equation (1.13)*

where $a$, $b$ correspond to the 2 dimensions (i.e. “width” and “length”, 7.2 nm and 9 nm, respectively) of the MM’s tip; resulting in $R_{eq}$ ~ 4.54 nm, thus requiring ~ 12 opening strands.

**Supplementary Note 2: Converting FLIM-FRET lifetimes into membrane indentation depths**

Fluorescence lifetime imaging microscopy (FLIM) does not depend on the local fluorophore concentration, and is robust against intensity-based FRET readouts, such as channel bleed through; hence FLIM-FRET is the preferred technique to quantify the degree of indentation by MM.

Lifetime-based FRET efficiency can be calculated as:

$$E_{FRET} = 1 - \frac{\tau_{DA}}{\tau_D}$$

*Equation (2.1)*

where $\tau_{DA}$ represents the donor's (Cy3) lifetime in the presence of the acceptor (Cy5), and $\tau_D$ represents the fluorescence lifetime of the donor on its own. Importantly, we note that Cy3 is a well-known environmentally sensitive probe[67]; its photophysical behaviour will depend on the local DNA geometry (Supplementary Fig. 5). Therefore, specific Cy3 lifetime calibration should be performed when attached to different nucleic acid nanostructures.

On the other hand, the FRET efficiency between a dye with Förster radius $D_0$ can be related to the donor-acceptor separation $d$ as:

$$E_{FRET} = \frac{1}{1 + \left(\frac{d}{D_0}\right)^6}$$

*Equation (2.2)*

Therefore:

$$d = D_0 \sqrt[6]{\frac{1}{1 - \frac{\tau_{DA}}{\tau_D}} - 1}$$

*Equation (2.3)*

Based on MM's design, the distance between the FRET pair can be used to calculate the opening angle $\theta$ as:

$$\theta = \cos^{-1}\left(\frac{1}{2L_D L_A}(L_D^2 + L_A^2 - d^2)\right)$$

*Equation (2.4)*

where $L_D$ and $L_A$ represent the distance from the dyes to the pivot's point (Supplementary Fig. 13). Once the opening angle is known, it is possible to calculate the indentation depth $\delta$ at the point $P_{tip}$ as:

$$\delta = P_y \sin(\theta) + P_x \cos(\theta) - P_y - \rho$$

*Equation (2.5)*

where $P_x = -20$ nm and $P_y = -15$ nm represents the tip position relative to the pivot point (Supplementary Fig. 13) and $\rho = -5$ nm represents any offset (e.g. due to the cholesterol-TEG linker).

Taking the total indentation as the difference between the estimated indentation depths before and after the addition of the OS:

$$\Delta\delta = \delta_{OS} - \delta_0$$

*Equation (2.6)*

Then Eq. 1.2, Eq. 1.5 and Eq. 2.6 can be combined to estimate the bending rigidity, where $F_{OS}$ is derived from the Worm-Like Chain model (Eq. 3.1):

$$\kappa_{c,estimate} = \frac{F_{OS} H^3}{24 R \Delta\delta}$$

*Equation (2.7)*

Combining the estimated force and the inferred $\Delta\delta$ from FRET measurements, recalling the tip radius $R_{eq} \sim 4.54$ nm and setting the membrane thickness $H = 4.52$ nm we estimated the bending rigidity $\kappa_{c,estimate}$ for the different lipid membranes (Supplementary Table 10). The bending rigidities are slightly lower compared to values obtained with other techniques (Supplementary Table 11), possibly due to some flexibility in the anchoring of MM to membranes.

**Supplementary Table 10. Parameters used to calculate bending rigidity *k*c.** The indentation depths were derived from FLIM-fitting analysis. The same membrane thickness was used throughout the calculations for $k_c$ for reasons of consistency. A total force of 32 pN was assumed, based on Eq. 3.1. Error represents ±SD except for $\kappa_c$, representing SEM. Note the fitted lifetimes can differ slighty from the 'Fast-FLIM' values (representing the average photon arrival times) provided in Fig. 4 and Supplementary Figs. 32-34, 41-42.

| Parameter | DOPC | POPC | $L_d$ | $L_o$ | Plasma membrane | Internal organelles |
|---|---|---|---|---|---|---|
| $\tau_{Cy3only}$ (ns) | 3.4 ± 0.1 | 3.5 ± 0.1 | 3.3 ± 0.1 | | | |
| $\tau_{closed}$ (ns) | 2.6 ± 0.2 | 2.7 ± 0.2 | 2.6 ± 0.2 | 2.6 ± 0.2 | 2.9 ± 0.2 | 2.9 ± 0.2 |
| $\tau_{open}$ (ns) | 3.2 ± 0.2 | 3.1 ± 0.1 | 3.1 ± 0.2 | 2.8 ± 0.3 | 3.1 ± 0.1 | 3.2 ± 0.1 |
| $d_{FRET,closed}$ (nm) | 6.5 ± 0.3 | 6.5 ± 0.3 | 6.4 ± 0.4 | 6.3 ± 0.3 | 6.9 ± 0.5 | 7.4 ± 1.0 |
| $d_{FRET,open}$ (nm) | 9.1 ± 1.6 | 8.3 ± 1.1 | 8.0 ± 0.7 | 7.1 ± 0.8 | 7.6 ± 0.4 | 8.5 ± 0.8 |
| $\theta_{closed}$ (deg) | 36 ± 2 | 36 ± 2 | 35 ± 2 | 35 ± 2 | 38 ± 3 | 41 ± 6 |
| $\theta_{open}$ (deg) | 47 ± 3 | 45 ± 3 | 45 ± 4 | 40 ± 4 | 43 ± 2 | 49 ± 5 |
| $\delta_{closed}$ (nm) | -3.9 ± 0.3 | -3.9 ± 0.2 | -3.8 ± 0.3 | -3.8 ± 0.2 | -4.2 ± 0.3 | -4.4 ± 0.3 |
| $\delta_{open}$ (nm) | -4.9 ± 0.2 | -4.7 ± 0.2 | -4.7 ± 0.3 | -4.3 ± 0.5 | -4.6 ± 0.2 | -4.9 ± 0.1 |
| $\Delta\delta$ (nm) | -0.96 ± 0.22 | -0.77 ± 0.24 | -0.86 ± 0.30 | -0.57 ± 0.42 | -0.41 ± 0.20 | -0.46 ± 0.14 |
| **$\kappa_c$ (x$10^{-20}$ J)** | **3.4 ± 0.5** | **4.3 ± 0.7** | **4.0 ± 0.7** | **6.7±2.0** | **8.9 ± 2.0** | **7.0 ± 0.7** |

**Supplementary Table 11**. **Bending rigidities for DOPC and POPC** from the literature obtained via various techniques.

| **Analysis techn.** | **DOPC** | **POPC** | **Source** |
|---|---|---|---|
| Micropippete | $8.5x10^{-20}$ J | $9x10^{-20}$ J (C18:0/1, comparable to POPC | Rawicz et al. Biophys Journal, 2000 (Ref [8]) |
| Fluctuation analysis | $11x10^{-20}$ J | $14x10^{-20}$ J | Elani et al. Chem Comm 2015 (Ref.[9]) |
| Simulation | $7.4x10^{-20}$ J | $9.9x10^{-20}$ J | Doktorova et al. PCCP 2017 (Ref.[10]) |
| Fluctuation analysis | $2.4x10^{-20}$ J | $3.9x10^{-20}$ J | Niggemann et al, Journal de Physique II, 1995 (Ref.[11]) |
| Electrodeformation | $6.1x10^{-20}$ J | $5.8x10^{-20}$ J | Niggemann et al, Journal de Physique II, 1995 (Ref.[11]) |
| Fluctuation analysis (in water) | $17.8x10^{-20}$ J | $21.2x10^{-20}$ J | Drabik et al. BBA-Biomembranes 2016 (Ref.[12]) |
| Fluctuation analysis (in sucrose) | $13.3x10^{-20}$ J | $15.1x10^{-20}$ J | Drabik et al. BBA-Biomembranes 2016 (Ref.[12]) |
| XRD | $6.7x10^{-20}$ J | $7.9x10^{-20}$ J | John F. Nagle, Chemistry and Physics of Lipids 2017 (Ref.[13]) |
| Simulation | $11.8x10^{-20}$ J | $13.2x10^{-20}$ J | Venable et al. Chemistry and Physics of Lipids 2015 (Ref.[14]) |

**Supplementary Note 3: Mechanical characterization of the molecular machine**

To assess the capability of MM to indent lipid membranes, we quantified the force exerted by the addition of opening strands.

We modelled the force caused by the opening of MM as that of an entropic spring. According to the Worm-Like Chain model, the addition of each opening strand will contribute:

$$F_{OS} \approx \frac{k_B T}{L_p}\left[\frac{1}{4\left(1-\frac{R}{L_c}\right)^2}-\frac{1}{4}+\frac{R}{L_c}\right]$$

*(Equation 3.1)*

where $L_p$ is the persistence length for ssDNA (1.5 nm), $R = 0.34N - 2$ nm/bp is the difference in the end-to-end extension upon hybridisation of the opening strands, and $L_c = 0.65\ N\ nm/bp$ is the contour length for the strands connecting the tip and base of the MM where *N* is the number of base pairs. Using these values, we estimated that the total force to be 31.97 pN upon full actuation of the MM. Similar forces have been reported by other DNA origami-based nanoactuators.

Using this insight, we explored whether the readout from our MM followed Hertz's indentation model by recording the fluorescence readout from MM bound to DOPC and POPC membranes upon the addition of six short, six long, or all opening strands. Using Eq. 3.1, we estimated the force for a given set of opening strands and compared that to the change in Cy5 signal (Fig. 3d, Supplementary Table 12).

**Supplementary Table 12. Estimated indentation force and change in Cy5 intensity** upon OS addition for different OS sets. Contrast refers to Cy5_POPC-Cy5_DOPC and is related to the sensitivity of our MM to the membrane's elasticity.

| Set | Force (pN) | Cy5_DOPC | Cy5_POPC | *Contrast* |
|---|---|---|---|---|
| Short | 13.70 | -17.310 | -16.929 | -0.381 |
| Long | 18.26 | -22.386 | -20.660 | -1.726 |
| Both | 31.97 | -28.198 | -21.777 | -6.421 |

If comparing the change in relative indentation depth $\Delta\Delta\delta = \Delta\delta_{POPC} - \Delta\delta_{POPC}$ then, for a given force:

$$\Delta\Delta\delta = \frac{F_{OS}H^3}{24R\kappa_{c,POPC}} - \frac{F_{OS}H^3}{24R\kappa_{c,POPC}} = \frac{F_{OS}H^3}{24R}\left[\frac{\kappa_{POPC}-\kappa_{DOPC}}{\kappa_{POPC}\kappa_{DOPC}}\right]$$

*(Equation 3.2)*

On the other hand, the FRET efficiency from intensity-based measurements may be approximated as:

$$E_{FRET} = \frac{I_A}{I_A + I_D} = \frac{1}{1+\left(\frac{d}{R_0}\right)^6}$$

*(Equation 3.3)*

Therefore:

$$I_A = I_D\left(\frac{d}{R_0}\right)^6$$

*(Equation 3.4)*

Now, considering the relative change in Cy5 acceptor intensity, then:

$$\Delta Cy5 = 1 - \frac{I_{A,OS}}{I_{A,0}} = 1 - \left(\frac{d_{OS}}{d_0}\right)^6$$

*(Equation 3.5)*

Hence:

$$d_{OS} = d_0 \sqrt[6]{1 - \Delta Cy5}$$

*(Equation 3.6)*

where $d_0$ is the FRET distance in the absence of opening strands, which we take as 6.5 nm from our FRET-FLIM measurements.

If we assume that the indentation depth and FRET distance are proportional, e.g. $d_{OS} \sim \delta_{OS}$ and $d_0 \sim \delta_0$, then we can express:

$$\Delta\delta = \delta_{OS} - \delta_0 = d_0 \left(\sqrt[6]{1 - \Delta Cy5} - 1\right)$$

*(Equation 3.7)*

Finally, combining Eq. 3.7 and Eq. 3.2 yields Eq. 3.8, which is plotted in Supplementary Fig. S45a:

$$\Delta Cy5 = 1 - \left[1 + \left(\frac{FH^3}{24R\kappa_c d_0}\right)\right]^6$$

*(Equation 3.8)*

Comparison of Supplementary Fig. S45a to the experimental Cy5 intensity changes in Fig. 3d, Supplementary Fig. S45b yields a good match in terms of two observables. Increasing the number of OS increased the change of Cy5 intensity, and the change is less prominent for stiffer vesicles (POPC). Altogether, this suggests that MM has the characteristics required to report on the bending rigidity of lipid membranes.

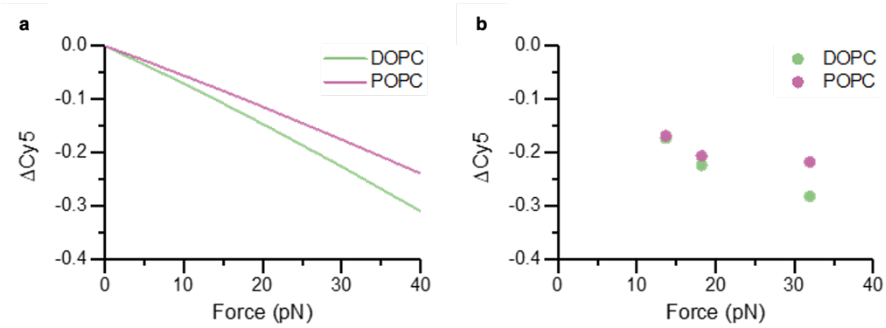

**Supplementary Figure 45. a)** Plot of relative change in Cy5 against force given Eq. 3.8, compared to **b)** the experimental data obtained using different sets of opening strands.

The maximum of 32 pN generated by MM opening is too small to pull out MM's membrane anchors as this would require a force between 600 and 1650 pN. The lower boundary of the force range of pulling out the membrane anchors is derived from the extraction of cholesterol anchors from the membrane (20 pN/cholesterol[15] )multiplied by the 30 membrane anchors. The upper boundary is obtained when considering that the cholesterol-modified DNA strands are bound via a duplex of 20 bp length to the MM whereby each duplex shearing requires >55 pN force[16] which multiplies to more than 1650 pN for 30 duplexes.

**Supplementary Note 4: Error propagation, and the method's analytical resolution and dynamic range**

A bootstrapping approach was used to establish error propagation and the MM's measurement resolution and analytical range.

4.1. Error propagation

A set of three lifetimes was randomly sampled from each lifetime distribution ($\tau_D, \tau_0, \tau_{OS}$) and then used to calculate $\kappa_c$ according to the formulas Eq. 2.1 – 2.7 (Supplementary Note 2). To account for structural flexibility, the dimensions of the DNA origami (Supplementary Fig. 2) were allowed to fluctuate within ±1 nm. Unrealistic values (e.g. where $E_{FRET} < 0 \text{ or } E_{FRET} > 1, \text{ or } \Delta\delta > 0$) were removed from the analysis.

Supplementary Figs. 46 and 47 show the distributions for each parameter used for deriving $\kappa_c$. When fluctuations were considered (Supplementary Figs. 46a and 47a), broader distributions and a systematically lower $\kappa_c$ estimate was observed, owing to the wider distribution of indentation depths.

**a**

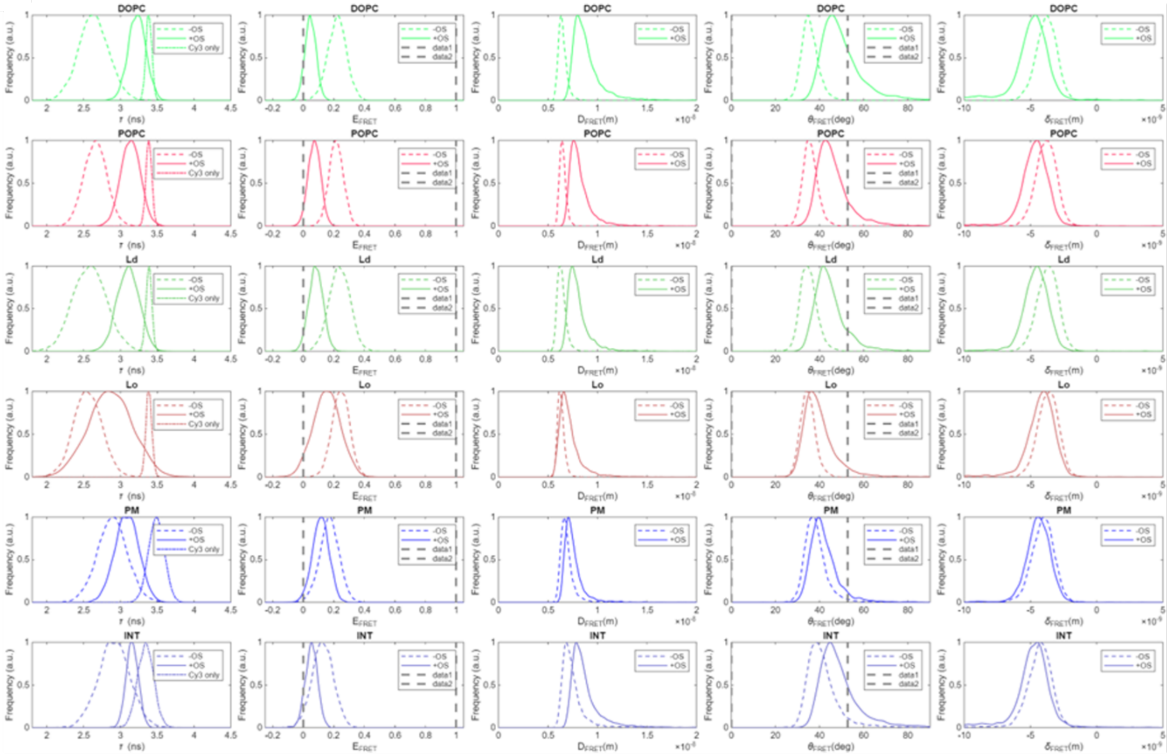

**b**

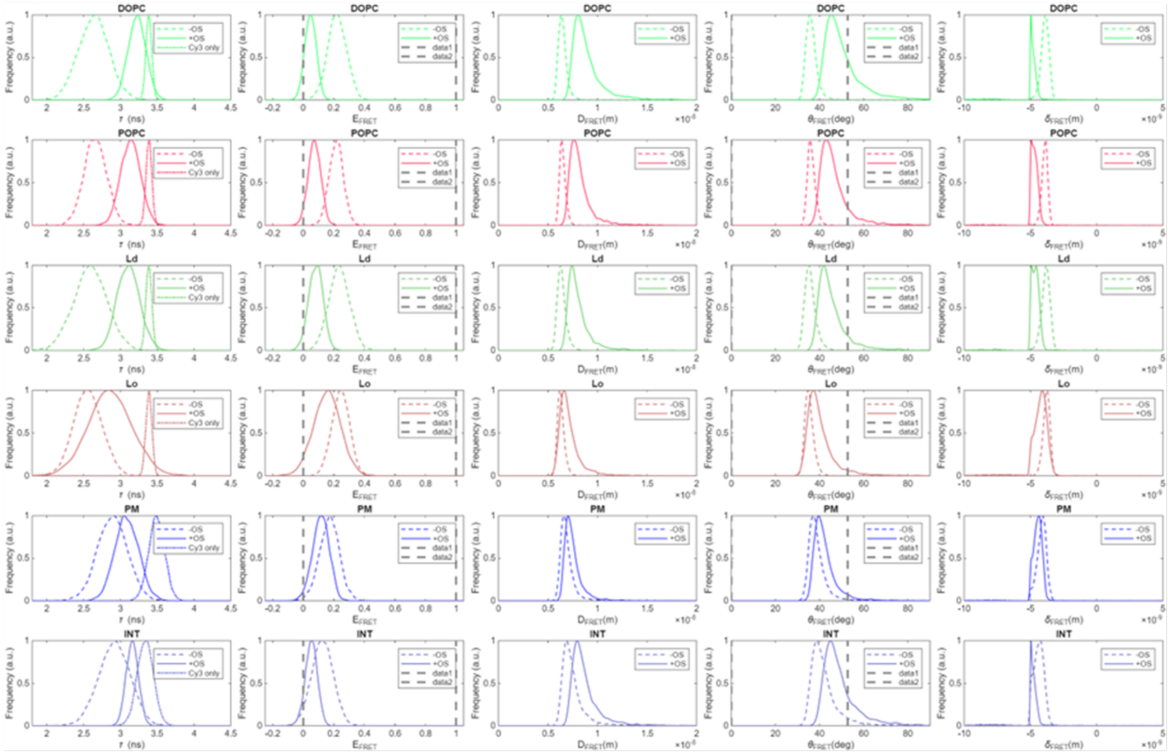

**Supplementary Figure 46. Effect of error propagation in the calculation of MM geometry.** From left to right, lifetime distributions, FRET efficiency, FRET distance, opening angle, and estimated indentation for the MM before (-OS) and after (+OS) actuation. Calculations were performed for measurements in GUV and cells, **a)** considering fluctuations of the DNA structure or **b)** assuming the DNA origami to be a rigid body. Vertical dashed lines represent the threshold for unrealistic values.

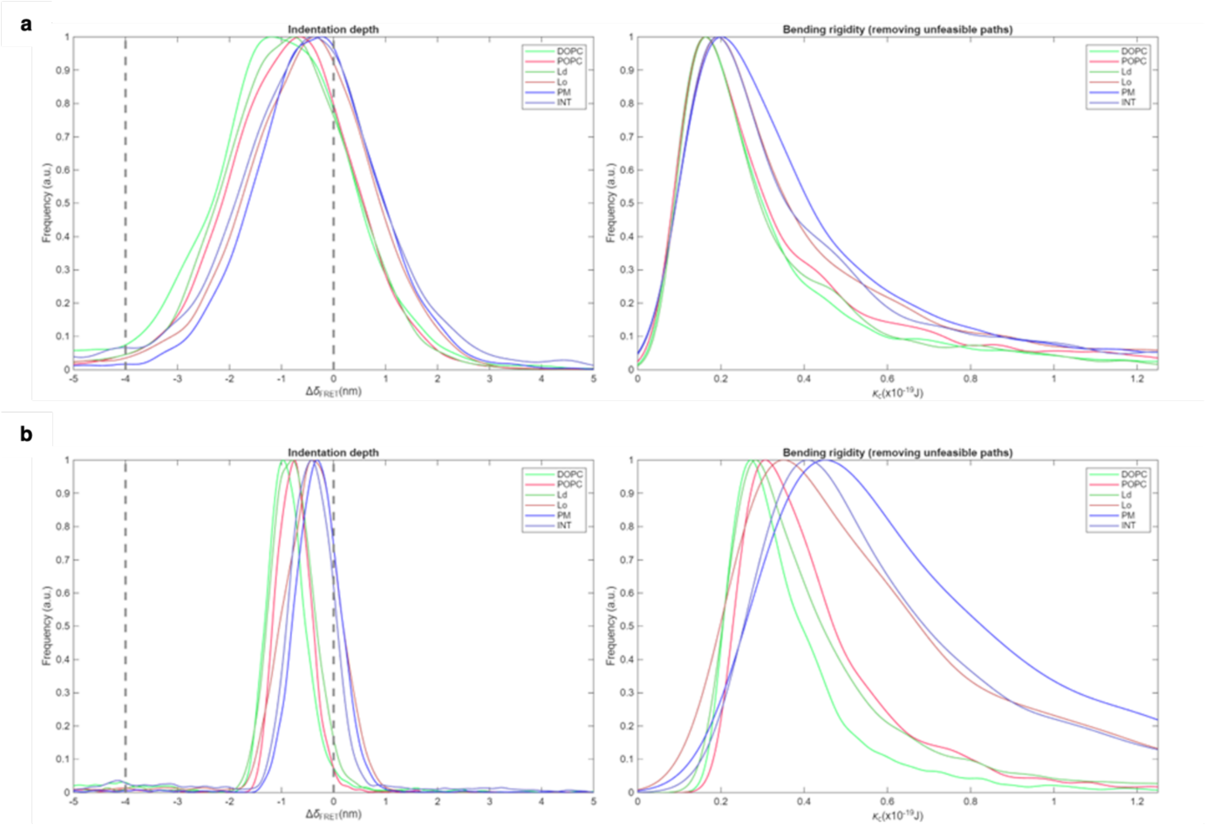

**Supplementary Figure 47. Effect of error propagation in the calculation of membrane elasticity.** Change in indentation depth and estimated bending rigidity when **a)** fluctuations of the DNA structure are considered or **b)** the DNA origami is assumed to be a rigid body. Vertical dashed lines represent the threshold for unrealistic values.

Non-parametric Kolmogorov-Smirnov (K-S) test was used to assess the difference between the curves. To estimate how many points should be drawn from the distributions, the KS score was calculated by drawing an increasing number the generated points from the same distribution. As the parent distribution is the same, the KS score vs $N_{points}$ curve should flatten when enough points are drawn to faithfully represent the distribution. Based on the data from Supplementary Fig. 48, a value of N = 250 when performing the KS test to assess differences between bending rigidities.

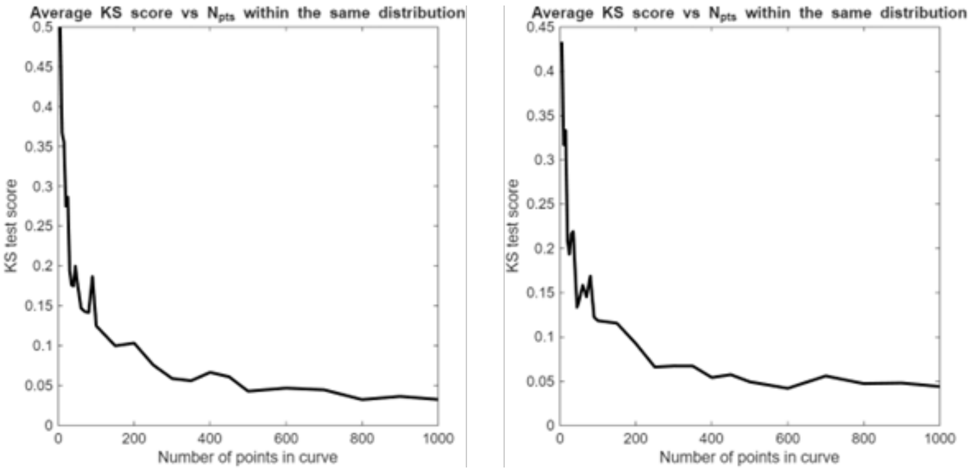

**Supplementary Figure 48. Effect number of points drawn from an empirical distribution on the KS score. a)** When considering fluctuations of the DNA structure, and **b)** assuming the DNA origami to be a rigid body.

Supplementary Fig. 49 shows the pair-wise asymptotic p-values obtained by performing the KS test between a given pair of membrane type, indicating the MM could discriminate between different lipid compositions.

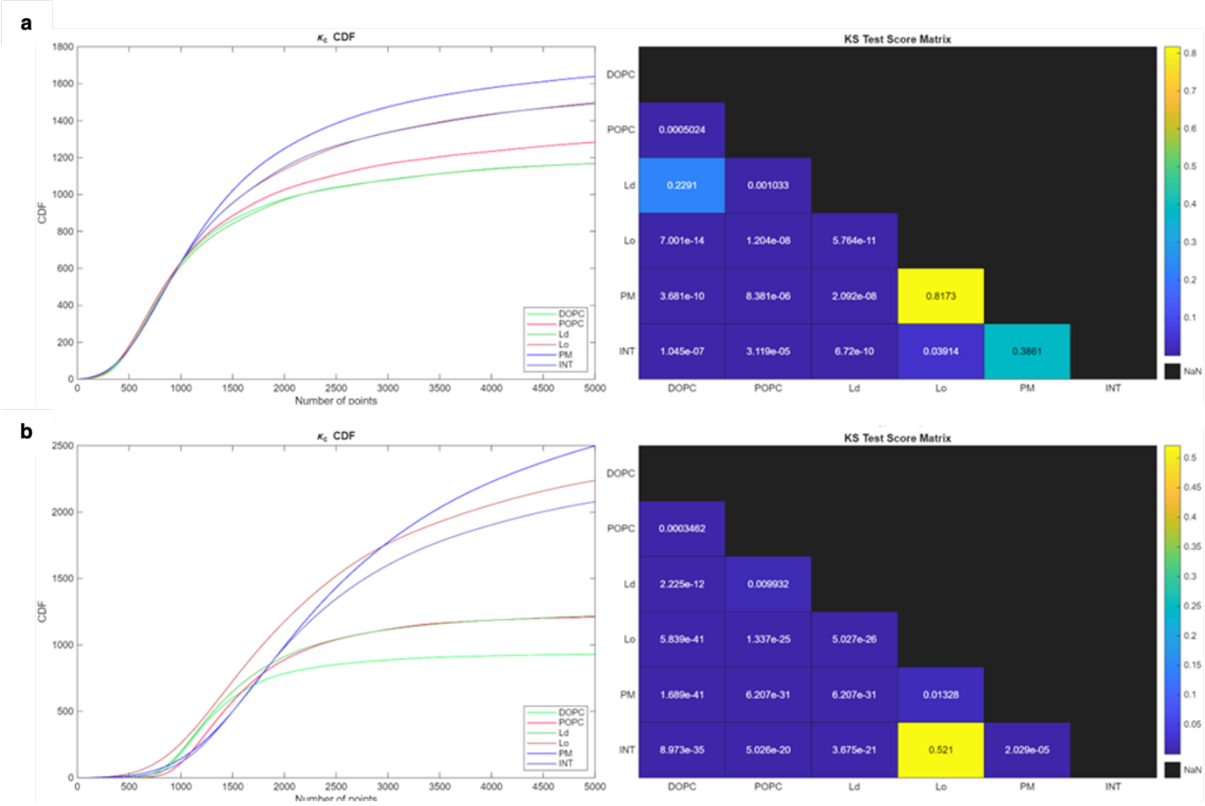

**Supplementary Figure 49. Analysis of differences in $\kappa_c$ distributions according to membrane type.** Plots show the cumulative density function (CDF) of the distributions in Supplementary Fig. 47, and the associated KS p-value. **a)** When considering fluctuations of the DNA structure, and **b)** assuming the DNA origami to be a rigid body.

4.2. Resolution

To estimate the resolution – the minimum difference in $\kappa_c$ that can be detected – the empirical $\kappa_c$ distributions (Supplementary Fig. 50) corresponding to the six experimental conditions (DOPC, POPC, Ld, Lo, PM and INT membranes) were systematically shifted. Plotting the KS test asymptotic p-values against the curve shift ($\Delta\kappa_c$) showed an inflexion point consistent across experimental conditions where p<0.05. This value was taken as the resolution limit of the MM, $\Delta\kappa_c \sim 3\text{x}10^{-21}$ J.

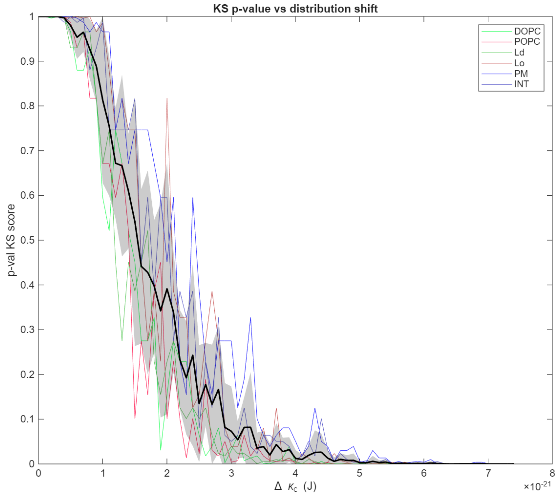

**Supplementary Figure 50. Estimation of MM resolution when measuring differences in $\kappa_c$.**

4.3. Dynamic range

To investigate the lower and upper limits of $\kappa_c$ values that could be reported, we fixed the $\tau_{Cy3}$ and $\tau_{closed}$ distributions and systematically shifted the $\tau_{open}$ distributions between 3.4 and 2.7 ns ($\tau_{Cy3}$ and $\tau_{closed}$, respectively). KS test asymptotic p-values < 0.05 against both reference distributions defined the dynamic range ($\kappa_c \sim 3 \text{ x } 10^{-20}$; $0.8 \text{ x } 10^{-20}$ J) (Supplementary Fig. 51).

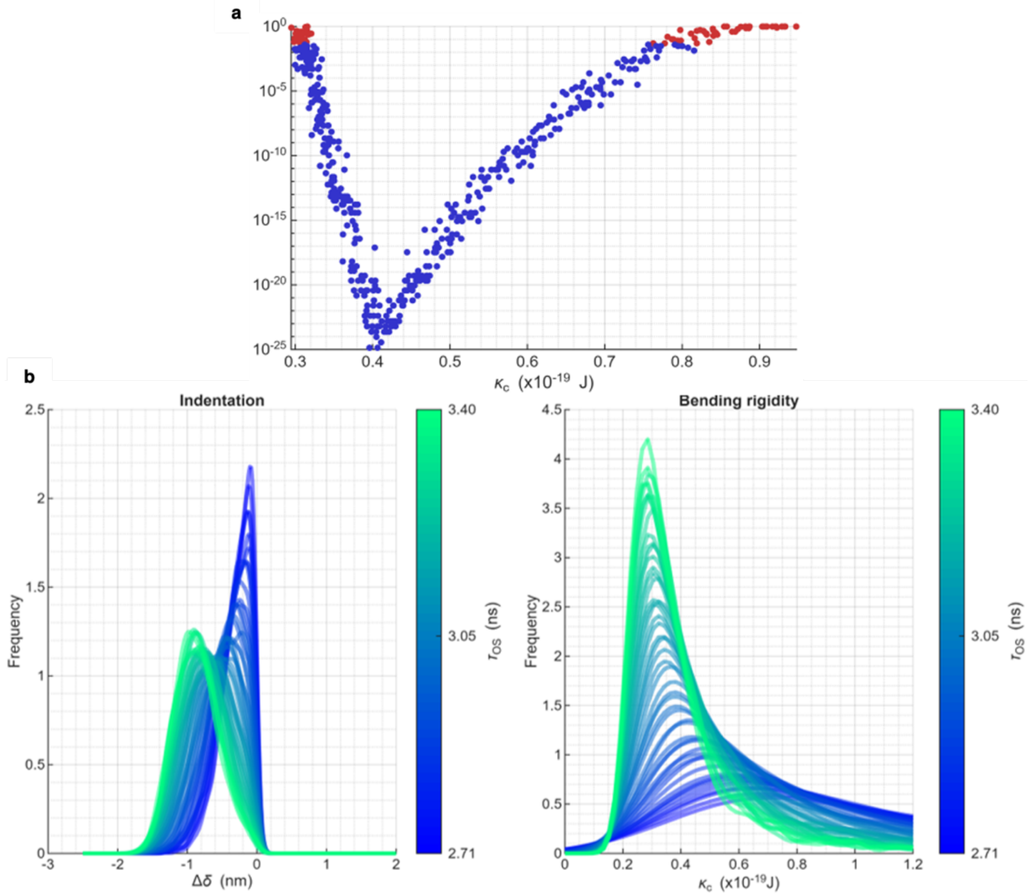

**Supplementary Figure 51. Estimation of the MM dynamic range. a)** Bootstrap distributions of bending rigidities for different values of $\tau_{open}$. Lower $\tau_{open}$ values (higher FRET) indicated more rigid membranes. **b)** Scatter plot of $\kappa_c$, taken as the peak position of the distributions in a), against the maximum asymptotic p-values obtained by comparing the trace against $\tau_{Cy3}$ and $\tau_{closed}$ distributions. p < 0.05 occurred at $\kappa_c \sim 0.3 \text{ x } 10^{-19}$; $\kappa_c \sim 0.5 \text{ x } 10^{-19}$ J.

**Supplementary Note 5: Analysis of FlipTR® probe response to membrane deformation by MM**

Actuation of MM is expected to deform the lipid bilayer. Deformation was measured by membrane-tension probe FliptR to explore whether the degree of deformation correlated with the opening angle and membrane indentation as quantified by FLIM analysis of MM signalling. FliptR reports on changes in membrane tension by its fluorescence lifetime (Supplementary Fig. 52a).[17–19] The twist angle between the two FliptR dithienothiophene (DTT) groups is affected by the membrane's deformation, so that a higher degree of lipid packing within a deformed membrane should lead to an increased lifetime of the dye. We recorded fluorescence emission decay curves of FliptR under four conditions: (1) incubation with GUVs (Ctrl), followed by sequential addition (2) of cholesterol-tagged MM (MM), (3) opening strands (+ OS), and (4) closing strands (+ CS). The decay curves were fitted with double exponentials to obtain two lifetimes ($\tau_1$and $\tau_2$) (Supplementary Table 13). The FLIM-fit images and bar charts (Supplementary Figs. 52-54) correspond to the longer lifetime ($\tau_2$) with greater number of emission events and directly reports on membrane tension.[19]

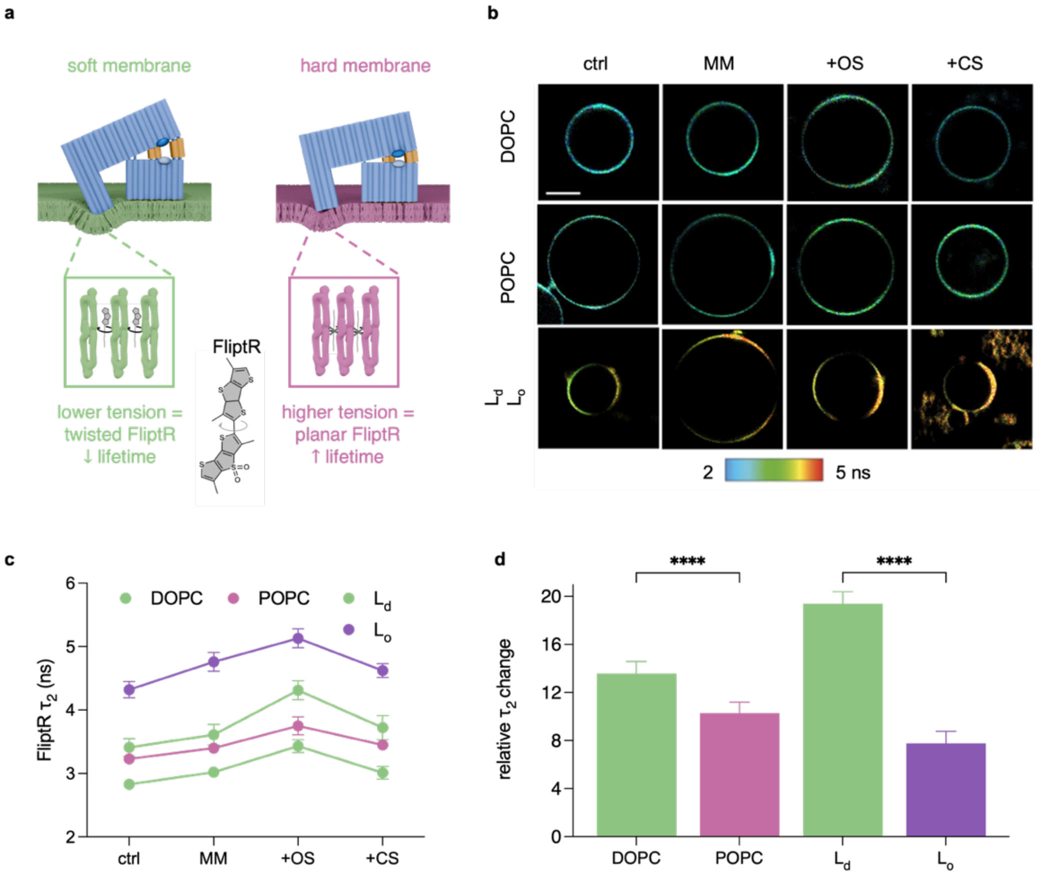

**Supplementary Figure 52. MM deformation in lipid bilayers is confirmed by FliptR. a)** Schematic diagram of FliptR lifetime shifting from green to red as order in the membrane increases. Figure adapted from *Colom et al.*[19] **b)** FLIM-fit images of FliptR within GUV membranes (Ctrl) and upon the addition of MM, opening strands (OS), and closing strands (CS). The images are colour-coded for $\tau_2$ . For reasons of clarity, the microscopy images of biphasic GUVs are arranged to show the DOPC domains on the left side of the GUV, and DPPC/CL on the right side. Scale bar, 20 µm. **c)** Line plot showing the response of FliptR $\tau_2$ to above conditions for MM bound to GUVs. The values represent the average ± SD for n > 10 GUVs per condition, across three independent experiments. Separate plots with the calculated statistical significance for each lifetime change upon addition are shown in Supplementary Figure 43. **d)** Bar chart on the calculated relative changes in $\tau_2$ (%) following addition of OS. One-way ANOVA, Tukey's multiple comparison test, ****P <0.0001; n > 10.

Lipid composition profoundly affected FliptR lifetime (Supplementary Figs. 52, 53, Supplementary Table 13) and inferred membrane tension. In the absence of DNA (Ctrl), DOPC membranes had the lowest membrane tension (2.83 ± 0.06 ns) while POPC bilayers featured higher tension (3.23 ± 0.04 ns). In the case of phase-separated GUVs, FliptR lifetimes in the DOPC-rich $L_d$ domains and DPPC-CL rich $L_o$ domains were 3.4 ± 0.14 ns and 4.32 ± 0.13 ns, respectively, in agreement with existing reports[19] with slight deviations attributed to buffer conditions. The increased lifetime in the $L_d$ region compared to pure DOPC GUVs could be due to a small proportion of cholesterol residing in the DOPC domain, as also suggested in the FRET-FLIM dataset (Supplementary Figure 52).

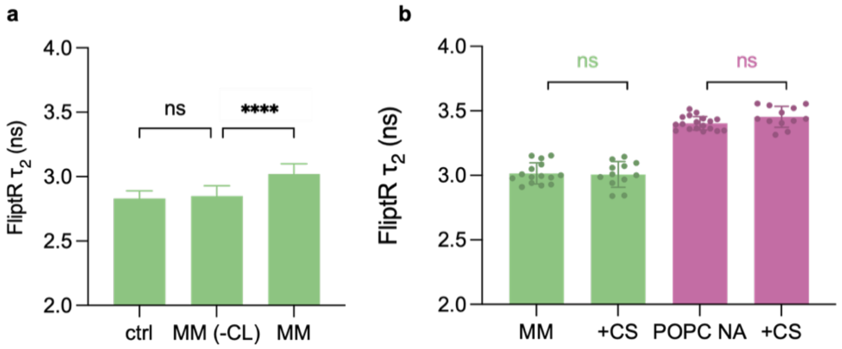

**Supplementary Figure 53. FliptR fluorescence lifetime in GUVs for control experiments using MM**. **a)** Bar chart of mean $\tau_2$ ± SD upon the addition of MM without any cholesterol tags (MM (-CL)) (n = 10) to GUVs composed of DOPC. One-way ANOVA, Tukey's multiple comparison test, *****P < 0.0001; ns = not significant. **b)** Bar chart showing mean $\tau_2$ ± SD upon the addition of CS to MM in a closed conformation (MM (closed)) to GUVs composed of DOPC and POPC. Ordinary one-way ANOVA, Tukey's multiple comparison test, ns = not significant.

**Supplementary Table 13. Lifetime components ($\tau$) extracted from FliptR analysis of MM binding to membrane vesicles**. The components are extracted from double-fitted photon lifetime histograms.

| *Lipid* | $\tau_1$ (mean ± SD) | $\tau_2$ (mean ± SD) |
|---|---|---|
| DOPC | 0.5 ± 0.1 | 2.8 ± 0.1 |
| DOPC MM | 0.6 ± 0.1 | 3.0 ± 0.1 |
| DOPC MM+OS | 0.6 ± 0.1 | 3.4 ± 0.1 |
| DOPC MM+CS | 0.6 ± 0.1 | 3.0 ± 0.1 |
| POPC | 0.6 ± 0.1 | 3.2 ± 0.1 |
| POPC MM | 0.7 ± 0.1 | 3.4 ± 0.1 |
| POPC MM+OS | 0.7 ± 0.2 | 3.7 ± 0.1 |
| POPC MM+CS | 0.6 ± 0.1 | 3.4 ± 0.1 |
| *Phase-separated vesicles* | | |
| $L_d$ | 0.5 ± 0.1 | 3.4 ± 0.1 |
| $L_d$ MM | 0.5 ± 0.1 | 3.6 ± 0.2 |
| $L_d$ MM+OS | 0.7 ± 0.2 | 4.3 ± 0.1 |
| $L_d$ MM+CS | 0.5 ±0.1 | 3.7 ± 0.2 |
| $L_o$ | 0.6 ± 0.2 | 4.3 ± 0.1 |
| $L_o$ MM | 0.7 ± 0,2 | 4.8 ± 0.1 |
| $L_o$ MM+OS | 0.8 ± 0.2 | 5.1 ± 0.1 |
| $L_o$ MM+CS | 0.7 ± 0.1 | 4.6 ± 0.1 |

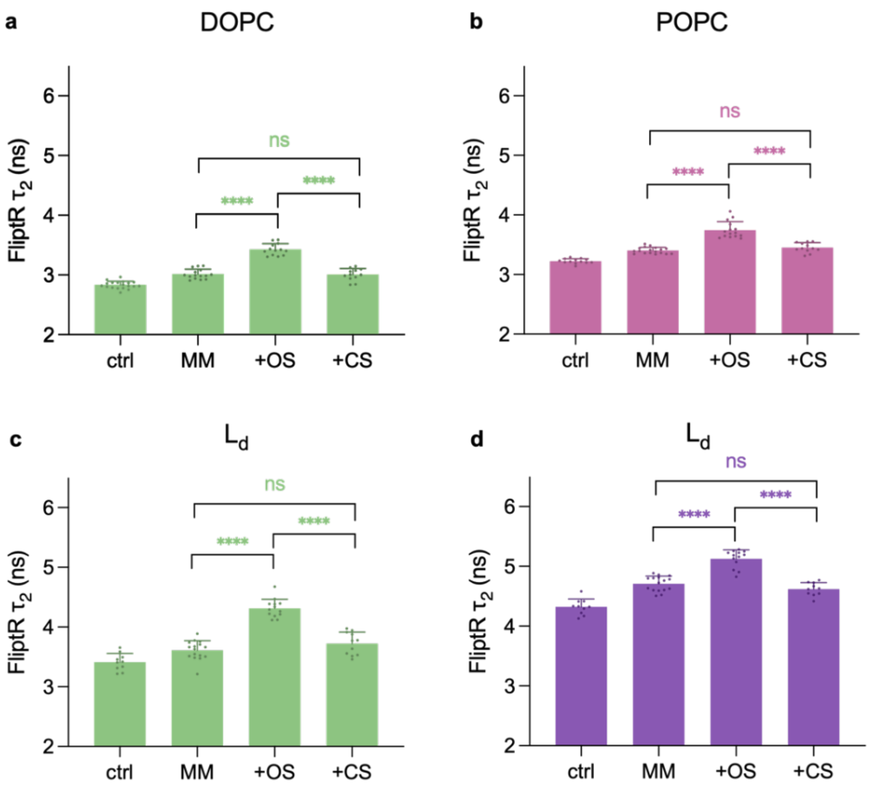

**Supplementary Figure 54. Response of FliptR fluorescence lifetime upon addition of MM to GUVs of different membrane composition and elasticities. a)** Bar charts show mean $\tau_2$ ± SD in GUVs composed of DOPC (n = 13) and upon addition of MM (n = 15), OS (n = 13), CS (n = 12). **b)** Bar charts show mean $\tau_2$ ± SD in GUVs composed of POPC (n = 13) and upon addition of MM (n = 18), OS (n = 15), CS (n = 12). **c)** Bar charts show mean $\tau_2$ ± SD in GUVs composed of $L_d$ and **d)** $L_o$. (n = 10) and upon addition of MM (n = 18), OS (n = 13), CS (n = 11). a-c) One-way ANOVA, Tukey's multiple comparison test, *****P < 0.0001; ns = not significant.

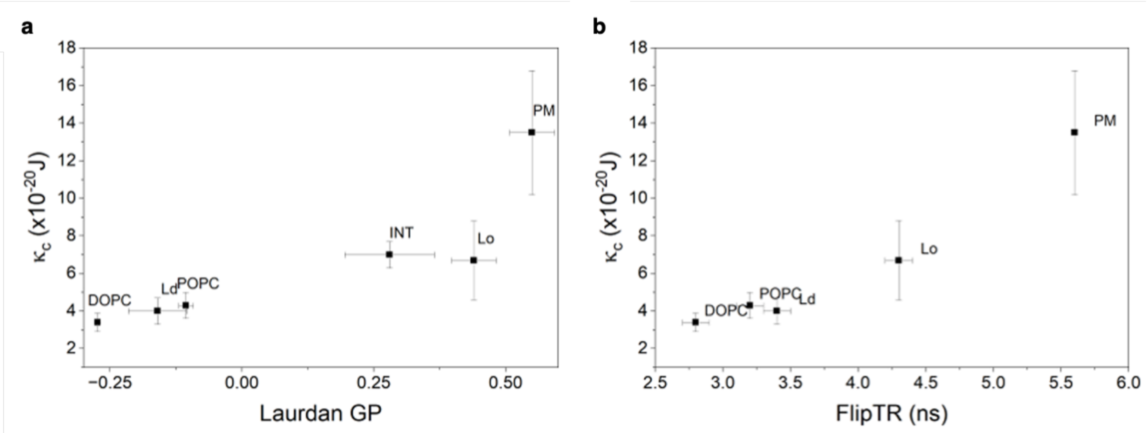

**Supplementary Figure 55. Non-linear relationship between membrane elasticity and lipid packing.** Lipid packing, as measured by a, Laurdan's general polarization (GP), or b, FlipTR lifetime does not scale linearly with bending rigidity. The FlipTR data point for INT membrane compartments is not available as the FlipTR dye does not cross the plasma membrane to reach internal membranes. The FlipTR value for PM is from the literature and lacks the corresponding error bar.

After experimentally proving by the FlipTR lifetime changes the different bilayer deformations caused by MM (Supplementary Fig. 54), we explored whether the measured changes could be, indeed, explained by the nanoscale membrane deformations in the vicinity of the DNA nanostructure's tip. If this were to be the case, then the calibration factor ξ describing FlipTR lifetime change response to membrane deformation would for our MM-mediated indentation be within the range found in literature. To this end, we first defined the relative FlipTR lifetime $\chi$ between DOPC and POPC GUVs after MM addition as:

$$\chi := \frac{\tau_{DOPC}}{\tau_{POPC}} - 1$$

*Equation (5.1)*

where the lifetime after MM actuation can be described as:

$$\tau_x = \tau_{x,0} + \Delta\tau_x$$

*Equation (5.2)*

with $\tau_{x,0}$ being the lifetime after the MM is attached to the bilayer, but before the opening strands are added, and $\Delta\tau_x$ being the change in lifetime coming from MM action and membrane indentation and resulting membrane tension.

$\Delta\tau_x$ may then be related to the membrane area strain $u_x$ as:

$$\Delta\tau_x := \xi_x K_{A,x} u_x = \Lambda_x u_x$$

*Equation (5.3)*

where $K_{A,x}$ represents the area stretching modulus, $u_x$ represents the area strain and $\xi_x$ represents a calibration constant accounting for the tension/lifetime dependence of the FliptR probe. Therefore, the measured lifetime ratio after MM opening can be expressed as:

$$\chi = \frac{\tau_{DOPC,0} + \Lambda_{DOPC} u_{DOPC}}{\tau_{POPC,0} + \Lambda_{POPC} u_{POPC}} - 1$$

*Equation (5.4)*

This can be expanded as:

$$\chi = \frac{\tau_{DOPC,0}\tau_{POPC,0}}{\tau_{POPC,0}^2} + \frac{\tau_{POPC,0}\Lambda_{DOPC} u_{DOPC}}{\tau_{POPC,0}^2} + \frac{\tau_{DOPC,0}\Lambda_{POPC} u_{POPC}}{\tau_{POPC,0}^2} - 1$$

*Equation (5.5)*

Taking $\tau_{DOPC,0} = 2.26\, ns$ and $\tau_{DOPC,0} = 2.29\, ns$, $\chi$ can be rewritten as:

$$\chi \approx \frac{\tau_{DOPC,0}\Lambda_{POPC} u_{POPC} + \tau_{POPC,0}\Lambda_{DOPC} u_{DOPC}}{\tau_{POPC,0}^2}$$

*Equation (5.6)*

Given that both membranes are in the same phase and do not exhibit phase separation, it is reasonable to assume the calibration factor ξ will be similar for both DOPC and POPC bilayers, and therefore:

$$\xi \approx \frac{\chi\, \tau_{POPC,0}^2}{\tau_{POPC,0}\alpha_{DOPC} K_{DOPC} u_{DOPC} + \tau_{DOPC,0}\alpha_{POPC} K_{POPC} u_{POPC}}$$

*Equation (5.7)*

Assuming membrane coverage and area extension moduli will be the similar in both cases (i.e. $\alpha_{DOPC}$ = $\alpha_{POPC}$ and $K_{DOPC}$ = $K_{POPC}$) allows to express the calibration factor $\xi$ as:

$$\xi \approx \frac{1}{\alpha} \frac{\chi\, \tau_{POPC,0}^2}{K(\tau_{POPC,0} u_{DOPC} + \tau_{DOPC,0} u_{POPC})}$$

*Equation (5.8)*

On the other hand, one also needs to consider which percentage of GUV membrane surface experiences MM-mediated deformation. Towards this end, membrane area strain *u* can be estimated as the transition from a planar membrane patch of radius *a* to that of a spherical cap whose height is determined by the indentation depth $\delta$ (Supplementary Fig. 55). By taking the values of $\delta$ derived previously for DOPC and POPC membranes and using *a* = 9.8 nm (based on the MM tip geometry, Supplementary Fig. 50) it is possible to obtain the area of the spherical cap after deformation ($A_\delta$) which yields $A_{\delta,DOPC}$ = 328 nm$^2$ and $A_{\delta,POPC}$ = 320 nm$^2$ respectively, which would correspond to an area strain $u_{DOPC}$ = 0.085 and $u_{POPC}$ = 0.062, respectively. We note that these values are around the limit for bilayer breaking (~8%), although they are a limiting case, as we reckon the height of the cap base will be larger, and consequently the area strain will be reduced.

Taking *K* ~ 250 mN/m, it is possible to calculate the FliptR calibration factor as a function of membrane coverage, as shown in Supplementary Fig. 55. We can then define GUV coverage as the ratio between the total "indentation area" and the accessible bilayer surface so that:

$$\alpha = \frac{A_\delta}{A_{GUV}} = \frac{V_{MM} C_{MM} a_\delta}{V_{GUV} a_{lipid}/2}$$

*Equation (5.9)*

Given $V_{MM}$ = 15 µL, $C_{MM}$ = 50 nM, $V_{GUV}$ = 15 µL, $C_{GUV}$ ~ 2 mM (see Methods section) and considering the area per lipid to be 72.4 Å$^2$ and 69.3 Å$^2$ for DOPC and POPC, respectively, with an indentation area of 361 nm$^2$ per MM, it is possible to estimate under the assumption of complete binding of MM to GUVs a vesicle coverage of $\alpha_{DOPC}$ ~ 2.5 % and $\alpha_{POPC}$ ~ 2.6 % respectively. These coverage values correspond to a calibration value of ~-1.36 ns m/mN, within the range of those reported by *Colom et al.* Therefore, we conclude that the observed changes in FliptR lifetime are compatible with an MM-induced membrane deformation.

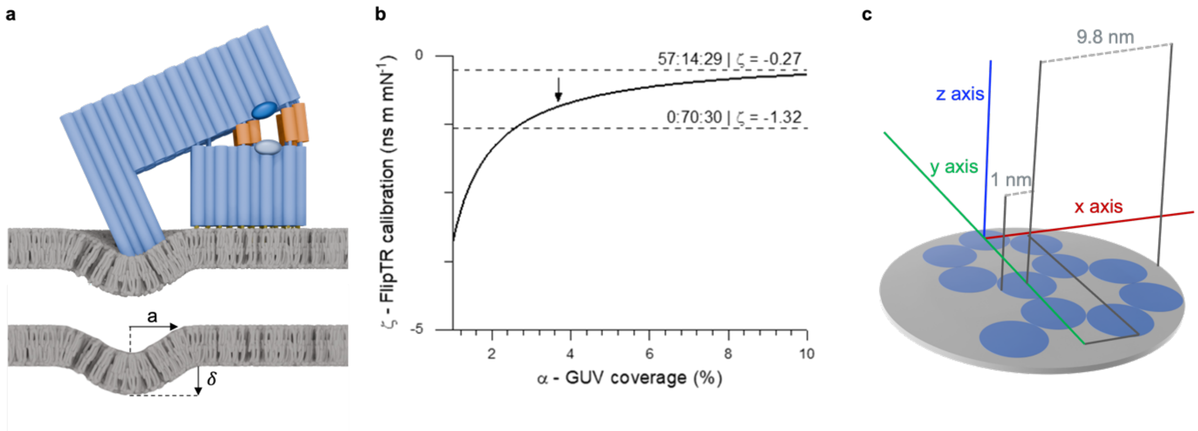

**Supplementary Figure 56. FliptR-based calibration. a)** Model employed to estimate the FlipTR calibration factor ξ, showing the "spherical cap"-like deformation induced by the MM. **b)** Plot showing how $\xi$ depends on the estimated membrane coverage. The black arrow indicates the estimated coverage for our system. The corresponding FlipTR calibration value lies between the ones reported by *Colom et al.*[19] **c)** Schematic illustration of the footprint of the MM tip which touches the membrane. The blue discs indicate the DNA duplexes of the tip. The grey discs describe the size and radius of the tip model used for the calibration calculations.

**Supplementary Note 6: Biophysical membrane properties and their limited effect on MM-based readout**

6.1. Thermal membrane fluctuations are of smaller amplitude than MM-induced membrane indentation

Calculations show that the amplitude of the membrane fluctuations has a vertical height amplitude which is considerably smaller than the membrane indentation depths obtained with MM. For these calculations, we use Helfrich's curvature elasticity model for lipid membranes, which calculates the energy cost of bending lipid bilayers.[20] Applying Helferich's model shows that the membrane fluctuations have a vertical height amplitude of ~0.1 nm at the MM tip's length scale. This is considerably smaller than the membrane indentation depths obtained with MM, ranging from 0.4 to 1.0 nm.

In more detail, Helfrich's model predicts the thermally induced vertical height fluctuation $h_q$ for a set frequency mode q as:

$$\langle |h_q|^2 \rangle = \frac{k_B T}{L^2[\kappa_c q^4 + \sigma q^2]}$$

given a membrane patch of dimensions L x L, and with membrane rigidity $\kappa_c$ and tension $\sigma$.

Assuming a vanishing tension, the average across modes, referring to the mean-square amplitude of thermal fluctuations, are:

$$\langle h^2 \rangle = \sum_q \langle |h_q|^2 \rangle = \cdots = \frac{L^2 k_B T}{16\pi^3 \kappa_c}$$

For fluctuations compared to the MM-induced deformation (i.e. ~1 nm), we obtain:

$$L \sim \sqrt{\frac{16\pi^3 \kappa_c \langle h^2 \rangle}{k_B T}} \xrightarrow[\kappa_c = 10^{-19} J]{h = 1nm} L \sim 10^{-7} m = 100\, nm$$

This sets a lower (~100 nm) length scale limit for the membrane's fluctuations. When scaling this to MM's smaller tip dimensions with $L \sim 10\, nm$ diameter, we obtain a considerably smaller vertical height amplitude of ~0.1 nm

$$\Delta h = \sqrt{\langle h^2 \rangle} = \sqrt{\frac{L^2 k_B T}{16\pi^3 \kappa_c}} \xrightarrow[\kappa_c = 10^{-19} J]{L = 10nm} \Delta h \sim 10^{-10} m = 0.1\, nm$$

6.2. Thermal membrane fluctuations are considerably faster than MM-induced membrane deformations

Thermal membrane fluctuations are at the time scale of microseconds[21] are much faster than the min-long time-course of the MM-based measurements. Thus, the changes seen at our experimental timescales are driven by the MM.

The minute-scale kinetics for the FRET signal are not linked to membrane-molecular processes. Rather, they arise from a combination of the hybridisation kinetics of the opening strands to the complementary DNA strands and hairpins[22], steric effects delaying the opening strands from accessing the hairpins situated between the two DNA origami blocks (cantilever and base), and the diffusion of the OS themselves. Similar minute-scale kinetics have also been observed for other hybridisation-actuated DNA constructs[23]. As the minute-scale timescales of recording the fluorescent signals are also much longer than the µs thermal fluctuations[21], they average and cancel out thermal effects.

It is also noted that close interactions of nanostructures with lipid membranes usually lower their thermal membrane fluctuations. For example, α-synuclein interaction reduces thermal membrane fluctuations as shown with NMR, biophysical, and computational methods[24]. A similar effect on lipids is observed when cytoskeletal proteins or biomolecular condensates closely interact with membranes[25,26].

### 6.3. MM action induces membrane bending but not membrane thinning, and bending rigidity dominates over membrane tension

Lipid membranes can deform in two ways along the bilayer's normal direction: By bending or by thinning or thickening. For free-standing membranes, without an underlying solid support, bending is preferred over thinning, as explained below. Furthermore, the extent of membrane deformations is controlled by the bending rigidity and the membrane's tension, the 'pulling' force per unit length controlling of how tightly packed lipids are. Bending rigidity dominates in MM-induced membrane deformation, given the small-scale deformations as demonstrated below.

*Thinning vs bending:*

For our free-floating membrane system, the energy cost of compression is significantly higher (~x$10^3$) than that of membrane bending, as shown in the following calculations below. This is also supported by the small nN-scale forces required for bending liposomal membranes compared to the larger nN forces for compressing SLBs[27].

The energetic cost for membrane bending is derived for a given membrane area and radius of curvature Rc using:

$$u_{bend} = \frac{1}{2R_c^2}\kappa_c = \frac{1}{2R_c^2}\frac{1}{48}K_{stretch}h^2 = \frac{1}{2R_c^2}\frac{1}{48}\Upsilon h_0^3$$

where $K_{stretch}$ represents the stretching modulus and $\Upsilon$ the elastic Young's modulus. The bending energy $u_{bend}$ per unit area is for two uncoupled elastic sheets (e.g. non-registered or interdigitated leaflets).

By contrast, the energy cost per unit area for membrane compression or thinning, assuming small deformations, is:

$$u_{thinning} = \frac{1}{2}\Upsilon\frac{(h-h_0)^2}{h_0}$$

Therefore:

$$\frac{u_{bend}}{u_{thinning}} = \frac{h_0^4}{48R_c^2(\mathrm{h}-\mathrm{h}_0)^2}$$

The radius of curvature can be estimated from a spherical cap of height $h_c$~1 nm and base radius a~10 nm (Supplementary Note 5), so that:

$$R_c = \frac{h_c^2 + a^2}{2h_c} \sim 50\,nm$$

Hence,

$$\frac{u_{bend}}{u_{thinning}} \approx \frac{4nm^4}{48(50nm)^2(1nm)^2} \approx 2\mathrm{x}10^{-3}$$

Indicating that bending is more likely to occur than membrane compression or thinning.

*Tension vs bending rigidity for bending deformations:*

The contribution of membrane tension is limited due to the small deformation length scale (~10 nm) compared to the fluctuation wavelength of around 200 nm, which is calculated in the following. According to Helfrich theory, thermal membrane fluctuation energy for amplitude $h_q$ in mode **q** is:

$$U_{bend} = L^2 \sum_q \left|h_q\right|^2 \frac{1}{2}[\kappa_c q^4 + \sigma q^2]$$

From which we can find a cross-over length

$$\kappa_c q^4 = \sigma q^2 \rightarrow q_c = \sqrt{\frac{\sigma}{\kappa_c}}$$

Considering typical values $\kappa_c \sim 10^{-19} J$, $\sigma \sim 0.1\ \mathrm{mN/m}$[28], then $q_c \sim 3\mathrm{x}10^7$, which corresponds to a fluctuation wavelength of:

$$\lambda_c = \frac{2\pi}{q_c} \sim 2\mathrm{x}10^{-7}\mathrm{m} = 200\mathrm{nm}$$

Below this length (which is the MM case), increase in q implies that bending will dominate the membrane's response.

**Bibliography:**

1. Roy, R., Hohng, S. & Ha, T. A practical guide to single-molecule FRET. *Nat. Methods* **5**, 507–516 (2008).
2. Acerbi, I. *et al.* Integrin-specific mechanoresponses to compression and extension probed by cylindrical flat-ended AFM tips in lung cells. *PLOS ONE* **7**, e32261 (2012).
3. Jadidi, T., Seyyed-Allaei, H., Tabar, M. R. R. & Mashaghi, A. Poisson's ratio and Young's modulus of lipid bilayers in different phases. *Front. Bioeng. Biotechnol.* **2**, 1–6 (2014).
4. Weaver, J. C. & Chizmadzhev, Y. A. Theory of electroporation : A review. *Biochem. Bioenerg.* **41**, 135–160 (1996).
5. Wohlert, J., den Otter, W. K., Edholm, O. & Briels, W. J. Free energy of a trans-membrane pore calculated from atomistic molecular dynamics simulations. *J. Chem. Phys.* **124**, 154905 (2006).
6. Thompson, A. J. *et al.* Molecular rotors provide insights into microscopic structural changes during protein aggregation. *J. Phys. Chem. B* **119**, 10170–10179 (2015).
7. Paez-Perez, M. & Kuimova, M. K. Molecular rotors: Fluorescent sensors for microviscosity and conformation of biomolecules. *Angew. Chem. Int. Ed.* **63**, e202311233 (2024).
8. Rawicz, W., Olbrich, K. C., McIntosh, T., Needham, D. & Evans, E. Effect of chain length and unsaturation on elasticity of lipid bilayers. *Biophys. J.* **79**, 328–339 (2000).
9. Elani, Y. *et al.* Measurements of the effect of membrane asymmetry on the mechanical properties of lipid bilayers. *Chem. Commun.* **51**, 6976–6979 (2015).
10. Doktorova, M., Harries, D. & Khelashvili, G. Determination of bending rigidity and tilt modulus of lipid membranes from real-space fluctuation analysis of molecular dynamics simulations. *Phys. Chem. Chem. Phys.* **19**, 16806–16818 (2017).
11. Niggemann, G., Kummrow, M. & Helfrich, W. The bending rigidity of phosphatidylcholine bilayers: Dependences on experimental method, sample cell sealing and temperature. *J. Phys. II* **5**, 413–425 (1995).
12. Drabik, D., Przybyło, M., Chodaczek, G., Iglič, A. & Langner, M. The modified fluorescence based vesicle fluctuation spectroscopy technique for determination of lipid bilayer bending properties. *Biochim. Biophys. Acta BBA - Biomembr.* **1858**, 244–252 (2016).
13. Nagle, J. F. Experimentally determined tilt and bending moduli of single-component lipid bilayers. *Chem. Phys. Lipids* **205**, 18–24 (2017).

14. Venable, R. M., Brown, F. L. H. & Pastor, R. W. Mechanical properties of lipid bilayers from molecular dynamics simulation. *Chem. Phys. Lipids* **192**, 60–74 (2015).
15. Stetter, F. W. S., Cwiklik, L., Jungwirth, P. & Hugel, T. Single Lipid Extraction: The Anchoring Strength of Cholesterol in Liquid-Ordered and Liquid-Disordered Phases. *Biophys. J.* **107**, 1167–1175 (2014).
16. Ho, D. *et al.* Force-Driven Separation of Short Double-Stranded DNA. *Biophys. J.* **97**, 3158–3167 (2009).
17. Dal Molin, M. *et al.* Fluorescent flippers for mechanosensitive membrane probes. *J. Am. Chem. Soc.* **137**, 568–571 (2015).
18. Goujon, A. *et al.* Mechanosensitive fluorescent probes to image membrane tension in mitochondria, endoplasmic reticulum, and lysosomes. *J. Am. Chem. Soc.* **141**, 3380–3384 (2019).
19. Colom, A. *et al.* A fluorescent membrane tension probe. *Nat. Chem.* **10**, 1118–1125 (2018).
20. Helfrich, W. Elastic properties of lipid bilayers: theory and possible experiments. *Z. Naturforschung Teil C Biochem. Biophys. Biol. Virol.* **28**, 693–703 (1973).
21. Rodríguez-García, R. *et al.* Bimodal Spectrum for the Curvature Fluctuations of Bilayer Vesicles: Pure Bending plus Hybrid Curvature-Dilation Modes. *Phys. Rev. Lett.* **102**, 128101 (2009).
22. Zhang, J. X. *et al.* Predicting DNA hybridization kinetics from sequence. *Nat. Chem.* **10**, 91–98 (2018).
23. Burns, J. R., Seifert, A., Fertig, N. & Howorka, S. A biomimetic DNA-based channel for the ligand-controlled transport of charged molecular cargo across a biological membrane. *Nat. Nanotechnol.* **11**, 152–156 (2016).
24. Viennet, T. *et al.* Structural insights from lipid-bilayer nanodiscs link α-Synuclein membrane-binding modes to amyloid fibril formation. *Commun. Biol.* **1**, 44 (2018).
25. Ioannou, I. A., Brooks, N. J., Marina K. Kuimova & Elani, Y. Visualizing Actin Packing and the Effects of Actin Attachment on Lipid Membrane Viscosity Using Molecular Rotors. *JACS Au* **4**, 2041–2049 (2024).
26. Mangiarotti, A. *et al.* Biomolecular condensates modulate membrane lipid packing and hydration. *Nat. Commun.* **14**, 6081 (2023).

27. Et-Thakafy, O. *et al.* Mechanical Properties of Membranes Composed of Gel-Phase or Fluid-Phase Phospholipids Probed on Liposomes by Atomic Force Spectroscopy. *Langmuir* **33**, 5117–5126 (2017).
28. Kozlov, M. M. & Chernomordik, L. V. Membrane tension and membrane fusion. *Curr. Opin. Struct. Biol.* **33**, 61–67 (2015).
